# Maturation-dependent complement production and C3 processing in human retinal pigment epithelium cells

**DOI:** 10.1101/2025.02.05.636432

**Authors:** Juliane Schikora, Antonia Nickel, Jasmin Bergert, Rike Hähnel, Aaron Dort, Stella Y. Schayan-Araghi, Pratiti Banerjee, Hannah N. Wolf, Helen May-Simera, Diana Pauly

## Abstract

Induced pluripotent stem cell-derived retinal pigment epithelial (iPSC-RPE) cells, closely resembling healthy RPE, offer valuable insights for retinal disease modelling. This study evaluates immature and mature phenotypes of iPSC-RPE and ARPE-19 cells, comparing RPE characteristics, cell-associated complement profiles, and TGF-β1-mediated stress responses across transcript expression, secretion, and protein levels. Statistical analyses were performed using independent means *t*-test and Wilcoxon rank-sum test. Mature iPSC-RPE cells exhibited characteristic RPE-morphology, with elevated secretion of complement components (C3, FH/FHL-1, FI), unique FB secretion, and apical complement localisation. Intracellular C3 processing revealed cleavage products different from known blood-derived C3 fragments and showed maturation-dependent differences; only mature iPSC-RPE cells secreted active C3 forms (C3(H_2_O), C3a) and exhibited the intact C3 β-chain. Mature ARPE-19 and iPSC-RPE cells demonstrated resistance to TGF-β1 treatment, which reduced complement secretion without affecting C3a release. In conclusion, ARPE-19 and iPSC-RPE cells demonstrated local production of complement components and maturation-dependent intracellular processing of C3 into active forms. These findings highlight the impact of RPE maturation on local C3 activation, providing a basis for future studies on C3 functionality in the RPE and its potential pathological effects.

**Impact statement:** Cell state-dependent local processing of complement C3 in retinal pigment epithelium cells generates active C3 forms that may evade anti-C3 therapies, influencing age-related macular degeneration and treatment outcomes.

**Funding statement:** This project has received funding from the PRO RETINA, Amberg, Germany, under grant agreement number “Pro-Re/Projekt/Schikora – Pauly.05-2022”, and Deutsche Forschungsgemeinschaft (DFG), under grant agreement number “498244102 (MA 6139/5-1)“. Open access funding provided by the Open Access Publishing Fund of University Marburg. Funding sources were not involved in study design, data collection and interpretation, or the decision to submit the work for publication.

**Graphical abstract:** 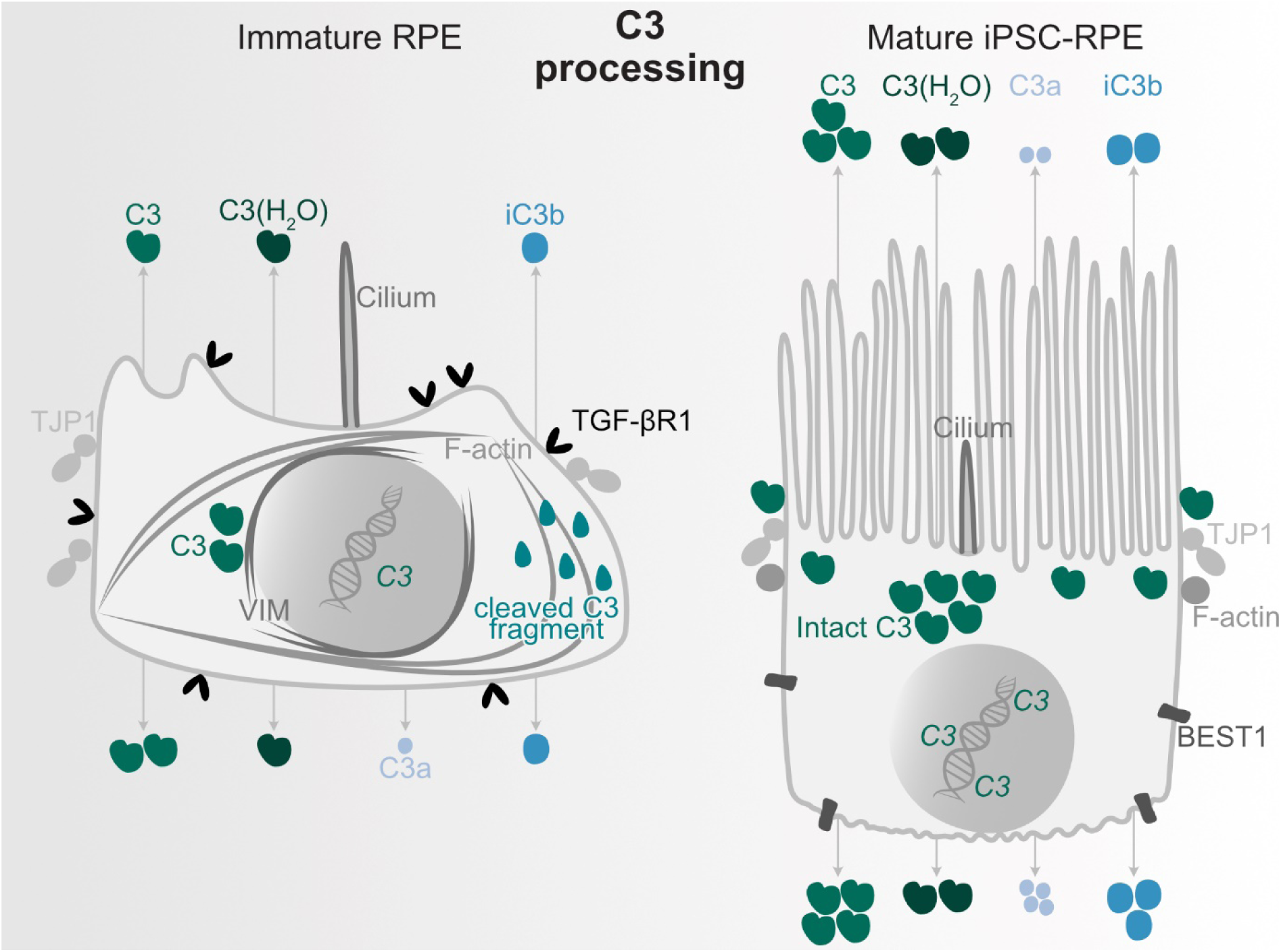

## 1. Introduction

Age-related macular degeneration (AMD) is a leading cause of vision loss among the elderly, driven by multiple risk factors, including ageing, smoking, and genetic variations (Flaxman et al., 2017; Heesterbeek et al., 2020; Seddon, 2005; Wong et al., 2014). Clinically, late AMD is categorised into two main subtypes: the exudative and non-exudative forms (Ferris et al., 2013). Therapeutic options for the non-exudative form remain limited, particularly in advanced stages that involve geographic atrophy (GA) developments. GA is characterised by degeneration of the retinal pigment epithelium (RPE) and photoreceptors, ultimately resulting in progressive vision loss and blindness (Ferris et al., 2013).

The RPE is a monolayer of specialised epithelial cells within the retina, distinguished by its hexagonal morphology and polarised organisation, which contributes to several critical functions necessary for retinal health and functionality. These include the phagocytosis of photoreceptor outer segment tips and maintenance of the blood-retina barrier (Bharti et al., 2011; Lakkaraju et al., 2020). The primary cilium of RPE cells plays a pivotal role in signalling pathways essential for RPE maturation and function (Kretschmer et al., 2023; May-Simera et al., 2018).

Several research groups have investigated hyperreflective foci (HRF) and their origin as potential indicators of AMD progression. These studies have identified morphologically different RPE cells in and near atrophic regions of the retina, which have lost their specific RPE characteristics (Balaratnasingam et al., 2017; Bonilha et al., 2020; Cao et al., 2021; Gambril et al., 2019; Zanzottera et al., 2016; Zanzottera, Messinger, Ach, Smith, Freund, et al., 2015). Epithelial-to-mesenchymal transition (EMT), triggered by AMD-associated risk factors, has been proposed as a possible mechanism contributing to RPE pathology and HRF formation (Bonilha et al., 2020; Ghosh et al., 2018). Furthermore, the activation of the transforming growth factor-beta (TGF-β) signalling pathway, a known inducer of EMT, has been implicated in the progression of AMD pathogenesis (Little, Llorián-Salvador, Tang, Du, Marry, et al., 2020; Newman et al., 2012; Radeke et al., 2015). Transforming growth factor-beta 1 (TGF-β1) treatment of RPE cells is a widely accepted model for inducing EMT, pushing RPE cells toward a mesenchymal phenotype, and impairing their functionality (S.-J. Kim et al., 2020; Lee et al., 2013; M. Li et al., 2021; Y. Li et al., 2015; Ma et al., 2023; Su et al., 2021; M. Wang et al., 2022; Wei et al., 2018). As a consequence, RPE dysfunction is a key factor in AMD development and progression, as the RPE performs a wide range of functions essential for retinal health. Another critical process implicated in AMD pathogenesis is the complement system. Genome-wide association studies have identified variants in complement-related genes, including complement 3 (*C3*), complement factor B (*CFB*), complement factor H (*CFH*), and complement factor I (*CFI*), that are associated with an increased risk of AMD (Despriet et al., 2009; Edwards et al., 2005; Fritsche et al., 2013, 2016; Hageman et al., 2005; Klein et al., 2005; Maller et al., 2007; Park et al., 2009; Yates et al., 2007). Additionally, components and regulators of the complement system, such as C3, complement 3b (C3b), inactivated complement 3b (iC3b), complement 3dg (C3dg), FH, and FI, have been detected in drusen, extracellular deposits characteristic of AMD located between the RPE and Bruch’s membrane (Anderson et al., 2002, 2010; Crabb et al., 2002; P. T. Johnson et al., 2006; L. V. Johnson et al., 2001; Laine et al., 2007). Despite the immune privilege of the eye, enforced by the blood-retina barrier that separates the retina from the liver-produced systemic complement system, recent studies have revealed the local production of complement components within the RPE/choroid complex and intracellularly in RPE cells (Anderson et al., 2010; Enzbrenner et al., 2021; Y. H. Kim et al., 2009; Luo et al., 2011; Pauly et al., 2019; Rutar et al., 2012; Schäfer et al., 2017, 2020, 2021; Sugita et al., 2018; Trakkides et al., 2019; Zauhar et al., 2022).

In recent years, the approval of complement inhibitors by the Food and Drug Administration has introduced a novel therapeutic strategy for GA. Pegcetacoplan (Syfovre™), a C3 inhibitor, prevents the cleavage of C3 into its pro-inflammatory fragments, complement 3a (C3a) and C3b, thereby reducing excessive complement activation (Liao et al., 2020). The central complement component, C3, consists of α and β chains. Spontaneous hydrolysis non-proteolytically activates C3 into its reactive form, C3(H_2_O). Additionally, C3 cleavage by C3 convertases generates C3a and C3b, with C3b subsequently cleaved by FI and its co-factors to form iC3b, which maintains inflammatory capacity, and complement 3f (C3f). iC3b is then degraded into inactive complement 3c (C3c) and C3dg, the latter serving as an immune mediator before its final cleavage into complement 3d (C3d) and complement 3g (C3g) (Merle, Church, et al., 2015; Merle, Noe, et al., 2015; Ricklin et al., 2016).

Given the potential of HRF as an indicator of AMD progression and the critical role of the complement system in AMD pathogenesis, we aimed to investigate the cellular complement system in dysfunctional RPE cells, potentially resembling those associated with HRF. We developed a model that utilised the reverse process of EMT, known as mesenchymal-to-epithelial transition (MET), incorporating both immature and mature ARPE-19 (adult retinal pigment epithelial cell line) and induced pluripotent stem cell-derived RPE (iPSC-RPE) cells to represent dysfunctional RPE and healthy RPE, respectively. We characterised general RPE cell characteristics in this model and subsequently examined the local cellular complement system, focussing specifically on C3 and its intracellular activation products. Finally, we subjected the mature ARPE-19 and iPSC-RPE cells to EMT-inducing TGF-β1 treatment to assess both RPE cell characteristics and alterations in C3 processing.

## 2. Results

### 2.1. Mature ARPE-19 and iPSC-PRE cells exhibit an epithelial phenotype and elevated RPE markers

Since HRF as indicators of AMD progression are potentially composed of transformed RPE cells, we established a model comparing immature and mature RPE cells to represent transdifferentiated dysfunctional and healthy RPE states, respectively. The differently matured ARPE-19 cells were obtained by culturing the cells for two weeks in distinct media with high and low concentrations of fetal bovine serum and additional differentiation-inducing supplements (Hazim et al., 2019). iPSC-RPE cells were cultivated for one week to represent immature cells and for six weeks to represent mature cells.

A characterisation of the cytoskeleton in immature and mature ARPE-19 and iPSC-RPE cells was performed to assess epithelial versus mesenchymal characteristics. Immature ARPE-19 cells exhibited a mesenchymal-like filamentous actin (F-actin) cytoskeleton, characterised by prominent stress fibres and elongated F-actin structures traversing the cells (**Figure 1A**). In contrast, mature ARPE-19 cells displayed a cytoskeletal organisation leaning towards an epithelial phenotype, characterised by circumferential F-actin distribution and reduced presence of stress fibres. Fluorescence intensity analysis confirmed these observations, revealing a significant reduction in F-actin intensity in mature ARPE-19 cells compared with immature cells (*P* = .017) (**Figure 1 - supplement 1A**). Both immature and mature iPSC-RPE cells showed an epithelial circumferential distribution of F-actin (**Figure 1A**); however, immature iPSC-RPE cells displayed more stress fibres compared with mature iPSC-RPE cells, which showed an apical localisation of F-actin with no observable stress fibres (**Figure 1 - video 1**). No significant differences in F-actin fluorescence intensity were observed in iPSC-RPE cells (**Figure 1 - supplement 1A**). Negative controls for all immunostainings are presented in **Appendix 1.**

**Figure 1:**
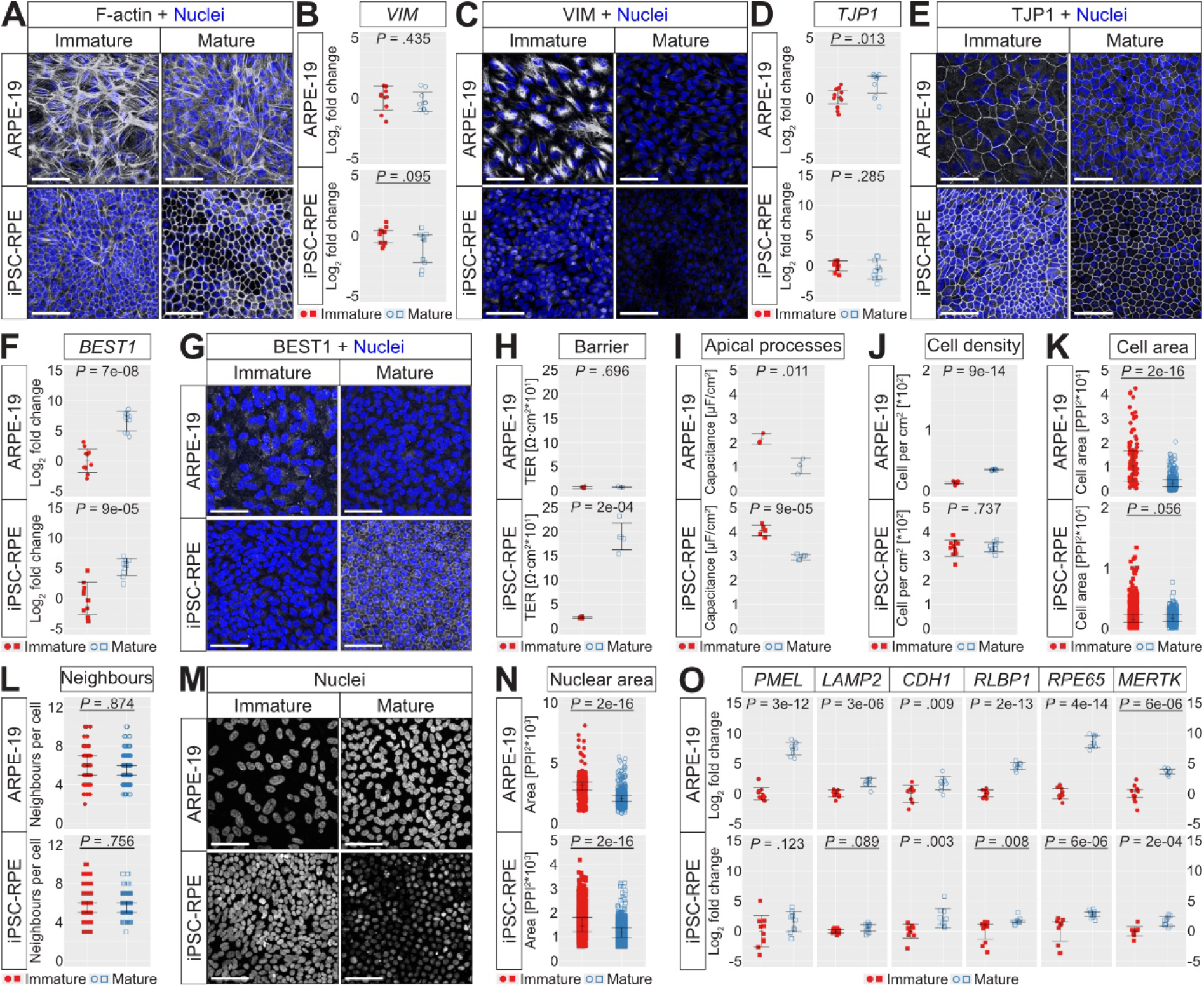
Mature ARPE-19 and iPSC-RPE cells show an epithelial phenotype and increased RPE-specific markers. **(A)** Immature ARPE-19 show mesenchymal F-actin with stress fibres. Mature ARPE-19 display epithelial-like F-actin with fewer stress fibres. iPSC-RPE exhibit circumferential F-actin, with stress fibres present in immature, but absent in mature cells (**Figure 1 - video 1**). **(B)** *VIM* expression remains unchanged (*n* = 9-10). **(C)** Immature cells show perinuclear VIM, which is reduced in mature ARPE-19 and absent in mature iPSC-RPE cells (**Figure 1 - video 2**). **(D)** *TJP1* expression is elevated in mature ARPE-19 (*n* = 10). **(E)** Mature cells display organised circumferential TJP1 (**Figure 1 - video 3**). **(F)** *BEST1* expression is elevated in mature cells (*n* = 10). **(G)** Mature iPSC-RPE exhibit circumferential BEST1 (**Figure 1 - video 4**). **(H)** TER is higher in mature iPSC-RPE (*n* = 3/5). **(I)** Mature cells display decreased electric capacitance (*n* = 3/5). **(J)** Mature ARPE-19 show increased cell density (*n* = 10). **(K)** Mature ARPE-19 decrease in cell cross-sectional area (*n* = 104/404/682/619 per immature/mature ARPE-19/iPSC-RPE). **(L)** Both cell types have a median of 6 neighbours (*n* = 103/339/656/508 per immature/mature ARPE-19/iPSC-RPE). **(M)** Nuclei in immature cells are larger. **(N)** Cross-sectional nuclear area decreases in mature cells (*n* = 633/1626/2727/1773 per immature/mature ARPE-19/iPSC-RPE). **(O)** Mature cells exhibit higher expression of key RPE markers (*n* = 8-10). Data represent pooled results from three independent culture replicates. Non-parametric data (underlined) were analysed with the Wilcoxon test (Mdn and IQR); parametric data using independent-means *t*-test (M and SD). Tests were two-tailed, and statistical significance was defined as *P* < .050. Scale bar = 50 µm.

Vimentin (VIM), a cytoskeletal marker specifically expressed in mesenchymal but not epithelial cells, was analysed to assess cell phenotype. *VIM* transcript expression levels were comparable between immature and mature cells (**Figure 1B**). However, at protein level, immature cells exhibited increased VIM accumulation near the nucleus, a characteristic feature of mesenchymal cells, whereas mature ARPE-19 cells showed only faint VIM signals and no VIM accumulation was detectable in mature iPSC-RPE cells (**Figure 1C**, **Figure 1 - video 2**). No significant differences in VIM fluorescence intensity were observed in ARPE-19 and iPSC-RPE cells (**Figure 1 - supplement 1B**).

To compare immature with mature ARPE-19 and iPSC-RPE cells and assess their RPE maturation, general RPE markers were further analysed. Mature ARPE-19 cells exhibited a significant increase in tight junction protein zonula occludens 1 (*TJP1*) transcript expression compared with immature cells (*P* = .013), whereas no significant difference was observed in iPSC-RPE cells (**Figure 1D**). At protein level, TJP1 exhibited organised circumferential signals in mature ARPE-19 cells, located more apically than in immature cells (**Figure 1E**, **Figure 1 - video 3**). No significant differences in TJP1 fluorescence intensity were observed in ARPE-19 cells (**Figure 1 - supplement 1C**). Similarly, mature iPSC-RPE cells demonstrated a more structured, regular circumferential and clearly defined TJP1 staining compared with immature cells (**Figure 1E**). Overall, signal intensity was more intense in immature iPSC-RPE cells, where TJP1 structures appeared thicker and less defined compared with mature cells. In mature iPSC-RPE cells, TJP1 was positioned apically above the nuclear layer, a feature not observed in immature iPSC-RPE cells (**Figure 1 - video 3**). Fluorescence intensity analysis confirmed these observations, revealing a significant reduction in TJP1 intensity in mature iPSC-RPE cells compared with immature cells (*P* = .006) (**Figure 1 - supplement 1C**).

Bestrophin 1 (*BEST1*), a transmembrane protein specifically expressed at the basolateral membrane of mature RPE cells, showed a significantly higher transcript expression in mature ARPE-19 and iPSC-RPE cells compared with immature cells (*P* = 7e-08, *P* = 9e-05) (**Figure 1F**). At protein level, BEST1 staining was detectable in mature iPSC-RPE cells, with localisation observed at the central and basal sides of the cell (**Figure 1G**, **Figure 1 - video 4**). No significant differences in BEST1 fluorescence intensity were observed in ARPE-19 and iPSC-RPE cells (**Figure 1 - supplement 1D**). Western blot analysis of cell lysates further validated the presence of BEST1, showing a distinct 70 kDa band exclusively in mature iPSC-RPE cells (**Figure 1 - supplement 2**).

To assess cell barrier function, transepithelial resistance (TER) was measured (**Figure 1H**). While no significant difference was found in TER between ARPE-19 cells, mature iPSC-RPE cells demonstrated a significantly higher TER, with a mean value of 190.14 ± 27.70 Ω·cm^2^ compared with 22.47 ± 1.62 Ω·cm^2^ in immature cells (*P* = 2e-04). Cell capacitance, a negatively correlated indicator of presence of apical processes, was significantly reduced in both mature ARPE-19 and iPSC-RPE cells (*P* = .011, *P* = 9e-05), suggesting an increased presence of apical processes upon maturation (**Figure 1I**).

To further characterise cell properties, image analysis and measurement of cross-sectional area of cells and nuclei were performed (**Figure 1 - supplement 3**). Cell density was significantly increased in mature ARPE-19 cells compared with immature cells (*P* = 9e-14), with no significant difference observed in iPSC-RPE cells (**Figure 1J**). Correspondingly, cross-sectional cell area based on TJP1 staining was significantly reduced in mature ARPE-19 cells compared with immature cells (*P* = 2e-16), while no difference was noted in iPSC-RPE cells (**Figure 1K**). Analysis of neighbouring cells revealed that both immature and mature cells had a median of six neighbours per cell, with a range of three to ten neighbours (**Figure 1L**). Nuclei staining indicated variations in nuclear cross-sectional area (**Figure 1M**). Mature ARPE-19 and iPSC-RPE cells exhibited a significant reduction in nuclear area compared with immature cells (*P* = 2e-16, *P* = 2e-16) (**Figure 1N**).

An analysis of several RPE marker transcripts using quantitative real time polymerase chain reaction (qRT-PCR) revealed significant differences between immature and mature ARPE-19 and iPSC-RPE cells (**Figure 1O**). Mature ARPE-19 cells showed an increase in premelanosome protein (*PMEL*) and lysosome-associated membrane protein 2 (*LAMP2*) transcript expression compared with immature cells (*P* = 3e-12, *P* = 3e-06), with no significant difference observed in iPSC-RPE cells. Additionally, both mature ARPE-19 and iPSC-RPE cells exhibited significant upregulation of cadherin 1 (*CDH1*) (*P* = .009, *P* = .003), retinaldehyde binding protein 1 (*RLBP1*) (*P* = 2e-13, *P* = .008), retinoid isomerohydrolase (*RPE65*) (*P* = 4e-14, *P* = 6e-06), and MER proto-oncogene, tyrosine kinase (*MERTK*) transcript expression (*P* = 6e-06, *P* = 2e-04).

In summary, immature cells exhibited more pronounced F-actin stress fibres and mesenchymal VIM staining, while mature cells showed a more epithelial-like cytoskeleton. Mature iPSC-RPE cells displayed a more epithelial phenotype and significantly improved barrier function compared with immature cells. In conclusion, mature iPSC-RPE cells demonstrated the closest resemblance to healthy *in vivo* RPE cells in terms of RPE specificity and functionality.

### 2.2. GT335 and ARL13B cilia length is reduced in mature iPSC-RPE cells

Ciliation was analysed as another marker of RPE maturation by comparing the primary cilium in dysfunctional, immature, and healthy, mature RPE cells. Cilia were characterised by the analysis of polyglutamylated tubulin (GT335) found in the basal body and ciliary transition zone and ADP-ribosylation factor-like protein 13B (ARL13B), a small GTPase specifically localised to the ciliary axoneme. Co-localisation of GT335 and ARL13B was observed in both ARPE-19 and iPSC-RPE cells (**Figure 2A**, **Figure 2 - video 1**), with almost all cells being ciliated.

**Figure 2:**
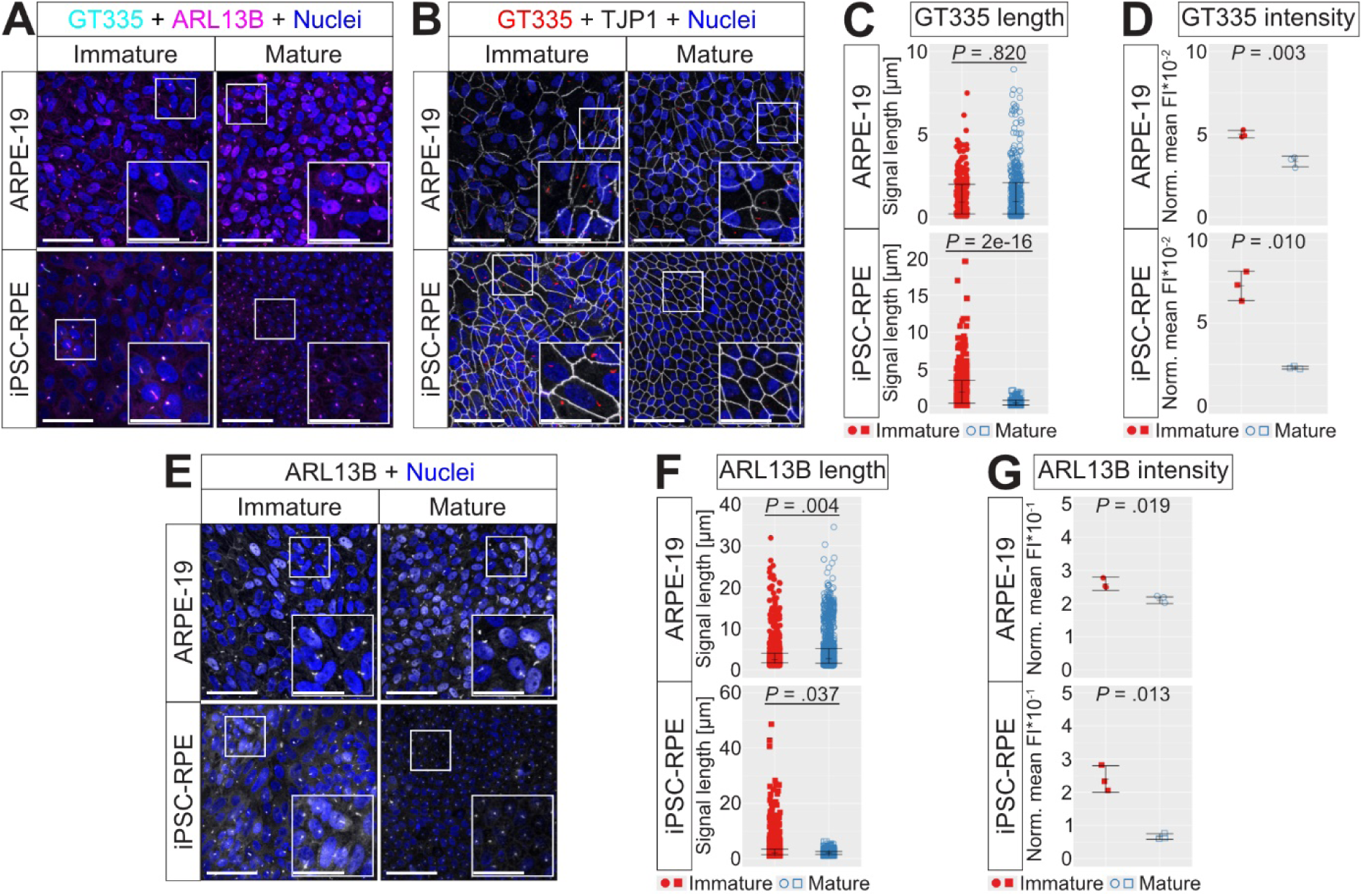
Mature iPSC-RPE cells show decreased cilia lengths. **(A)** GT335 and ARL13B signals show co-localisation (**Figure 2 - video 1**). The image extract in the lower right corner presents a twofold magnified view of the highlighted region in the small square. **(B)** GT335 signals appear as distinct dots in mature cells, whereas immature cells display more linear patterns (**Figure 2 - video 2**). The image extract in the lower right corner presents a twofold magnified view of the highlighted region in the small square. **(C)** GT335 signal length is decreased in mature iPSC-RPE cells (*n* = 264/459/321/140 per immature/mature ARPE-19/iPSC-RPE). **(D)** Fluorescence intensity of GT335 is reduced in mature ARPE-19 and iPSC-RPE cells compared with immature cells (*n* = 3). **(E)** ARL13B co-localised with the cell nucleus in immature cells and mature ARPE-19, whereas only mature iPSC-RPE display punctate and apical ARL13B staining independent of the basal nucleus (**Figure 2 - video 3**). The image extract in the lower right corner presents a twofold magnified view of the highlighted region in the small square. **(F)** ARL13B signal length is increased in mature ARPE-19 cells and decreased in mature iPSC-RPE cells (*n* = 1012/1231/1339/619 per immature/mature ARPE-19/iPSC-RPE). **(G)** Fluorescence intensity of ARL13B is reduced in mature ARPE-19 and iPSC-RPE cells compared with immature cells (*n* = 3). Data represent pooled results from two independent culture replicates. Non-parametric data (underlined) were analysed using the Wilcoxon test (Mdn and IQR); parametric data with independent-means *t*-test (M and SD). Tests were two-tailed, and statistical significance was defined as *P* < .050. Scale bar = 50 µm, scale bar image extract = 25 µm.

Mature cells exhibited reduced GT335 signals, which appeared as apical punctate dots, whereas immature cells showed more elongated signals (**Figure 2B**, **Figure 2 - video 2**). GT335 signal length was measured, revealing that mature iPSC-RPE cells had a significantly reduced GT335 length compared with immature cells (*P* = 2e-16), while ARPE-19 cells showed no difference (**Figure 2C**). Consistent with these results, fluorescence intensity analysis revealed a significant reduction in GT335 intensity in both mature iPSC-RPE (*P* = .010) and mature ARPE-19 cells (*P* = .003) compared with immature cells (**Figure 2D**). ARL13B staining confirmed the observations made with GT335, appearing as apical, punctate dot-like signals (**Figure 2E**, **Figure 2 - video 3**). In immature and mature ARPE-19 cells, as well as immature iPSC-RPE cells, ARL13B co-localised with the cell nucleus; however, this co-localisation was absent in mature iPSC-RPE cells. Consistent with GT335 findings, measurement of ARL13B length revealed a significant reduction in mature iPSC-RPE cells compared with immature cells (*P* = .037) (**Figure 2F**). In contrast, ARPE-19 cells displayed the opposite trend, with a significantly increased ARL13B length in mature cells compared with immature cells (*P* = .004). However, fluorescence intensity analysis showed significant reduction in ARL13B intensity in both mature iPSC-RPE (*P* = .013) and mature ARPE-19 cells (*P* = .019) compared with immature cells (**Figure 2G**).

In summary, these findings suggest that ciliary structures retract during maturation in iPSC-RPE cells, consistent with observations in rodent RPE. The absence of this retraction in mature ARPE-19 cells may indicate an incomplete maturation process in these cells.

### 2.3. ARPE-19 and iPSC-RPE cells produce C3 and its regulators with different expression and secretion patterns in immature and mature cells

Given the critical role of the complement system in AMD pathogenesis, as evidenced by genome-wide association studies linking numerous complement genes to AMD risk and the detection of complement components in drusen, we analysed the local complement production in our model, focussing on central complement component C3 and its regulators (**Figure 3A**). *C3* transcript levels were significantly increased in mature ARPE-19 and iPSC-RPE cells compared with immature cells (*P* = .001, *P* = 1e-07) (**Figure 3B**). In addition, both apical and basal C3 secretion were elevated in mature ARPE-19 cells (*P* = .008, *P* = .004) and iPSC-RPE cells (*P* = 7e-06, *P* = 3e-06) compared with immature cells (**Figure 3C**).

**Figure 3:**
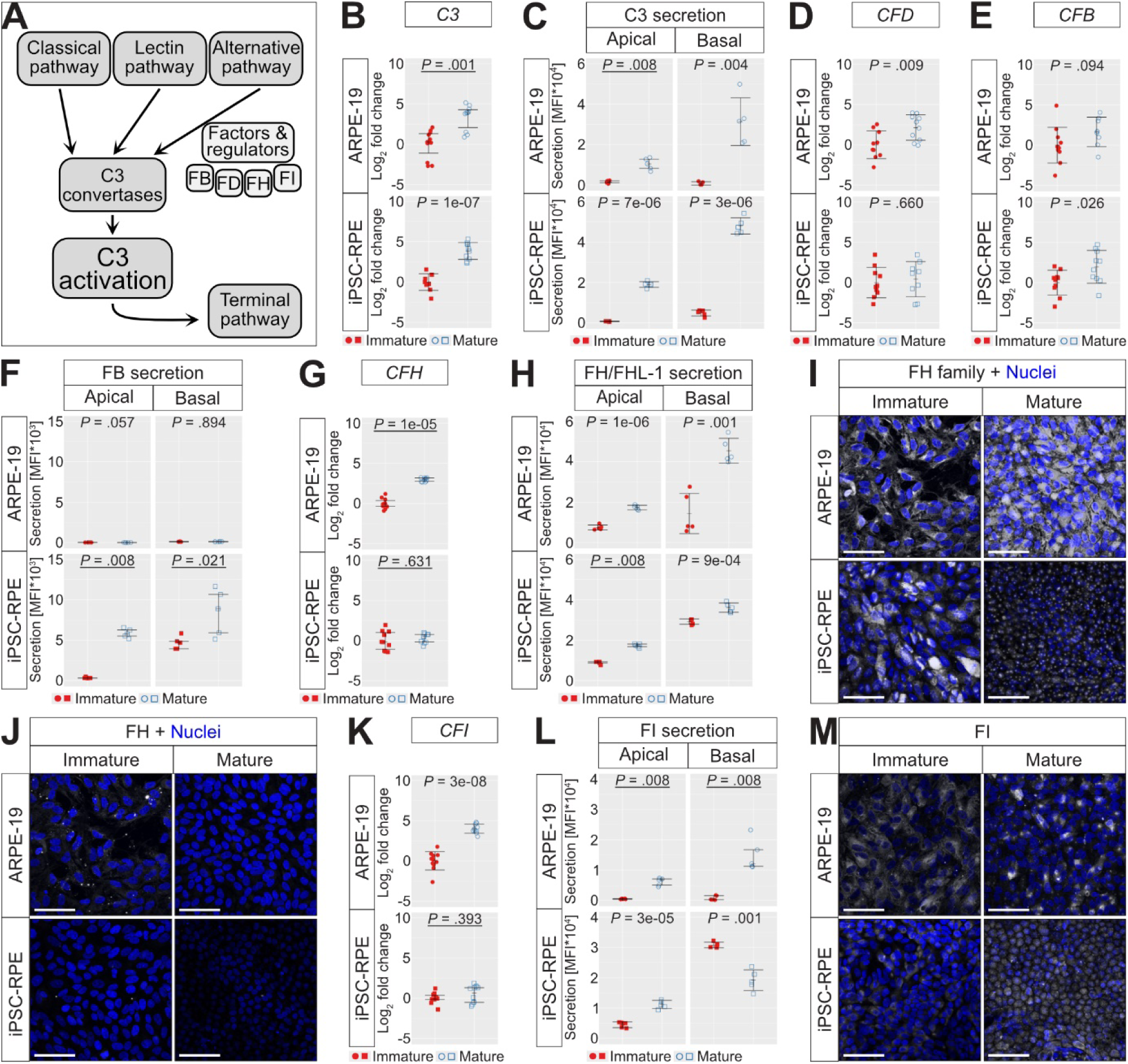
ARPE-19 and iPSC-RPE cells produce C3 and its regulators in different patterns during maturation. **(A)** Complement system overview. **(B)** *C3* expression (*n* = 10) and **(C)** apical/basal C3 secretion (*n* = 5) are increased in mature cells. **(D)** Mature ARPE-19 show increased *CFD* expression (*n* = 9-10). **(E)** *CFB* expression is increased in mature iPSC-RPE (*n* = 8-10). Apical/basal FB secretion is increased in mature iPSC-RPE (*n* = 5), but low in ARPE-19. *CFH* expression is elevated in mature ARPE-19 (*n* = 9-10). **(H)** FH/FHL-1 secretion is increased in mature cells (*n* = 5). **(I)** FH family staining signals are higher in mature ARPE-19 and immature iPSC-RPE, with apical FH localisation in mature iPSC-RPE (**Figure 3 - video 1**, antibody specificity in **Figure 3 - supplement 2**). **(J)** FH is intracellular in immature ARPE-19, but absent in others (**Figure 3 - video 2**, antibody specificity in **Figure 3 - supplement 3**). **(K)** *CFI* expression is elevated in mature ARPE-19 (*n* = 10). **(L)** FI secretion varies between immature and mature cells (*n* = 5). **(M)** FI signal distribution shows different patterns across cell types and maturation states (**Figure 3 - video 3**). Additional complement data are displayed in **Figure 3 - supplement 1**. Data represent pooled results from three independent culture replicates. Non-parametric data (underlined) were analysed using the Wilcoxon test (Mdn and IQR); parametric data with independent-means *t*-test (M and SD). Tests were two-tailed, and statistical significance was defined as *P* < .050. Scale bar = 50 µm.

Analysis of complement factor D (*CFD*) transcript expression revealed an increase in mature ARPE-19 cells compared with immature cells (*P* = .009), but not in iPSC-RPE cells (**Figure 3D**). FD was not detected above the assay cut-off in either apical or basal supernatants from ARPE-19 and iPSC-RPE cells (**Figure 3 - supplement 1**). *CFB* transcript expression was significantly elevated in mature iPSC-RPE cells compared with immature cells (*P* = .026) (**Figure 3E**), with corresponding increases in both apical and basal FB secretion (*P* = .008, *P* = .021) (**Figure 3F**). ARPE-19 cells showed no significant difference in *CFB* transcript expression, and apical and basal FB secretion remained low (**Figure 3E, F**).

*CFH* transcript expression was significantly increased in mature ARPE-19 cells compared with immature cells (*P* = 1e-05), with no significant difference observed in iPSC-RPE cells (**Figure 3G**). Both apical and basal FH/factor H-like protein 1 (FHL-1) secretion were elevated in mature ARPE-19 cells (*P* = 1e-06, *P* = .001) and iPSC-RPE cells (*P* = .008, *P* = 9e-04) compared with immature cells (**Figure 3H**). Immunostaining with an antibody against the FH protein family (**Figure 3 - supplement 2**) revealed fewer FH family signals in immature ARPE-19 cells compared with mature cells (**Figure 3I**), consistent with transcript and secretion data (**Figure 3G, H**). No significant differences in FH family fluorescence intensity were observed in ARPE-19 cells (**Figure 3 - supplement 4A**). In contrast, fewer FH family signals were observed in mature iPSC-RPE cells, along with a pronounced localisation in the apical region (**Figure 3 - video 1**). Fluorescence intensity analysis confirmed this observation, revealing a significant reduction in FH family intensity in mature iPSC-RPE cells compared with immature cells (*P* = .004) (**Figure 3 - supplement 4A**). Specific FH staining (**Figure 3 - supplement 3**) indicated intracellular FH in immature ARPE-19 cells, while no signals were detected in the other groups (**Figure 3J**, **Figure 3 - video 2**). Fluorescence intensity analysis confirmed these observations, revealing a significant increase in FH intensity in immature ARPE-19 cells compared with mature cells (*P* = .004), while no difference was observable in iPSC-RPE cells (**Figure 3 - supplement 4B**).

Similar to *CFH* transcripts (**Figure 3G**), *CFI* transcript expression was significantly elevated in mature ARPE-19 cells compared with immature cells (*P* = 3e-08), while no significant difference was observed in iPSC-RPE cells (**Figure 3K**). Both apical and basal FI secretion were increased in mature ARPE-19 cells compared with immature cells (*P* = .008, *P* = .008) (**Figure 3L**). In iPSC-RPE cells, apical FI secretion was significantly increased in mature cells (*P* = 3e-05), while basal FI secretion was decreased in mature cells (*P* = .001) compared with immature cells. Immunostainings showed FI signals with no distinct trend between immature and mature cells, except for a specifically apical localisation in mature iPSC-RPE cells (**Figure 3M**, **Figure 3 - video 3**). Fluorescence intensity analysis revealed a significant reduction in FI intensity in immature ARPE-19 cells compared with mature cells (*P* = 6e-04), while no difference was observable in iPSC-RPE cells (**Figure 3 - supplement 4C**).

It is noteworthy that components complement 1q (C1q), complement 2 (C2), complement 4 (C4), complement 4B (C4B), and properdin were secreted at low levels and proteins such as complement 5 (C5), complement 5a (C5a), complement 9 (C9), and mannan-binding lectin (MBL) were below the assay cut-off (**Figure 3 - supplement 1**).

To sum up, mature ARPE-19 and iPSC-RPE cells exhibit increased expression and secretion of C3 and its regulators compared with immature cells. Notably, the secretion profiles of FB and the complement regulators FH and FI vary significantly between immature and mature cells.

### 2.4. Mature iPSC-RPE cells exhibit increased secretion of active C3 forms compared with immature cells

With the recent focus on C3 in the context of AMD, highlighted by the use of the C3 inhibitor Pegcetacoplan as a therapeutic for GA, we specifically analysed local C3 processing in our model, focussing initially on its secretion profile. An overview of C3 activation and cleavage is presented in **Figure 4A**. The spontaneously activated and reactive form of C3, hydrolysed C3 (C3(H_2_O)), was detected in both apical and basal supernatants of RPE cells (**Figure 4B**). Mature iPSC-RPE cells secreted more C3(H_2_O) apically and basally than immature cells (*P* = .011, *P* = .008). The detection signal was low in ARPE-19 supernatants, with no difference in apical secretion and significantly increased basal secretion in mature ARPE-19 cells compared with immature cells (*P* = .008) (**Figure 4B**).

**Figure 4:**
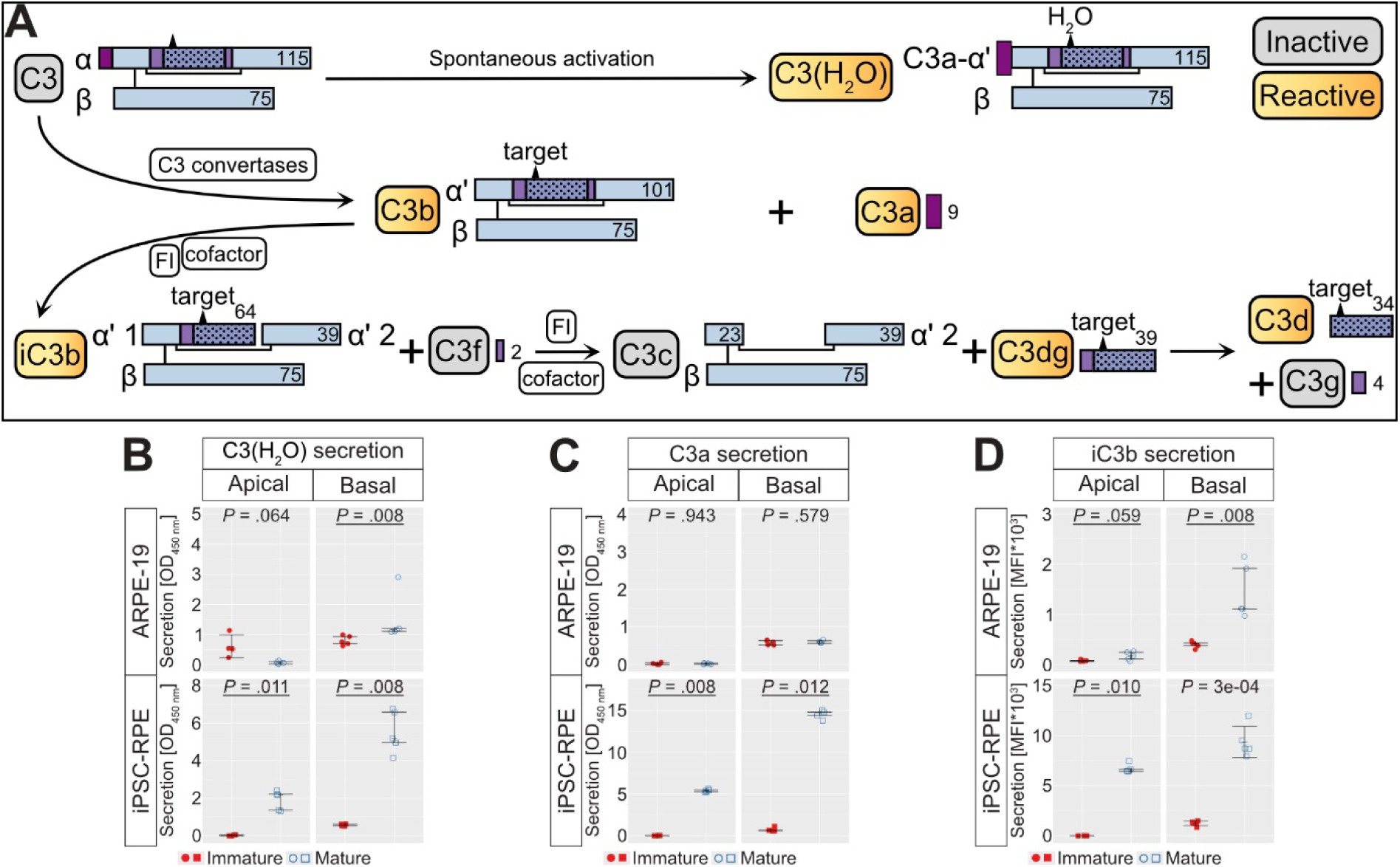
Secretion of active C3 forms is increased in mature iPSC-RPE cells compared with immature cells. **(A)** C3 consists of an α and β chain; the α chain contains an internal thioester bond (triangle). Spontaneous hydrolysis non-proteolytically activates C3 to C3(H_2_O), exposing the C3a neoepitope and a hydrolysed thioester. C3 cleavage by convertases generates reactive C3a and C3b, exposing the thioester for surface attachment. Further cleavage by FI produces iC3b, C3f, C3c, C3dg, C3d and C3g. Molecular weights are indicated in kDa. **(B)** Apical C3(H_2_O) secretion is elevated in mature iPSC-RPE, and basal secretion increases in mature ARPE-19 and iPSC-RPE cells (*n* = 4-5). **(C)** Apical/basal C3a secretion is elevated in mature iPSC-RPE (*n* = 5). **(D)** Apical/basal iC3b secretion increases in mature cells (*n* = 5). Data represent pooled results from three independent culture replicates. Non-parametric data (underlined) were analysed using the Wilcoxon test (Mdn and IQR); parametric data with independent-means *t*-test (M and SD). Tests were two-tailed, and statistical significance was defined as *P* < .050.

C3 cleavage by C3 convertases generates the inflammatory products C3a and C3b (**Figure 4A**). Apical and basal C3a secretion were elevated in mature iPSC-RPE cells compared with immature cells (*P* = .008, *P* = .012). In contrast, ARPE-19 cells exhibited very low apical C3a secretion and showed no differences in basal C3a secretion (**Figure 4C**).

Additionally, validation of the multiplex immunoassays confirmed that the kit C3 beads specifically detect C3, while C3b/iC3b beads primarily identify iC3b, with medium detection of C3c and minimal detection of C3b (**Figure 4 - supplement 1**). Consequently, the C3b/iC3b beads were redefined as iC3b beads for accurate representation. Apical iC3b secretion was higher in mature iPSC-RPE cells compared with immature cells (*P* = .010), with no significant difference observed in ARPE-19 cells (**Figure 4D**). Basal iC3b secretion was elevated in both mature ARPE-19 and iPSC-RPE cells compared with immature cells (*P* = .008, *P* = 3e-04).

In summary, mature ARPE-19 and iPSC-RPE cells show different patterns in the secretion of cell-autonomous, active C3 forms independent of added external complement components. In mature iPSC-derived RPE cells, the secretion of both C3(H_2_O) and C3a was detectable, whereas it was either absent or present at markedly lower levels in ARPE-19 cells. The secretion pattern in iPSC-RPE cells was consistent with the secretion of inactive C3, a pattern that was not observed in ARPE-19 cells.

### 2.5. Mature iPSC-RPE cells exhibit intracellular localisation of intact C3 β chain, unlike ARPE-19 and immature iPSC-RPE cells

To ensure comprehensive detection of various C3 fragments, we further analysed C3 processing in cell lysates using two antibodies: a monoclonal antibody targeting the C3c/C3d/C3f/C3 α‘ 2 epitope (antibody 1, **Figure 5A**) and a polyclonal antibody against C3 (antibody 2, **Figure 5D**). Negative controls for C3 antibody Western blots confirmed the specificity of the detected signals by demonstrating clear bands in purified proteins, while no signals were observed in cell culture media, as shown in **Figure 5 - supplement 1**.

**Figure 5:**
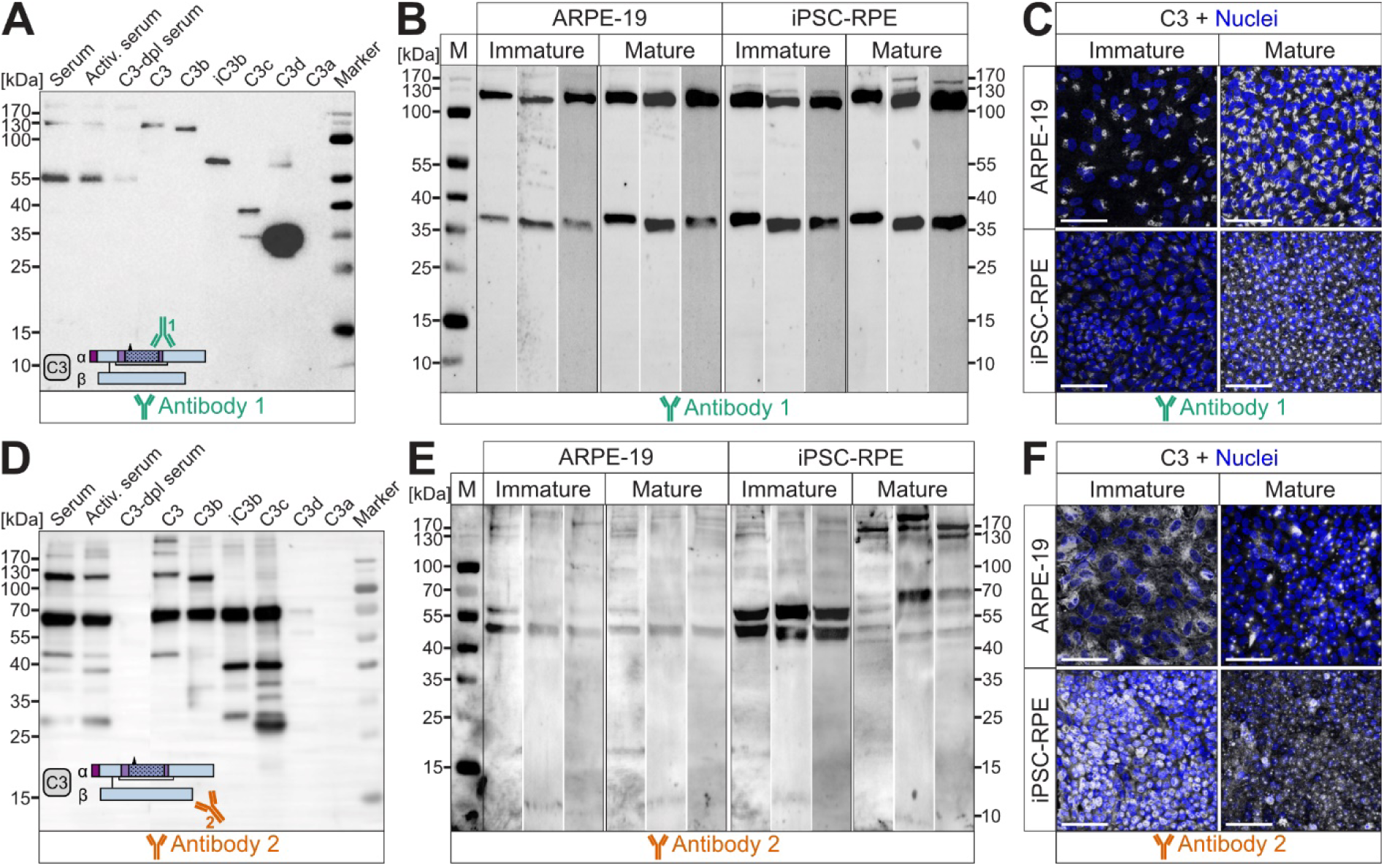
Mature iPSC-RPE cells show intracellular localisation of intact C3 β chain, absent in ARPE-19 and immature iPSC-RPE cells. **(A)** Specificity control for antibody 1 (localisation: anti-C3c/C3d/C3f/C3 α‘ 2) with purified proteins and human normal and C3-depleted (dpl) serum shows recognition of C3 α, α‘, α‘ 1, α‘ 2 chains, and C3d. **(B)** Three replicates of Western blot with immature and mature cell lysates using antibody 1 detect a ∼34 kDa band (C3d) and a ∼115 kDa band (C3 α). **(C)** Antibody 1 reveals intracellular C3 near the nucleus in ARPE-19 and immature iPSC-RPE. In mature iPSC-RPE, C3 is predominantly localised apically and circumferentially (**Figure 5 - video 1**). **(D)** Specificity control for antibody 2 (localisation: polyclonal anti-C3) detects C3 α, α‘, α‘ 2, and β chains. **(E)** Three replicates of Western blots with antibody 2 show C3 α and β chains in mature iPSC-RPE cells, with unique bands at ∼45 and ∼55 kDa in immature iPSC-RPE cells. **(F)** Antibody 2 staining shows widespread C3 signal in immature cells, with a basal cell border focus in mature cells (**Figure 5 - video 2**). Data represent pooled results from three independent culture replicates. Scale bar = 50 µm, M = marker.

Specificity testing for antibody 1 confirmed the recognition of C3 α, α‘, α‘ 1, α‘ 2 chains, and C3d in samples of purified C3, C3b, iC3b, C3c, and C3d, respectively (**Figure 5A**). Western blot analysis of cell lysates with antibody 1 revealed a ∼34 kDa band (C3d). A *∼*115 kDa band (C3 α) was more pronounced in mature ARPE-19 and iPSC-RPE cells (**Figure 5B**). A representative full membrane of the Western blot, displaying purified proteins alongside cell lysates, is shown in **Figure 5 - supplement 2A**. Immunostaining with antibody 1 showed specific intracellular C3 accumulations in ARPE-19 and iPSC-RPE cells (**Figure 5C**). In mature iPSC-RPE cells, C3 was localised apically and circumferentially in a punctate pattern, while in all other examined RPE cell variants, intracellular C3 staining was only observed near the cell nucleus (**Figure 5 - video 1**). No significant differences in C3 antibody 1 fluorescence intensity were observed in ARPE-19 and iPSC-RPE cells (**Figure 5 - supplement 3A**).

Specificity testing of antibody 2 demonstrated the recognition of C3 α, α‘, and α‘ 2 chains in samples of purified C3, C3b, iC3b, and C3c, as well as the β chain in samples of purified C3, C3b, iC3b, and C3c (**Figure 5D**). Western blot analysis using antibody 2 revealed strong bands corresponding to C3 α and β chains in mature iPSC-RPE cells (**Figure 5E**). In immature iPSC-RPE cells, two prominent bands were detected at ∼45 kDa and ∼55 kDa, perhaps representing fragments of the α or β chain (**Figure 5E**). These bands were almost absent in mature iPSC-RPE cells, suggesting differential processing of C3 during cell maturation. Immature and mature ARPE-19 cells displayed a band at *∼*45 kDa (**Figure 5E**). A representative full membrane of the Western blot, displaying purified proteins alongside cell lysates, is shown in **Figure 5 - supplement 2B**. Immunostaining with antibody 2 showed that C3 signal intensity was more diffuse throughout the cell body in immature cells, while mature cells exhibited more localised C3 signals (**Figure 5F**, **Figure 5 - video 2**). In particular, mature iPSC-RPE cells exhibited staining along the basal cell border, which was comparable to the cell border staining observed with antibody 1 (**Figure 5 - video 1**). No significant differences in C3 antibody 2 fluorescence intensity were observed in ARPE-19 and iPSC-RPE cells (**Figure 5 - supplement 3B**).

We further conducted immunoprecipitation to analyse the C3 β chain in RPE cell lysates (**Figure 5 - supplement 4**). Immunoprecipitation with a C3 β chain-specific antibody, followed by SDS-PAGE, Western blot, and detection with antibody 2 and the C3 β chain-specific antibody, revealed the presence of the full C3 β chain at approximately 75 kDa exclusively in mature iPSC-RPE cells, with no detectable signal in other groups.

In conclusion, the specificity and localisation of C3 and its fragments were assessed using two antibodies, revealing different expression patterns between immature and mature ARPE-19 and iPSC-RPE cells. Notably, mature cells showed more localised and intense C3 signals, while immature cells exhibited a broader distribution and stronger signals for specific C3 fragments. A key finding indicates that C3 is processed intracellularly, and the detected intracellular C3 cleavage products differ in size from known blood-derived C3 fragments. Additionally, the processing pattern appears to vary during the maturation process, notably, only mature iPSC-RPE cells exhibit the presence of the full β chain.

### 2.6. ARPE-19 cells display pronounced TβRI expression on protein level

To assess the susceptibility of RPE cells to TGF-β1-mediated stress, which induces a shift in cell expression patterns towards a mesenchymal state, the TGF-β1 signalling pathway was analysed in both immature and mature ARPE-19 and iPSC-RPE cells. Transforming growth factor-beta 1 (*TGFB1*) transcript levels were comparable in immature and mature ARPE-19 and iPSC-RPE cells (**Figure 6A**). The apical and basal secretion of TGF-β1 was significantly reduced in mature ARPE-19 (*P* = 3e-07, *P* = 2e-05) and iPSC-RPE cells (*P* = .012, *P* = .011) compared with immature cells (**Figure 6B**). In mature iPSC-RPE cells, TGF-β1 staining was primarily concentrated at the apical and central cell borders, whereas in all other groups it was more evenly distributed throughout the cell (**Figure 6C**, **Figure 6 - video 1**). Fluorescence intensity analysis revealed a significant reduction in TGF-β1 intensity in mature ARPE-19 cells compared with immature cells (*P* = .004), while the opposite trend was observed in iPSC-RPE cells (*P* = .027) (**Figure 6 - supplement 1A**).

**Figure 6:**
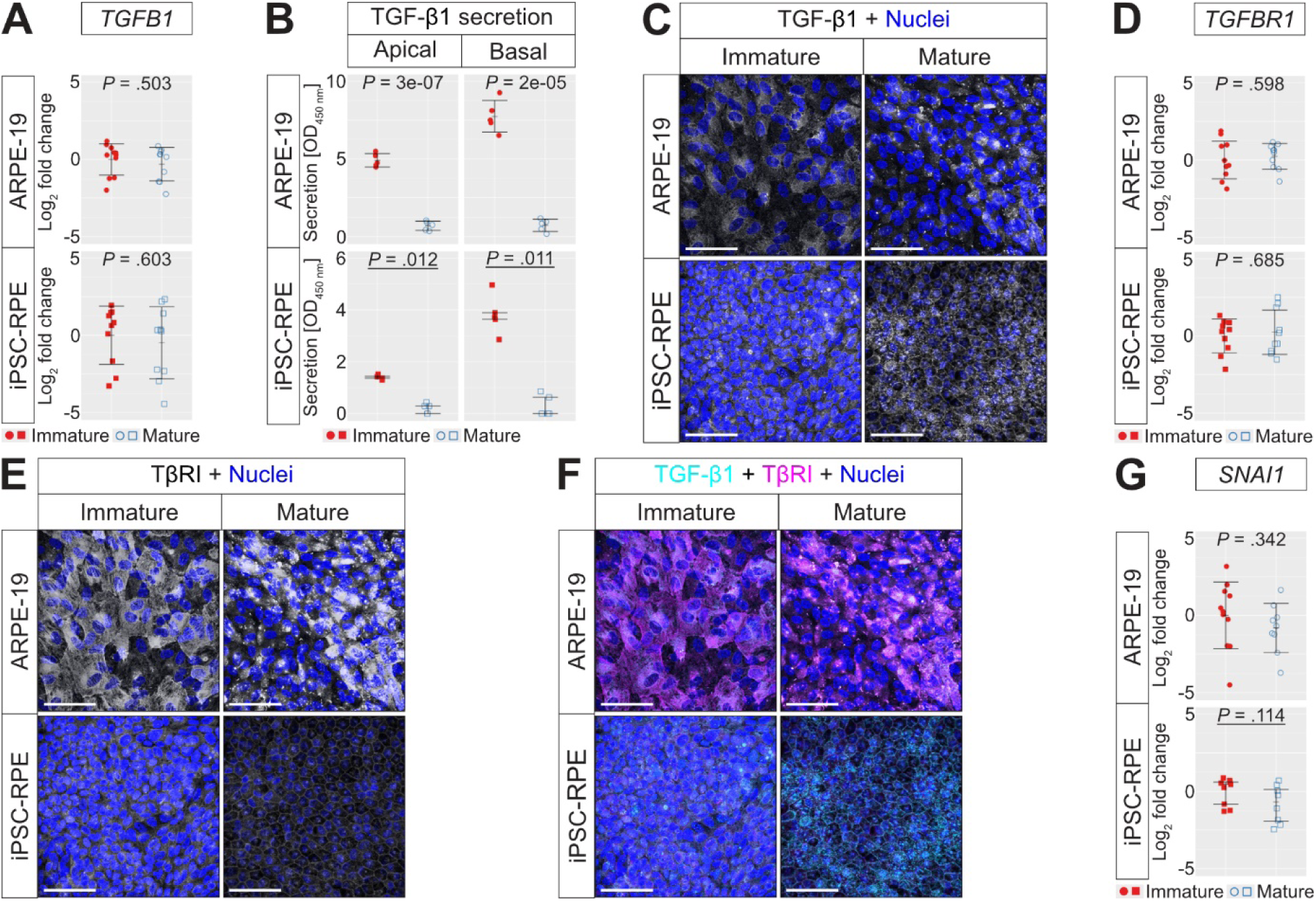
TβRI is strongly expressed in ARPE-19 cells. **(A)** No difference in *TGFB1* expression (*n* = 10). **(B)** TGF-β1 secretion is decreased in mature cells (*n* = 5). **(C)** TGF-β1 accumulates in mature cells, with a circumferential pattern in iPSC-RPE (**Figure 6 - video 1**). **(D)** No difference in *TGFBR1* expression (*n* = 10). **(E)** ARPE-19 display strong TβRI signals, while mature iPSC-RPE show a circumferential distribution (**Figure 6 - video 2**). **(F)** Co-staining of TGF-β1 (cyan) and TβRI (magenta) shows minimal co-localisation (white) in mature ARPE-19 cells (**Figure 6 - video 3**). **(G)** *SNAI1* expression shows no difference (*n* = 8-10). Data represent pooled results from three independent culture replicates. Non-parametric data (underlined) were analysed using the Wilcoxon test (Mdn and IQR); parametric data with independent-means *t*-test (M and SD). Tests were two-tailed, and statistical significance was defined as *P* < .050. Scale bar = 50 µm.

Transforming growth factor-beta receptor 1 (*TGFBR1*) transcripts were expressed at similar levels in both immature and mature ARPE-19 and iPSC-RPE cells (**Figure 6D**). At protein level, transforming growth factor-beta receptor 1 (TβRI) was detected across all cell types, but with varying cellular distributions (**Figure 6E**, **Figure 6 - video 2**). TβRI staining was bright in ARPE-19 cells, with immature ARPE-19 cells displaying fibre-like TβRI signals, whereas mature ARPE-19 cells exhibited TβRI concentrated in dot-like structures. Immature iPSC-RPE cells exhibited diffuse staining throughout the cell, except at the cell border, whereas mature iPSC-RPE cells displayed staining at the apical cell border with central dots (**Figure 6E**, **Figure 6 - video 2**). No significant differences in TβRI fluorescence intensity were observed in ARPE-19 and iPSC-RPE cells (**Figure 6 - supplement 1B**). Minimal co-localisation of TGF-β1 with TβRI was observed in mature ARPE-19 cells, while in the other groups, the two proteins were distributed separately (**Figure 6F**, **Figure 6 - video 3**).

Snail Family Transcriptional Repressor 1 (*SNAI1*), a key regulator of TGF-β-induced EMT, showed no differences in transcript levels between immature and mature cells (**Figure 6G**). In summary, ARPE-19 cells exhibited high expression of TβRI, indicating a high susceptibility to TGF-β1 treatment, while mature iPSC-RPE cells displayed weak TβRI signals, suggesting greater resistance to TGF-β1 treatment. Additionally, in iPSC-RPE cells, TGF-β1 and its receptor were localised in distinct cellular compartments, with non-overlapping distributions unlike the mature ARPE-19 cells where both were more closely co-localised, potentially contributing to the differential responsiveness to TGF-β1.

### 2.7. Stress induction of mature RPE cells by TGF-β1 addition is better tolerated by iPSC-RPE cells than ARPE-19 cells

Given the involvement of the TGF-β signalling pathway in AMD pathogenesis and its role in inducing EMT in RPE cells, mature ARPE-19 and iPSC-RPE cells were treated with or without 10 ng/mL human TGF-ꞵ1 under serum-free conditions. TGF-β1 treatment led to a significant increase in *TGFB1* transcript expression in ARPE-19 cells (*P* = .003), while no change was observed in iPSC-RPE cells (**Figure 7 - supplement 1A**). Basal TGF-β1 secretion was elevated in both cell types following treatment (*P* = .019, *P* = .029), while apical secretion remained unchanged (**Figure 7 - supplement 1B**). Immunostaining revealed diffuse intracellular TGF-β1 signals in TGF-β1-treated ARPE-19 cells, while signals were punctate in mature control ARPE-19 cells (**Figure 7 - supplement 1C, Figure 7- video 1**). In iPSC-RPE cells, mature control cells exhibited strong punctate signals, which became more diffuse and weak following treatment (**Figure 7 - supplement 1C, Figure 7 - video 1**). No significant differences in TGF-β1 fluorescence intensity were observed in ARPE-19 and iPSC-RPE cells (**Figure 7 - supplement 2A**).

**Figure 7:**
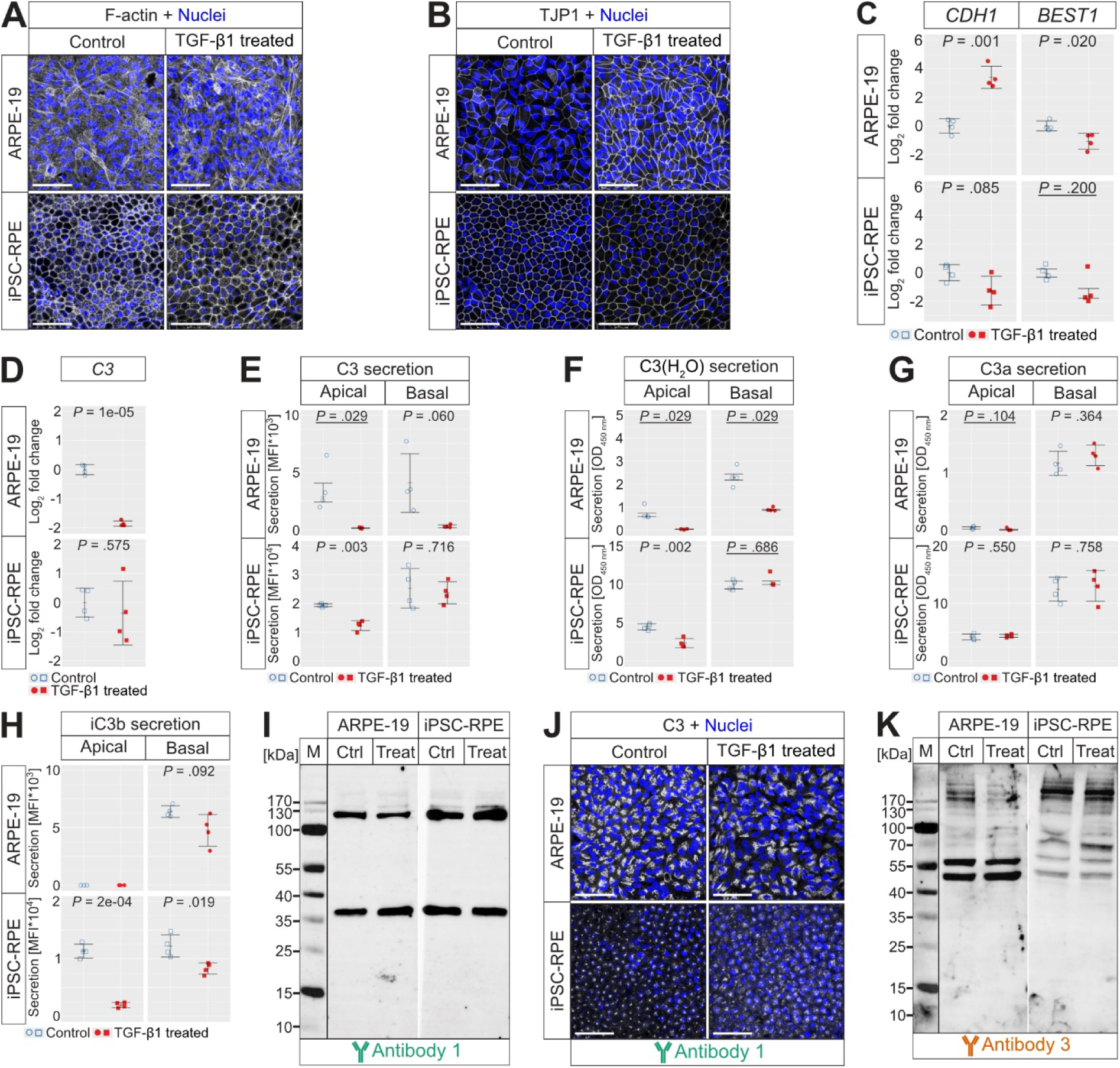
Mature RPE cells exposed to TGF-β1 show greater stress tolerance in iPSC-RPE compared with ARPE-19. **(A)** TGF-β1 increases basal stress fibres in ARPE-19 (**Figure 7** - **video 2**). **(B)** TGF-β1-treated ARPE-19 exhibit thickened TJP1 structures (**Figure 7 - video 3**). **(C)** TGF-β1-treated ARPE-19 display elevated *CDH1* and reduced *BEST1* transcript levels (*n* = 4). **(D)** TGF-β1 decreases *C3* transcript expression in ARPE-19, but not in iPSC-RPE (*n* = 4). **(E)** Apical C3 and **(F)** C3(H_2_O) secretion is reduced in both cell types following treatment, while basal C3(H_2_O) secretion is only reduced in treated ARPE-19 (*n* = 4). **(G)** C3a secretion remains unchanged (*n* = 4). **(H)** iC3b secretion is reduced in treated iPSC-RPE (*n* = 4). **(I)** Western blot with antibody 1 shows bands at ∼34 kDa (C3d) and ∼115 kDa (C3 α). **(J)** Immunostaining with antibody 1 reveals perinuclear C3 accumulations (**Figure 7 - video 4**). **(K)** Western blot with antibody 2 shows C3 α and β chains in iPSC-RPE, while ARPE-19 cells display bands at ∼45 and ∼55 kDa. Data represent pooled results from two independent treatment replicates. Non-parametric data (underlined) were analysed using the Wilcoxon test (Mdn and IQR); parametric data with independent-means *t*-test (M and SD). Tests were two-tailed, and statistical significance was defined as *P* < .050. Scale bar = 50 µm, M = marker.

Both control and TGF-β1-treated ARPE-19 cells exhibited basal stress fibres, whereas iPSC-RPE cells remained unaffected by the treatment, except for the absence of apical F-actin localisation in treated cells (**Figure 7A**, **Figure 7 - video 2**). No significant differences in F-actin fluorescence intensity were observed in ARPE-19 and iPSC-RPE cells (**Figure 7 - supplement 2B**). TGF-β1-treated ARPE-19 cells displayed thickened TJP1 filaments, while iPSC-RPE cells showed no significant structural changes between treatment groups, although apical TJP1 localisation in control iPSC-RPE cells was absent after TGF-β1 treatment (**Figure 7B**, **Figure 7 - video 3**). No significant differences in TJP1 fluorescence intensity were observed in ARPE-19 and iPSC-RPE cells (**Figure 7 - supplement 2C**). TGF-β1-treated ARPE-19 cells showed elevated transcript levels of *CDH1* (*P* = .001) and reduced *BEST1* transcript levels (*P* = .020) compared with control cells, with no changes in iPSC-RPE cells (**Figure 7C**).

Complement system analysis revealed a significant reduction in *C3* transcript expression in TGF-β1-treated ARPE-19 cells compared with control cells (*P* = 1e-05), while no changes were observed in iPSC-RPE cells (**Figure 7D**). Apical C3 secretion was reduced in both TGF-β1-treated ARPE-19 and iPSC-RPE cells compared with controls (*P* = .029, *P* = .003), whereas basal C3 secretion showed a tendency for TGF-β1-dependent decrease in ARPE-19 cells and remained unchanged in iPSC-RPE cells (**Figure 7E**).

Apical and basal C3(H_2_O) secretion was reduced in ARPE-19 cells following treatment compared with controls (*P* = .029, *P* = .029), while only apical secretion decreased in iPSC-RPE cells (*P* = .002) (**Figure 7F**). No differences in C3a secretion were observed in either cell type (**Figure 7G**). TGF-β1 treatment further decreased the apical and basal iC3b secretion in iPSC-RPE cells compared with control cells (*P* = 2e-04, *P* = .019), while in ARPE-19 cells, basal secretion remained unchanged and apical secretion was undetectable (**Figure 7H**).

A Western blot using C3 antibody 1 detected bands at ∼34 kDa (C3d) and ∼115 kDa (C3 α) in both control and TGF-β1-treated cells, with no notable differences (**Figure 7I**). Immunostaining with antibody 1 showed a slight increase in perinuclear C3 accumulation in TGF-β1-treated ARPE-19 cells (**Figure 7J**), however, no significant differences in C3 antibody 1 fluorescence intensity were observed in ARPE-19 (**Figure 7 - supplement 2D**). Control iPSC-RPE cells further displayed a distinct apical C3 localisation above the nuclear layer that was absent in TGF-β1-treated cells (**Figure 7J, Figure 7 - video 4**). Fluorescence intensity analysis revealed a significant increase in C3 antibody 1 intensity in TGF-β1-treated iPSC-RPE cells compared with control cells (*P* = .021) (**Figure 7 - supplement 2D**)

A Western blot with C3 antibody 2 revealed visible bands of C3 α and β chains in iPSC-RPE cells, while ARPE-19 cells displayed stronger bands at ∼45 and ∼55 kDa compared with iPSC-RPE cells (**Figure 7K**).

The secretion of FB, as well as transcript expression and secretion of complement regulators FH and FI, were further investigated after TGF-β1 treatment (**Figure 7 -supplement 1**). TGF-β1 treatment had no significant effect on apical and basal FB secretion in ARPE-19 cells (**Figure 7 - supplement 1D**). In iPSC-RPE cells, basal FB secretion significantly increased after treatment compared with controls (*P* = .041), while apical secretion remained unchanged (**Figure 7 - supplement 1D**). *CFH* transcript expression was significantly decreased in TGF-β1-treated ARPE-19 cells compared with control cells (*P* = .029), while the reduction was not significant in iPSC-RPE cells (**Figure 7 - supplement 1E**). Apical FH/FHL-1 secretion was significantly lower in TGF-β1-treated ARPE-19 and iPSC-RPE cells compared with control cells (*P* = .001, *P* = .008), while basal secretion showed a significant reduction only in ARPE-19 cells (*P* = .029) (**Figure 7 - supplement 1F**). *CFI* transcript expression was also significantly reduced in TGF-β1-treated ARPE-19 cells compared with control cells (*P* = .002), with no significant changes in iPSC-RPE cells (**Figure 7 - supplement 1G**). Apical FI secretion was similarly reduced in both TGF-β1-treated ARPE-19 and iPSC-RPE cells compared with control cells (*P* = .029, *P* = .003), with basal FI secretion showing a significant reduction only in ARPE-19 cells (*P* = .029) (**Figure 7 - supplement 1H**).

In conclusion, mature iPSC-RPE cells were considerably more resistant to TGF-β1-mediated stress compared with ARPE-19 cells. However, regarding complement secretion, TGF-β1 induced a similar pattern in both mature cell types, characterised by a reduction in the secretion of C3, FH/FHL-1, and FI, while the release of activated C3a remained unchanged and cellular cleavage pattern of C3 was not altered.

## 3. Discussion

### 3.1. Evaluating RPE cell maturation as a model for disease-related RPE pathology

The need for effective treatments for non-exudative AMD is urgent, but the complexity of this disease has hindered a complete understanding of its diverse pathomechanisms. Several research groups have studied HRF and their origin as indicators of AMD progression, identifying morphologically distinct RPE cells in and near atrophic regions of the retina. These cells lose their specificity, undergo dysmorphic changes, transdifferentiate, and migrate through the retina, contributing to disease pathology (Balaratnasingam et al., 2017; Bonilha et al., 2020; Cao et al., 2021; Gambril et al., 2019; Zanzottera, Messinger, Ach, Smith, & Curcio, 2015; Zanzottera, Messinger, Ach, Smith, Freund, et al., 2015). EMT processes, triggered by ageing, smoking, and other AMD risk factors, have been implicated as a potential cause of RPE pathology and HRF development. RPE cells undergoing EMT lose their epithelial characteristics, downregulate cell adhesion markers (e.g., TJP1 and CDH1), adopt a mesenchymal morphology, and upregulate mesenchymal markers (e.g., VIM), enabling them to migrate (Bonilha et al., 2020; Ghosh et al., 2018).

Given the potential of HRF as an indicator of AMD progression, we aimed to further investigate the complement system in dysfunctional RPE cells, potentially resembling those identified as HRF. Therefore, we developed a model featuring both immature and mature RPE cells, representing the dysfunctional unspecific RPE and healthy specific RPE, respectively. For this model, we utilised the reverse process of EMT, known as MET, which occurs during RPE cell maturation *in vitro*, where immature RPE cells transition from initially mesenchymal characteristics to an epithelial, RPE-specific phenotype (Chtcheglova et al., 2020; Tian et al., 2018).

The mature cells, particularly mature iPSC-RPE cells, exhibited increased expression of RPE markers across various signalling pathways, enhanced barrier function with high TER values, and the development of apical processes, a hexagonal TJP1 distribution, and circumferential F-actin structures typical of epithelial RPE cells (**Figure 1**, **Figure 8**). In contrast, immature cells displayed a stressed phenotype, with elongated stress fibres and increased VIM signals characteristic of mesenchymal cells; differences in cell density, size, and nuclear area observed between immature and mature cells (**Figure 1**, **Figure 8**). These findings align with other studies using different RPE cultivation techniques, showing that under optimal culture conditions, using differentiation-promoting media supplementation and extended cultivation periods, human primary RPE, ARPE-19, and stem cell-derived RPE cells achieve a mature, unstressed phenotype with high expression of RPE markers, polarity, hexagonal TJP1 distribution, circumferential epithelial F-actin localisation, smaller cell size, and strong barrier formation (Al-Ani et al., 2020; Miyatani et al., 2024; Samuel et al., 2017; Tian et al., 2018). Notably, the RPE-typical apical location of the F-actin belt and TJP1, along with the more basal localisation of BEST1 and the cell nucleus, as described by Lindell et al. and Marmorstein et al., was only observed in our mature iPSC-RPE cells but not in mature ARPE-19 cells, using confocal microscopy (**Figure 1**, **Figure 1 - video 1, Figure 1 - video 3, Figure 1 - video 4**) (Lindell et al., 2023; Marmorstein et al., 2000). In contrast, studies using suboptimal cultivation methods, e.g., less supplemented media, high serum concentrations, shorter cultivation times, or frequent passaging, have reported stressed, fibroblast-like human primary RPE, ARPE-19, and stem cell-derived RPE cells with inadequate apical junctional complexes, formation of actin stress fibres, and reduced RPE marker expression (Al-Ani et al., 2020; Boles et al., 2020; Chtcheglova et al., 2020; Miyatani et al., 2024; Radeke et al., 2015; Samuel et al., 2017; Sripathi et al., 2021; Tian et al., 2018).

**Figure 8:**
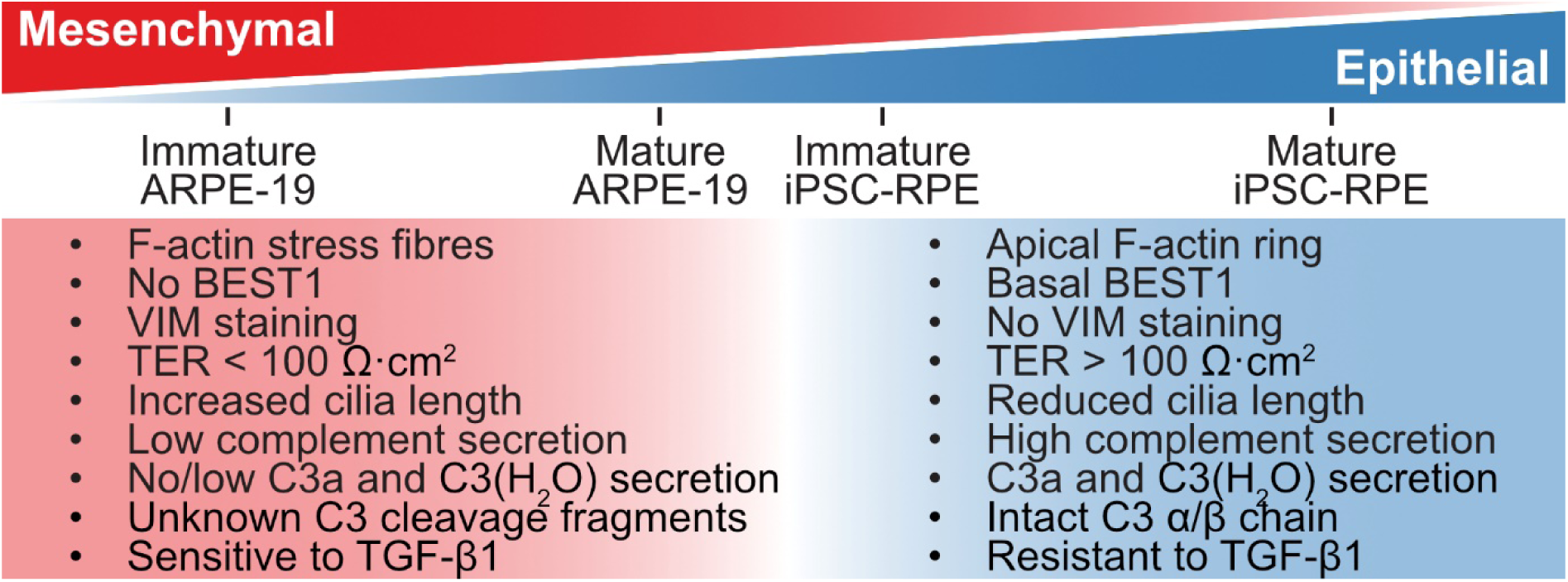
Immature and mature ARPE-19 and iPSC-RPE cells show distinct mesenchymal and epithelial characteristics. The results of the comparative analysis of mesenchymal and epithelial phenotypes in immature and mature ARPE-19 and iPSC-RPE cells are shown, categorised by maturation status. Immature ARPE-19 cells exhibit mesenchymal characteristics, including F-actin stress fibres, absence of BEST1, VIM expression, low TER (< 100 Ω·cm^2^), increased cilia length, minimal complement secretion, and intracellular C3 cleavage fragments. These cells also show sensitivity to TGF-β1 treatment. Mature ARPE-19 cells retain a mesenchymal phenotype that is more pronounced than that of immature iPSC-RPE cells. In contrast, mature iPSC-RPE cells demonstrate an epithelial RPE-specific phenotype with apical F-actin rings, basal BEST1, absence of VIM, high TER (> 100 Ω·cm^2^), reduced cilia length, and robust secretion of complement components C3a and C3(H_2_O), as well as intact C3 α/β chains. This phenotype, together with resistance to TGF-β1-induced stress, supports mature iPSC-RPE cells as the most physiologically relevant RPE model in this study.

The primary cilium plays a critical role in RPE cell maturation, contributing to intracellular transport and signalling pathways necessary for cellular functionality (May-Simera et al., 2018). Disruption of ciliary function can lead to RPE dysmorphia, dysfunction, EMT-like changes, and ultimately visual impairment and retinal dystrophy (Kretschmer et al., 2023; Ning et al., 2023; Schneider et al., 2021). Ciliogenesis, which enhances RPE maturation and polarisation, is crucial for achieving a natural cell cycle exit required for full maturation (May-Simera et al., 2018). A previous study on murine RPE demonstrated that ciliogenesis occurs during cell maturation, with cilia disassembling and retracting once the cells reach a mitotic, mature state (Patnaik et al., 2019). Given that ciliogenesis varies during RPE maturation and differs between epithelial and mesenchymal cells, we further characterised the primary cilium in our model (S. Kim & Dynlacht, 2013; May-Simera et al., 2018; S. Sorokin, 1962; S. P. Sorokin, 1968). Our findings demonstrate a significant reduction in the lengths of the ciliary proteins GT335 and ARL13B in mature iPSC-RPE cells compared with immature cells (**Figure 2**, **Figure 2 - video 2, Figure 2 - video 3**). In contrast, this trend was not observed in ARPE-19 cells, where the lengths of these proteins were either unchanged or showed an opposing increase upon maturation.

These observations suggest that our mature iPSC-RPE cells may have reached a stage of full maturation, probably undergoing ciliary retraction. To the best of our knowledge, the process of deciliation after maturation has not been previously reported in human iPSC-RPE cells, and requires further investigation with additional ciliary proteins to confirm this phenomenon.

### 3.2. Local production and secretion of complement components by RPE cells is dependent on the maturation

In the context of HRF in non-exudative AMD, we developed a model using ARPE-19 and iPSC-RPE cells at varying stages of maturation, to investigate the cellular complement system, a key process involved in AMD disease progression. Genome-wide association studies have linked variants in complement genes, including *C3*, *CFB*, *CFH*, and *CFI*, to increased AMD susceptibility (Despriet et al., 2009; Edwards et al., 2005; Fritsche et al., 2013, 2016; Hageman et al., 2005; Klein et al., 2005; Maller et al., 2007; Park et al., 2009; Yates et al., 2007). Furthermore, components and regulators of the complement system, such as C3, C3b, iC3b, C3dg, FH, and FI, have been identified in drusen, deposits that are characteristic of AMD found between the RPE and Bruch’s membrane (Anderson et al., 2002, 2010; Crabb et al., 2002; P. T. Johnson et al., 2006; L. V. Johnson et al., 2001; Laine et al., 2007). A study by Sharma et al. demonstrated that treatment of iPSC-RPE cells with complement-competent serum, serving as a source of activated anaphylatoxins, led to the development of an AMD-like cellular phenotype (Sharma et al., 2021). This included the formation of sub-RPE lipid deposits, increased F-actin stress fibres, and the loss of hexagonal morphology and epithelial phenotype.

Despite the blood-retina barrier separating the retina from the systemic complement system produced in the liver and establishing the immune privilege of the eye, recent studies have demonstrated that complement components are locally produced in the murine and human RPE/choroid complex (Anderson et al., 2010; Enzbrenner et al., 2021; Pauly et al., 2019; Rutar et al., 2012; Schäfer et al., 2017; Zauhar et al., 2022). Specifically, *in vitro* studies involving ARPE-19, human primary RPE, and iPSC-RPE cells have shown that components such as C3, C3a, C3b, C3d, FB, FH, and FI are expressed and produced independently of exogenous sources and can be modulated by oxidative stress or pro-inflammatory cytokine exposure (Y. H. Kim et al., 2009; Luo et al., 2011; Schäfer et al., 2020, 2021; Sugita et al., 2018; Trakkides et al., 2019).

Our study focused on the production of complement components in immature and mature ARPE-19 and iPSC-RPE cells. We analysed the transcript expression, along with apical and basal secretion and intracellular localisation of several complement components and regulators (**Figure 3**, **Figure 3 - supplement 1**). Notably, the expression and secretion of these factors were dependent on the maturation of the cells, with mature cells showing primarily increased apical and basal secretion compared with immature cells including components such as C3, FB, FH/FHL-1, and FI (**Figure 3**). This indicates a local complement production in both ARPE-19 and iPSC-RPE cells, with maturation-dependent changes in complement activity and regulatory mechanisms. Moreover, in mature iPSC-RPE cells, we observed distinct trends in polarised secretion of components like C3, FI, and FH/FHL-1, which was evidenced by the specific apical intracellular localisation of these factors in the mature cells (**Figure 3**, **Figure 3 - video 1, Figure 3 - video 3, Figure 5, Figure 5 - video 1, Figure 5** - **video 2).**

Several studies have conducted transcriptomic analyses comparing poorly differentiated RPE cells lacking typical RPE morphology with control cells with an RPE-specific phenotype. Sripathi et al. reported that enzymatic dissociation-induced EMT in iPSC-RPE cells resulted in significant modulation of transcript levels for AMD-associated genes, including decreased expression of *CFH* and *CFI* compared with controls, consistent with our findings in immature ARPE-19 cells (Sripathi et al., 2021). However, their observed increase in *C3* transcript levels after EMT induction contrasts with our observation of reduced *C3* transcript expression in immature cells. Al-Ani et al. noted a non-significant trend toward increased *CFH* expression in confluent mature ARPE-19 cells compared with immature cells, which also supports our findings (Al-Ani et al., 2020). Finally, we analysed the dataset of Samuel et al. using the National Center for Biotechnology Information (NCBI) Gene Expression Omnibus (GEO) tool GEO2R (Barrett et al., 2012; Samuel et al., 2017). We observed elevated *CFH*, *CFI*, and *C3* transcript levels in ARPE-19 cells with an RPE-typical phenotype following four months of cultivation, compared with shortly cultivated cells, which aligns with our data on mature ARPE-19 cells (data accessible at NCBI GEO database, accession GSE88848) (**Appendix 2A**).

Comparing our maturation model with other studies that analysed deficiently matured RPE cells with RPE-specific mature cells is complicated due to the use of different cell lines and methods for achieving immature and mature phenotypes. Our study, along with those of Samuel et al. and Al-Ani et al., focused on MET during RPE cell cultivation, while Sripathi et al. used enzymatic dissociation as an EMT-inducing treatment (Al-Ani et al., 2020; Samuel et al., 2017; Sripathi et al., 2021). Despite these methodological differences, a consistent trend is observed: RPE-specific mature RPE cells express increased *CFH* and *CFI* transcript levels, with elevated *C3* transcript expression observed in the MET-based maturation models by Samuel et al., and in our study. These results support the hypothesis that complement components are locally expressed in the RPE and are regulated by the maturation state and morphology of the RPE.

When investigating the local production of complement components by RPE cells *in vitro*, it is crucial to account for potential extracellular sources of complement, particularly from sera used in the culture medium. To mitigate this issue and eliminate complement activity, we utilised heat-inactivated fetal bovine serum for ARPE-19 cells, and knockout serum replacement for iPSC-RPE cells during cultivation (Soltis et al., 1979; Triglia & Linscott, 1980). Additionally, appropriate positive and negative controls, along with medium blanks, were employed as controls in further experiments to ensure the accuracy of our observations. These measures enabled the analysis of the local production of complement components in RPE cells, underscoring the significance of maturation-dependent changes in complement activity and secretion patterns.

### 3.3. Local C3 activation in RPE cells increases under pathological conditions

In recent years, the approval of complement inhibitors by the Food and Drug Administration has introduced a new therapeutic approach for GA. Pegcetacoplan (Syfovre™), a C3 inhibitor, prevents the cleavage of C3 into C3a and C3b, thus mitigating excessive complement activation (Liao et al., 2020). Clinical trials, such as the FILLY phase 2 and OAKS phase 3, demonstrated that monthly or bimonthly intravitreal injections of Pegcetacoplan significantly slowed GA lesion growth over at least twelve months, though this effect did not reach statistical significance in the DERBY phase 3 trial (Dascalu et al., 2024; Heier et al., 2023; Liao et al., 2020; Steinle et al., 2021). Moreover, Pegcetacoplan-treated eyes exhibited reduced photoreceptor thinning outside the RPE-atrophic area compared with sham-treated eyes (Pfau et al., 2022). Importantly, tested genetic risk factors did not significantly impact the therapeutic effect of Pegcetacoplan, indicating its efficacy is independent of these factors (Liao et al., 2020).

With the current emphasis on C3 activation and inhibition in the context of GA therapy, we aimed to analyse C3, its isoforms, and processing in our RPE cell model. C3 is composed of α and β chains, with the α chain, which contains an internal thioester bond (**Figure 4A**). Spontaneous hydrolysis activates inactive C3 non-proteolytically to the reactive form, C3(H_2_O), exposing a C3a neoepitope and a hydrolysed thioester. This form binds to hydroxyl or amino groups on target surfaces. Furthermore, cleavage of inactive C3 by C3 convertases generates the two fragments C3a and C3b. C3a is a small soluble molecule and pro-inflammatory anaphylatoxin that is inactivated by removal of the C-terminal arginine, resulting in the production of C3a-desArg. C3b exposes its thioester bond for covalent binding and promotes opsonisation and further complement activation. C3b is cleaved by FI and its co-factors to produce iC3b, which retains opsonising capabilities but no longer promotes complement amplification. iC3b is further degraded into inactive C3c and C3dg, with C3dg acting as an immune mediator prior to its degradation into C3d and C3g, completing the complement activation process (Merle, Church, et al., 2015; Merle, Noe, et al., 2015; Ricklin et al., 2016). Focusing on secretion, we observed a significant increase in apical and basal secretion of active C3 forms, including C3(H_2_O), C3a, and iC3b, in mature iPSC-RPE cells compared with immature cells, while ARPE-19 cells exhibited very low or no secretion of these proteins (**Figure 4B, C, D**). Our findings suggest that inactive C3 is produced and processed into active forms in RPE cells and independent of added complement components. Active C3 forms are further secreted both apically and basally with mature iPSC-RPE cells secreting at higher levels, potentially serving as extracellular opsonins (iC3b) or chemotactic agents (C3a) to maintain the immune privilege of the retina.

We further examined intracellular C3 isoforms and activation products. Mature iPSC-RPE cells displayed a greater number of punctate and apically localised signals of C3 (**Figure 5C, F**, **Figure 5 - video 1**), whereas immature cells exhibited stronger signals for cleaved C3 forms (**Figure 5E, F**, **Figure 5 - video 2**). Furthermore, signals specifically for the α chain and C3d increased during maturation, while the full β chain was only present in mature iPSC-RPE cells (**Figure 5B, E, Figure 5 - supplement 4**). This indicates that the intracellular accumulation of C3 isoforms and activation products varies with maturation. Additionally, intracellular C3 activation products in RPE cells differ from known blood-derived C3 fragments, such as the ∼45 and ∼55 kDa C3 fragments (**Figure 5E, F**, **Figure 5 - video 2**).

In previous studies, increased C3 activation products, including C3(H_2_O), C3a, C3b, C3d, C3c, and C3dg, were detected in cell lysates or supernatants of various in vitro RPE models, including complement treatment, oxidative stress treatments or altered extracellular matrix (ECM) conditions (Chinchilla & Fernandez-Godino, 2021; Ebrahimi et al., 2013; Fernandez-Godino, 2018; Fields et al., 2017; Kunchithapautham et al., 2014; Schäfer et al., 2020, 2021; L. Wang et al., 2014; Y. Wang et al., 2023). Collectively, the findings of other groups indicate that C3 activation occurs in RPE cells under stressed pathological conditions, such as oxidative stress or abnormal ECM. This trend of increased C3 forms is consistent with our observation of higher accumulation in immature iPSC-RPE cells. However, while other studies observed increased C3 activation product secretion, we found that mature iPSC-RPE cells primarily secrete more C3 activation products. Nevertheless, it is important to note that models utilising oxidative stress induction or ECM modification are not directly comparable to our maturation-based model. Furthermore, to the best of our knowledge, no other group has specifically analysed both the secretion and intracellular distribution of C3 and its activation products in human RPE cells using different antibodies targeting various C3 isoforms as we have done in our study. This limits direct comparisons between our work and previous studies. Nonetheless, the findings suggest that human RPE cells activate locally produced C3 autonomously and intracellularly, either retaining the products or secreting them apically and basally. This process of C3 activation appears to be enhanced under immature or pathological conditions.

The mechanism of maturation-dependent C3 cleavage in RPE cells requires further investigation. A potential hypothesis is that the incomplete structural maturation of C3 in immature RPE cells could contribute to its susceptibility to autoproteolytic degradation. This is supported by the absence of the C3 β-chain in immature and mature ARPE-19 cells, as well as in immature iPSC-RPE cells, which contrasts with the intact α- and β-chains observed in mature iPSC-RPE cells (**Figure 5E**). The presence of an autolytically cleaved 46-47 kDa fragment of the C3 α-chain during sodium dodecyl sulphate (SDS)-based denaturation, as previously reported by Isaac et al. and Sim et al., aligns with the 45 kDa band detected in immature iPSC-RPE cells and ARPE-19 cells in this study (Isaac & Isenman, 1992; Sim & Sim, 1983). These findings suggest that autoproteolytic processes may either occur intracellularly or during sample preparation for SDS polyacrylamide gel electrophoresis (SDS-PAGE). In contrast, mature iPSC-RPE cells, with their fully assembled and stable C3 structure, appear less susceptible to such degradation.

As another possible mechanism, intracellular FH has been shown to enhance cathepsin L (CTSL)-mediated cleavage of endogenously expressed C3, indicating a potential pathway to explore (Martin et al., 2016). Additionally, future studies should elucidate the specific intracellular functions and downstream signalling pathways of the C3 forms, as C3 has been implicated in non-canonical roles, such as in nutrient and oxygen metabolism and cellular homeostasis (Arbore et al., 2017). Furthermore, given the strong association between AMD risk and polymorphisms in complement genes, as described in **3.2**, it would be essential to investigate genotype-dependent variations in intracellular complement production, secretion, and C3 activation in iPSC-RPE models with different risk single nucleotide polymorphisms. Additionally, *in vitro* RPE cell models have limitations, as they cannot account for tissue-dependent responses or the systemic complement cascade’s influence on cellular complement production and C3 processing. These factors highlight the need for more comprehensive studies to understand the role of local cellular complement activity in AMD pathogenesis.

### 3.4. Resistance of mature iPSC-RPE cells to TGF-β1-mediated stress and its impact on complement secretion

Beyond analysing cell phenotype and C3 processing in our maturation- and MET-based RPE model, we aimed to investigate these mechanisms in an EMT-based model. Our idea was to revert mature ARPE-19 and iPSC-RPE cells into EMT and a more immature phenotype. EMT and TGF-β signalling activation have been discussed as pathomechanisms associated with AMD progression (Little, Llorián-Salvador, Tang, Du, Marry, et al., 2020; Little, Llorián-Salvador, Tang, Du, O’Shaughnessy, et al., 2020; Radeke et al., 2015). For example, transcriptomic profiling showed a significant enrichment of wound response genes, including transforming growth factor 2 (*TGFB2*), in AMD phenotypes (Newman et al., 2012).

TGF-β1 treatment of RPE cells is a well-established approach to inducing EMT, as multiple studies have shown (S.-J. Kim et al., 2020; Lee et al., 2013; M. Li et al., 2021; Y. Li et al., 2015; Ma et al., 2023; Su et al., 2021; M. Wang et al., 2022; Wei et al., 2018). However, these studies used ARPE-19 cells or human primary RPE cells in an immature state, maintained in a similar medium as our immature ARPE-19 cells, and cultured the cells only until sub-confluency before initiating TGF-β1 treatment, which does not resemble *in vivo*-like conditions.

To induce an EMT-like transition in our mature ARPE-19 and iPSC-RPE cells, we followed the approach of Boles et al. with 10 ng/mL TGF-β1 applied for five consecutive treatment days, expecting that the mature cells would require an extended period of treatment to undergo EMT (Boles et al., 2020).

In our study, mature ARPE-19 cells displayed strong expression of TβRI, suggesting a higher sensitivity to TGF-β1 treatment; mature iPSC-RPE cells exhibited weaker TβRI signals, indicating potential resistance to the treatment (**Figure 6E**, **Figure 6 - video 2**). This hypothesis was supported by the changes observed: mature ARPE-19 cells showed alterations in TJP1 distribution and *CDH1* and *BEST1* transcript expression following TGF-β1 treatment, whereas mature iPSC-RPE cells exhibited minimal changes (**Figure 7A, B, C**, **Figure 7 - video 2, Figure 7 - video 3**).

However, neither ARPE-19 nor iPSC-RPE cells in our study fully transitioned into an EMT following treatment, suggesting they were not comparable to immature RPE cells used in previous studies (S.-J. Kim et al., 2020; Lee et al., 2013; M. Li et al., 2021; Y. Li et al., 2015; Ma et al., 2023; Su et al., 2021; M. Wang et al., 2022; Wei et al., 2018). Rather, the TGF-β1 treatment induced a stressed state in these cells, characterised by subtle morphological changes. This may be due to the high degree of maturation prior to treatment, which could indicate the need for a longer or more concentrated TGF-β1 exposure or additional external disturbances such as monolayer injury to induce EMT. In support of this, Tamiya et al. demonstrated that TGF-β2 treatment was insufficient to induce EMT in primary porcine RPE cells that maintained intact cell-cell junctions, with EMT only initiated upon disruption of the tight junction monolayer (Tamiya et al., 2010).

Despite the differences between treated ARPE-19 and iPSC-RPE cells in morphological responses, the effects of TGF-β1 on complement secretion were mostly consistent across both ARPE-19 and iPSC-RPE cells. TGF-β1 treatment significantly reduced the apical secretion of C3, C3(H_2_O), iC3b, FH/FHL-1, and FI, while the release of activated C3a remained unchanged (**Figure 7E**-**H**, **Figure 7 - supplement 1F**, **H**). However, TGF-β1 did not alter the cellular cleavage pattern of C3 (**Figure 7I**-**K**, **Figure 7 - video 4**).

Few studies have analysed complement components in the context of TGF-β1 treatment in RPE cells. Boles et al. reported no significant changes in *C3*, *CFH*, or *CFI* transcript expression following treatment of RPESC-RPE cells with 10 ng/mL TGF-β1 alone (Boles et al., 2020). We analysed the dataset from Boles et al. using GEO2R and observed an increase in *C3* transcript levels with combined treatment of 10 ng/mL TNF-α/TGF-β1 (data accessible at NCBI GEO database, accession GSE128143) (**Appendix 2B**). In contrast, our findings revealed a decrease in *C3* levels in ARPE-19 cells treated with TGF-β1 alone, while iPSC-RPE cells displayed no significant changes. Additionally, *CFH* and *CFI* transcript levels decreased in RPESC-RPE cells after TNF-α/TGF-β1-induced EMT, which aligns with our findings in treated ARPE-19 cells, while no differences were observed in treated iPSC-RPE cells (**Appendix 2B**). Furthermore, Li et al. found reduced *C3* transcript expression in ARPE-19 cells treated with 10 ng/mL of TGF-β1 for 48 hours, affirming our observation in treated ARPE-19 cells (Y. Li et al., 2015).

These findings suggest that TGF-β1-induced EMT and complement activation are interconnected mechanisms involved in retinal fibrosis, both of which play critical roles in AMD pathogenesis. The TGF-β pathway has yet to be specifically explored in relation to HRF. Given that HRF development has been associated with EMT processes as described in **3.1,** it represents a promising area for future investigation.

In summary, our study, along with others, demonstrates that both healthy and stressed RPE cells produce multiple components of the complement cascade, activate C3 intracellularly, and secrete pre-activated C3 forms, including C3a, C3(H_2_O), and iC3b. These fragments can exert their biological functions without interference from current anti-C3 therapies used in AMD treatment to slow retinal degeneration. Specifically, C3a binds to the C3a receptor, inducing cell activation, while iC3b engages complement receptor 3, promoting phagocytosis. Moreover, our findings suggest that C3 and its activation products may play intracellular roles, potentially influencing processes like MET and EMT, which are not targeted by extracellular C3 inhibitors. Additionally, anti-C3 therapies, such as Pegcetacoplan, are delivered intravitreally, primarily affecting apical C3. However, basal C3 or C3(H_2_O) may remain largely unaffected. These limitations could explain why anti-C3 antibodies slow the progression of GA, but do not fully stop the disease.

## 4. Materials and Methods

### 4.1. Cell culture

ARPE-19 cells (ATCC, Manassas, VA, USA; #CRL-2302) at passage 27 were used for all experiments. The cells were authenticated using short tandem repeat (STR) profiling by the ATCC Cell Line Authentication Service, following ISO 9001 and ISO/IEC 17025 quality standards. The STR profile matched the ARPE-19 reference profile in the ATCC STR database (CODIS 13 loci). Immature ARPE-19 cells were maintained in Dulbecco′s Modified Eagle′s Medium/Nutrient Mixture F-12 Ham (Merck KGaA, Darmstadt, Germany) supplemented with 10% fetal bovine serum (Thermo Fisher Scientific, Waltham, MA, USA), 2 mM L-glutamine (Thermo Fisher Scientific) and penicillin-streptomycin (Thermo Fisher Scientific). Mature ARPE-19 cells were cultured in MEM-Nic medium according to the protocol of Hazim et al. (Hazim et al., 2019). Cells were initially thawed in T175 culture flasks until approximately 80% confluency before being trypsinised for 4 minutes in a trypsin-ethylenediaminetetraacetic acid (EDTA)-solution 1X (Merck KGaA). A total of 1.75 x 10^5^ cells per insert was then transferred to Thincert cell culture inserts with 0.4 µm pores (Greiner Bio-One, Kremsmünster, Austria) pre-coated with 16 µg/mL laminin (Merck KGaA). ARPE-19 cells were further cultivated for 2 weeks.

iCell iPSC-RPE cells (Fujifilm Cellular Dynamics, Inc., Madison, WI, USA; #R1113) were cultivated in accordance with the manufacturer’s instructions. Following thawing, cells were seeded onto three 6-wells coated with 50 µg/cm^2^ Matrigel® matrix (Corning Inc., Corning, NY, USA). After one week, cells were detached using 1 mM EDTA in phosphate-buffered saline (PBS) for 10 minutes at room temperature, followed by incubation with TrypLE Select Enzyme 1X (Thermo Fisher Scientific) for 10 minutes at 37 C. Cells were then split at a 1:4 ratio and seeded into 6-well plates. After another week, they were trypsinised, and a total of 1.75 x 10^5^ cells per insert was transferred onto 50 µg/cm^2^ Matrigel-coated Thincert cell culture inserts with 0.4 µm pores (Greiner Bio-One). To obtain immature and mature cells, cultivation periods of 1 and 6 weeks, respectively, were chosen.

ARPE-19 and iPSC-RPE cells were cultured at 37 C in a humidified atmosphere with 5% CO₂, with medium changes every 72 hours or less. Prior to harvesting the cells, apical and basal supernatants were collected following a 72-hour incubation without medium change. Data for immature and mature RPE cells presented in this study were pooled from two to three independent cultivations.

### 4.2. TER measurement

At the end of cell cultivation, TER and cell capacitance were measured using the CellZscope3 (nanoAnalytics GmbH, Münster, Germany). Cells on Thincert cell culture inserts and medium were added to the device, and measurements were started immediately. To ensure cellular adaptation, a measurement point after 10 hours was selected for data analysis. Laminin- and Matrigel-coated cell inserts without cells were measured as blanks for ARPE-19 and iPSC-RPE cells, respectively; the values obtained were subtracted in the subsequent analysis.

### 4.3. TGF-ꞵ1 treatment

Mature ARPE-19 and iPSC-RPE cells were treated with 10 ng/mL recombinant human TGF-β1 (R&D Systems, Minneapolis, MN, USA) or vehicle control in serum-free conditions. TGF-β1 was added to both the apical and basal media, with daily medium changes for five consecutive days. Cells were left in the medium of the final treatment day on days 6 and 7, and samples were collected on the eighth day after treatment initiation. Data regarding TGF-β1 treatment of RPE cells were pooled from two independent treatment replicates.

### 4.4. Mycoplasma contamination analysis

Mycoplasma contamination in ARPE-19 and iPSC-RPE cells was assessed using the LookOut® Mycoplasma PCR Detection Kit (Merck KGaA) according to the manufacturer’s instructions. No mycoplasma contamination was detected in the cell cultures.

### 4.5. Single nucleotide polymorphism analysis

DNA was isolated using the DNeasy Blood & Tissue Kit (Qiagen, Hilden, Germany), followed by the analysis of single nucleotide polymorphisms using the TaqMan™ SNP Genotyping Assay (Thermo Fisher Scientific). The analysed single nucleotide polymorphisms are listed in **Appendix 3**.

### 4.6. Quantitative gene expression analysis

RNA was extracted using the NucleoSpin for RNA purification kit, following the manufacturer’s instructions (MACHEREY-NAGEL, Düren, Germany), followed by cDNA synthesis utilising the QuantiTect reverse transcription kit, according to the manufacturer’s instructions (Qiagen). qRT-PCR was conducted with the Power SYBR™ Green PCR Master Mix (Thermo Fisher Scientific) on the QuantStudio™ 5 Real-Time PCR System (Thermo Fisher Scientific). The amplification protocol included 40 cycles, with a 1-minute annealing step at 60 C. Primer sequences are listed in **Appendix 3**. The *CFH* primer was obtained from Qiagen (#QT00001624). Data analysis was based on the 2^(-ΔΔCT)^ method with log2-transformed scores. Transcript expression levels were normalised to the reference gene glyceraldehyde-3-phosphate dehydrogenase (*GAPDH*). Expression values were represented as multiples relative to those from immature cells. For TGF-ꞵ1 treatment, values were represented as multiples relative to the corresponding values from control cells.

### 4.7. Immunocytochemistry

Cells were fixed in 4% paraformaldehyde at 4 C for 20 minutes, followed by three washes with PBS. Permeabilisation was carried out in PBS with Tween20 (PBS-T) for 45 minutes, and blocking was performed using 3% bovine serum albumin (BSA) in PBS-T. Primary antibodies (listed in **Appendix 3**) were diluted in blocking solution and incubated with the cells overnight at 4 C. After three washing steps, secondary antibodies were applied and incubated in 3% BSA in PBS at room temperature for 45 minutes. The antibodies are listed in **Appendix 3**. Nuclei were stained with Hoechst 33342 solution (Thermo Fisher Scientific). Imaging was performed using the Fluoview Fv10i confocal microscope (Olympus, Shinjuku, Japan) at 60 x magnification.

Image processing and analysis were carried out using Fiji (ImageJ) version 1.54f as described below (Schindelin et al., 2012). A custom macro was developed to standardise background subtraction and generate summary images for each channel from recorded stacks using the Z-project function, with maximum intensity projection. Overlays were created and saved as Audio Video Interleave (AVI) files at 0.5 frames per second, which were subsequently compiled into MPEG-4 Part 14 (MP4) file collages using Canva (Canva, Sydney, Australia).

The brightness and contrast of nuclear staining were adjusted when the staining was very intense, to prevent it from overshadowing the focused staining. To enhance visibility, the brightness of BEST1 was increased by 100% in all images of all groups. Furthermore, the brightness of GT335 in the co-staining with TJP1 was increased by 100% in all images of all groups, facilitating differentiation from nuclear and TJP1 staining.

Cell density (cells/cm²) and the cross-sectional areas of cells and nuclei were calculated using a custom macro. Briefly, images were converted to 16-bit grayscale, filtered using a median filter, and subjected to auto thresholding to create binary images. Particle analysis was subsequently performed on the binary images. Neighbour analysis was performed manually by counting.

Fluorescence intensity analysis was performed by measuring the mean fluorescence intensity of the images and normalising the values to the number of cell nuclei present in each image. GT335 and ARL13B lengths were measured using the CiliaQ plugins for Fiji (Hansen et al., 2021). Recorded stacks were prepared with CiliaQ Preparator version 0.1.2 using default settings and the segmentation method *RenyiEntropy*, followed by editing with CiliaQ Editor version 0.0.3. Analysis of the edited stacks was performed using CiliaQ version 0.1.7, applying default settings and a minimum cilium size threshold of 1 voxel. For GT335 length measurements, values of 0 were excluded, and for ARL13B length, values below 1 µm were removed to eliminate nonspecific inclusion of ARL13B co-localisation with cell nuclei.

### 4.8. SDS-PAGE and Western blot

Protein from cell lysates was extracted using M-PER™ mammalian protein extraction lysis buffer, supplemented with 1X Halt™ Protease Inhibitor Cocktail, and 1X EDTA (Thermo Fisher Scientific). Lysate samples were concentrated by a factor of two using Amicon ultra centrifugal filter units with a 3 kDa molecular weight cut-off (Merck KGaA). Protein concentration was quantified using the Pierce™ bicinchoninic acid assay (Thermo Fisher Scientific).

Samples were prepared with reducing ROTI®Load 1 (Carl Roth, Karlsruhe, Germany) and separated by SDS-PAGE on TGX Stain-Free™ FastCast™ 12% gels, using Bio-Rad MiniPROTEAN® (Bio-Rad Laboratories, Inc., Hercules, CA, USA) equipment. 40 µg of protein were loaded per sample. Stain-free gel technology was used to visualise total protein content, as a loading control. Representative total protein staining of RPE cell lysates, used for the detection of BEST1, is shown in **Figure 1 - supplement 1**. Proteins were transferred onto polyvinylidene difluoride membranes by wet blotting for 100 minutes at 80 V. Membranes were blocked for two hours at room temperature in a blocking buffer consisting of 5% milk in Tris-buffered saline with Tween20 (TBS-T). Primary antibodies were diluted in blocking buffer and incubated overnight at 4 C, followed by three 10-minute washes in TBS-T. Secondary antibody incubation was then performed in blocking buffer at room temperature for 2 hours. After three additional wash steps, protein signals were detected using the WesternBright Sirius chemiluminescent substrate (Biozym Scientific GmbH, Hessisch Oldendorf, Germany) and visualised using the ChemiDoc MP Imaging System (Bio-Rad Laboratories, Inc.). A list of antibodies and purified proteins is provided in **Appendix 3**.

### 4.9. Immunoprecipitation

Invitrogen™ M-280 Tosyl-activated Dynabeads™ (Thermo Fisher Scientific) were vortex-mixed and washed with 1 mL of buffer A (0.1 M borate buffer). The DynaMag™-2 Magnet (Thermo Scientific) was applied for 2 minutes to remove the supernatant, followed by an additional wash with buffer A. To couple the beads, 50 µg of the C3 β chain antibody was added to 6.25 mg of beads in 100 µL buffer A and 100 µL buffer B (3 M ammonium sulphate in buffer A). The mixture was incubated at room temperature for 20 hours. Buffer C (0.88% (w/v) NaCl and 0.5% (w/v) BSA) was then added and mixed at room temperature for 1 hour. After three washes with buffer D (0.88% (w/v) NaCl and 0.1% (w/v) BSA), a solution of 20 mg/mL ligand-coupled beads was prepared.

For immunoprecipitation, 50 µL of ligand-coupled beads were added to ARPE-19 and iPSC-RPE cell lysates containing 50 µg of protein and incubated for 1 hour at room temperature, followed by incubation overnight at 4 C. Samples were then washed twice with PBS-T, once with PBS, and the supernatant was discarded. The ligand-coupled beads containing the precipitated target protein were resuspended in 50 µL of reducing ROTI®Load 1, and samples were denatured at 95 C for 10 minutes. Beads were magnetically separated from the protein mixture. SDS-PAGE and Western blot were performed as described in **4.8**.

### 4.10. Multiplex immunoassay

Parallel multiplex immunoassays were performed using the Luminex® xMAP® technology with the MILLIPLEX® Human Complement panels, Expanded Panel 1 and Panel 2 according to the manufacturer’s instructions (Merck KGaA). A total of 50 µL of undiluted cell supernatants, along with respective medium blanks, were incubated with 25 µL of magnetic beads at 4 C overnight. Following three washing steps, 25 µL of antibody detection mix were added and incubated at room temperature for 1 hour. After a 30-minute incubation with streptavidin-R-Phycoerythrin, samples were washed three times. Measurements were recorded as median fluorescence intensity (MFI) values using the Bio-Plex 200 system (Bio-Rad Laboratories, Inc.). MFI values below 100 were considered not biologically relevant. Quality controls were included in the measurements, and medium blanks were subtracted during data analysis.

The C3 and iC3b beads of the multiplex kits were validated using purified proteins, human sera and depleted sera, as detailed in **Appendix 3.** The validation of C4, C4B, and FH/FHL-1 beads was previously reported by Schikora et al. (2025, in proof).

### 4.11. ELISA

Cell supernatants were analysed using the human TGF-β1 enzyme-linked immunosorbent assay (ELISA) kit (Merck KGaA). Samples and blanks were first activated by incubation with 1 N HCl for 10 minutes, followed by neutralisation with 1.2 N NaOH/0.5 M Hepes (pH 7.0). A total of 100 µL of each sample was then added to the wells and incubated for 2.5 hours. After four wash steps, 100 µL of detection antibody was added and incubated for 1 hour. The wells were washed again before the addition of 100 µL of streptavidin solution, which was incubated for 45 minutes, followed by another wash cycle. Finally, 100 µL of TMB reagent was added to each well and incubated in the dark for 30 minutes. The reaction was stopped by adding 50 µL of stop solution.

To analyse C3a and C3(H_2_O), cell supernatants were further concentrated by a factor of five using Amicon ultra centrifugal filter units with a 3 kDa molecular weight cut-off (Merck KGaA). To measure C3a, the human C3a ELISA kit (Hycult Biotech, Uden, Netherlands) was used. A total of 100 µL of each 5-fold concentrated sample and corresponding blanks were incubated in the assay wells for 1 hour. Following this, four washing steps were conducted before the addition of 100 µL of diluted tracer, which was incubated for 1 hour. After another washing sequence, 100 µL of streptavidin solution was added and incubated for 1 hour, followed by a final washing cycle. To develop the assay, 100 µL of TMB reagent was added to each well and incubated in the dark for 30 minutes. The reaction was terminated by the addition of 100 µL of stop solution.

The C3(H_2_O) ELISA was performed following the protocol described by Elvington et al. (Elvington et al., 2019). Maxisorp 96-well plates were coated with 100 µL of 2 µg/mL 3E7 C3 antibody and incubated overnight at 4 C. After three washing steps, the plates were blocked with 4% BSA in PBS-T for 1 hour. Subsequently, 50 µL of concentrated samples and blanks were incubated for 1 hour. The wells were then incubated with biotin-labelled C3a antibody for 1 hour, followed by a washing cycle and a 30-minute incubation with streptavidin. After another wash sequence, 100 µL of TMB reagent was added and incubated for up to 30 minutes in the dark. The reaction was terminated by adding 100 µL of hydrochloric acid.

Unless otherwise indicated, all ELISA steps were conducted at room temperature and absorbance was measured as optical density (OD) at 450 nm using a FLUOstar Optima reader (BMG LABTECH, Ortenberg, Germany). Medium blanks were subtracted during further analysis. Antibodies and purified proteins are detailed in **Appendix 3**.

### 4.12. Statistical analysis

Data were organised using Microsoft Excel version 2108 (Microsoft Corporation, Redmond, WA, USA) and figures were prepared using Affinity Designer version 1.9.1.979 (Serif, West Bridgford, UK). Graphical and statistical analyses were conducted using R version 4.2.1 (R Core Team, 2021), along with the R packages tidyverse (Wickham et al., 2019), pastecs (Grosjean & Ibanez, 2018), car (Fox & Weisberg, 2019), ggsignif (Ahlmann-Eltze & Indrajeet, 2021), RColorBrewer (Neuwirth, 2022), gmodels (Warner et al., 2022), and polycor (Fox, 2022). Normality of data was checked with histograms, Q-Q plots, kurtosis and skew values, as well as Shapiro-Wilk test. Homogeneity of variance was assessed with Levene’s test. Parametric data were analysed using an independent-means *t*-test, with mean (M) and standard deviation (SD) displayed in the graphs. For non-parametric data, the Wilcoxon’s rank-sum test was utilised and median (Mdn), with first (Q1) and third (Q3) quartiles, are shown in graphs. All tests were two-tailed and an alpha level of .050 (*P* < .050) defined statistical significance. The NCBI GEO tool GEO2R was used to analyse of transcriptomic datasets from other studies (Barrett et al., 2012).

## label>5. List of abbreviations

AMD: (age-related macular degeneration)
ARL13B: (ADP-ribosylation factor-like protein 13B)
ARPE-19: (adult retinal pigment epithelial cell line)
AVI: (Audio Video Interleave)
BEST1: (bestrophin 1)
BSA: (bovine serum albumin)
C1q: (complement 1q)
C2: (complement 2)
C3: (complement 3)
C3a: (complement 3a)
C3aR: (C3a receptor)
C3b: (complement 3b)
C3c: (complement 3c)
C3d: (complement 3d)
C3dg: (complement 3dg)
C3f: (complement 3f)
C3g: (complement 3g)
C3(H_2_O): (hydrolysed C3)
C4: (complement 4)
C4B: (complement 4B)
C5: (complement 5)
C5a: (complement 5a)
C9: (complement 9)
CDH1: (cadherin 1)
CFB/FB: (complement factor B)
CFD/FD: (complement factor D)
CFH/FH: (complement factor H)
CFI/FI: (complement factor I)
CTSL: (cathepsin L)
dpl: (depleted)
ECM: (extracellular matrix)
EDTA: (ethylenediaminetetraacetic acid)
EFEMP1: (EGF containing fibulin-extracellular matrix protein 1)
ELISA: (enzyme-linked immunosorbent assay)
EMT: (epithelial-to-mesenchymal transition)
F-actin: (filamentous actin)
FHL-1: (factor H-like protein 1)
FHR-1: (factor H-related protein 1)
FHR-2: (factor H-related protein 2)
FHR-3: (factor H-related protein 3)
FHR-4: (factor H-related protein 4)
FHR-5: (factor H-related protein 5)
GA: (geographic atrophy)
GAPDH: (glyceraldehyde-3-phosphate dehydrogenase)
GEO: (Gene Expression Omnibus)
GT335: (polyglutamylated tubulin)
HRF: (hyperreflective foci)
iC3b: (inactivated complement 3b)
im: (immature)
iPSC-RPE: (induced pluripotent stem cell-derived RPE)
IQR: (interquartile range)
kDa: (kilodalton)
LAMP2: (lysosome-associated membrane protein 2)
M: (mean)
ma: (mature)
MBL: (mannan-binding lectin)
Mdn: (median)
MERTK: (MER proto-oncogene, tyrosine kinase)
MET: (mesenchymal-to-epithelial transition)
MFI: (median fluorescence intensity)
mL: (millilitre)
mM: (millimolar)
MP4: (MPEG-4 Part 14)
NCBI: (National Center for Biotechnology Information)
nm: (nanometre)
NRF2: (nuclear factor-erythroid 2-related factor 2)
OD: (optical density)
PBS: (phosphate-buffered saline)
PBS-T: (PBS with Tween20)
PMEL: (premelanosome protein)
PPI: (pixel per square inch)
Q_1_: (first quartile)
Q_3_: (third quartile)
qRT-PCR: (quantitative real time polymerase chain reaction)
RLBP1: (retinaldehyde binding protein 1)
RPE: (retinal pigment epithelium)
RPE65: (retinoid isomerohydrolase)
RPESC-RPE: (adult human RPE stem cell-derived RPE)
SD: (standard deviation)
SDS: (sodium dodecyl sulphate)
SDS-PAGE: (SDS polyacrylamide gel electrophoresis)
SNAI1: (snail family transcriptional repressor 1)
TBS-T: (Tris-buffered saline with Tween20)
TJP1: (tight junction protein zonula occludens 1)
TGF-β/TGFB: (transforming growth factor-beta)
TGF-β1/TGFB1: (transforming growth factor-beta 1)
TGF-β2/TGFB2: (transforming growth factor beta 2)
TβRI/TGFBR1: (transforming growth factor-beta receptor 1)
TβRII: (transforming growth factor-beta receptor 2)
TER: (transepithelial resistance)
TNF-α: (tumour necrosis factor alpha)
VIM: (vimentin)
µF/cm^2^: (micro farad per square centimetre)
µg: (microgram)
µg/cm²: (microgram per square centimetre)
µg/mL: (microgram per millilitre)
µL: (microlitre)
µm: (micrometre)
Ω·cm^2^: (ohm square centimetre)

## 6. Declarations

### 6.1. Ethics approval and consent to participate

Not applicable.

### 6.2. Consent for publication

Not applicable.

### 6.3. Data availability

All data generated or analysed during this study are included in the manuscript and its supplementary materials. Analytical data, images, videos, and Western Blot results have been deposited in the University Marburg’s open-access repository (data_UMR) as part of the data repository for retinal pigment epithelial cell maturation (https://data.unimarburg.de/handle/dataumr/628).

## 6.4. Acknowledgements

The authors would like to express their sincere gratitude to Knut Stieger and Maria Weller, Department of Ophthalmology, Justus-Liebig-University Giessen, Giessen, Germany, for their support and for providing access to the confocal microscope, and to Sanquin Research, Amsterdam, Netherlands, and Mihály Józsi, Department of Immunology, ELTE Eötvös Loránd University, Budapest, Hungary, for generously providing materials.

## Rich media files

**Figure 1 - video 1 (Figure 1 – video 1.mp4): Immature cells show a mesenchymal F-actin cytoskeleton, while mature cells show epithelial-like structures.**

The video shows F-actin in an apical to basal direction in immature and mature ARPE-19 and iPSC-RPE cells. Immature ARPE-19 cells show a distinct mesenchymal F-actin cytoskeleton featuring stress fibres and elongated fibres traversing the whole cell, while mature ARPE-19 cells show a distribution more typical for epithelial cells, alongside few stress fibres. Both immature and mature iPSC-RPE cells exhibit an epithelial circumferential F-actin distribution, although immature iPSC-RPE cells display more stress fibres. Furthermore, in mature iPSC-RPE cells F-actin is only located apically above the nuclear layer. Scale bar = 50 µm.

**Figure 1 - video 2 (Figure 1 – video 2.mp4): Mesenchymal VIM signals are detectable in immature cells.**

The video shows VIM in an apical to basal direction in immature and mature ARPE-19 and iPSC-RPE cells. Immature ARPE-19 and iPSC-RPE cells show prominent signals for VIM located in proximity to the nucleus. Mature iPSC-RPE cells show a weak circumferential VIM localisation. Scale bar = 50 µm.

**Figure 1 - video 3 (Figure 1 – video 3.mp4): TJP1 shows an apical and circumferential localisation in mature ARPE-19 and iPSC-RPE cells.**

The video shows TJP1 in an apical to basal direction in immature and mature ARPE-19 and iPSC-RPE cells. Mature cells show a more apical circumferential location of TJP1, while TJP1 in immature cells is located at the level of the nuclei. Scale bar = 50 µm.

**Figure 1 - video 4 (Figure 1 – video 4.mp4): Basal BEST1 localisation is displayed in mature iPSC-RPE cells.**

The video shows BEST1 in an apical to basal direction in immature and mature ARPE-19 and iPSC-RPE cells. Only mature iPSC-RPE cells exhibit BEST1 signals circumferentially that are more located at the basal site of the cell at the level of the nuclei. Scale bar = 50 µm.

**Figure 2 - video 1 (Figure 2 – video 1.mp4): Apical GT335 and ARL13B show co-localisation.**

The video shows GT335 (cyan) and ARL13B (magenta) in cilia in an apical to basal direction in immature and mature ARPE-19 and iPSC-RPE cells. Co-localisation is displayed in white. Scale bar = 50 µm.

**Figure 2 - video 2 (Figure 2 – video 2.mp4): Apical GT335 signals are reduced in mature cells.**

Scale bar = 50 µm. The video shows GT335 in cilia and TJP1 in an apical to basal direction in immature and mature ARPE-19 and iPSC-RPE cells. GT335 signals are reduced and appear as punctate dots in mature cells, while immature cells display more linear patterns. GT335 is located apically in the cells. Scale bar = 50 µm.

**Figure 2 - video 3 (Figure 2 – video 3.mp4): Apical ARL13B shows decreased signals in mature cells.**

The video shows ARL13B in cilia in an apical to basal direction in immature and mature ARPE-19 and iPSC-RPE cells. ARL13B is located apically in the cells and appears as punctate dots, that are reduced in mature cells. Furthermore, immature ARPE-19 and iPSC-RPE, as well as mature ARPE-19 cells show ARL13B co-localised with the cell nucleus.

**Figure 3 - video 1 (Figure 3 – video 1.mp4): FH family proteins are located intracellularly with different intensities in immature and mature cells.**

The video shows the staining of the FH family in an apical to basal direction in immature and mature ARPE-19 and iPSC-RPE cells. For antibody specificity see **Figure 3 - supplement 2**. Increased FH protein family signal intensity is detectable in mature ARPE-19 cells compared with immature cells, while iPSC-RPE cells show the opposite trend of increased signals in immature cells. Scale bar = 50 µm.

**Figure 3 - video 2 (Figure 3 – video 2.mp4): Immature ARPE-19 cells show a specific FH staining.**

The video shows the specific staining of FH in an apical to basal direction in immature and mature ARPE-19 and iPSC-RPE cells. It is detected with an antibody that is specific for FH domain 16. For antibody specificity see **Figure 3 - supplement 3**. Immature ARPE-19 cells display FH intracellularly, while the rest does not exhibit FH signals. Scale bar = 50 µm.

**Figure 3 - video 3 (Figure 3 – video 3.mp4): FI shows an apical location in mature iPSC-RPE cells.**

The video shows FI in an apical to basal direction in immature and mature ARPE-19 and iPSC-RPE cells. No uniform trend of FI signal distribution is detectable in ARPE-19 and iPSC-RPE cells. Mature iPSC-RPE cells show a FI distribution that is located in the apical part of the cells. Scale bar = 50 µm.

**Figure 5 - video 1 (Figure 5 – video 1.mp4): C3 shows perinuclear distribution and apical localisation in mature iPSC-RPE cells.**

The video shows C3 staining by antibody 1 in an apical to basal direction in immature and mature ARPE-19 and iPSC-RPE cells. C3 is accumulated intracellularly near the nucleus, with stronger overall signals in mature cells. Mature iPSC-RPE show a predominantly apical C3 localisation and further circumferential signal distribution. Scale bar = 50 µm.

**Figure 5 - video 2 (Figure 5 – video 2.mp4): Mature cells show localised C3 signals, while immature cells exhibit diffuse distribution.**

The video shows C3 staining by antibody 2 in an apical to basal direction in immature and mature ARPE-19 and iPSC-RPE cells. C3 signals in immature cells are diffuse and more widespread throughout the cell, while mature cells show more localised signals. Mature iPSC-RPE cells show a predominantly apical C3 localisation and further circumferential signal distribution. Scale bar = 50 µm.

**Figure 6 - video 1 (Figure 6 – video 1.mp4): Distribution of TGF-β1 varies in ARPE-19 and iPSC-RPE cells.**

The video shows TGF-β1 in an apical to basal direction in immature and mature ARPE-19 and iPSC-RPE cells. TGF-β1 shows diffuse distribution in ARPE-19 and immature iPSC-RPE cells, while mature iPSC-RPE cells display an apical and circumferential localisation. Scale bar = 50 µm.

**Figure 6 - video 2 (Figure 6 – video 2.mp4): Strong TβRI accumulations in mature ARPE-19 cells.**

The video shows TβRI in an apical to basal direction in immature and mature ARPE-19 and iPSC-RPE cells. TβRI shows fibre-like structures in immature ARPE-19 cells and strong punctate distributions in mature ARPE-19 cells. Immature iPSC-RPE cells exhibit diffuse staining throughout the cell, while mature iPSC-RPE cells show signals at the apical cell border. Scale bar = 50 µm.

**Figure 6 - video 3 (Figure 6 – video 3.mp4): Co-localisation of TGF-β1 and TβRI in ARPE- 19 and iPSC-RPE cells.**

The video shows the co-staining of TGF-β1 (cyan) and TβRI (magenta) from **Figure 6 - video 1** and **Figure 6 - video 2** in an apical to basal direction in immature and mature ARPE-19 and iPSC-RPE cells. Co-localisation is displayed in white. Scale bar = 50 µm.

**Figure 7 - video 1 (Figure 7 – video 1.mp4): TGF-β1-treated cells exhibit more diffuse TGF-β1 signals, while control cells show stronger signals.**

The video shows TGF-β1 in an apical to basal direction in control and TGF-β1-treated ARPE-19 and iPSC-RPE cells. TGF-β1 is detectable in a more diffuse pattern in TGF-β1-treated cells, while control cells show more focussed and stronger signals. Scale bar = 50 µm.

**Figure 7 - video 2 (Figure 7 – video 2.mp4): ARPE-19 cells display stress fibres, while treated iPSC-RPE cells show no strict apical F-actin localisation.**

The video shows F-actin in an apical to basal direction in TGF-β1-treated and control ARPE-19 and iPSC-RPE cells. Stress fibres are visible in control and TGF-β1-treated ARPE-19 cells, while no stress fibres are detectable in iPSC-RPE cells. The strict apical localisation of F-actin is only visible in control iPSC-RPE cells. Scale bar = 50 µm.

**Figure 7 - video 3 (Figure 7 – video 3.mp4): TGF-β1-treated ARPE-19 cells display thickened TJP1 structures, while treated iPSC-RPE cells show no strict apical TJP1 location.**

The video shows TJP1 in an apical to basal direction in TGF-β1-treated and control ARPE-19 and iPSC-RPE cells. TJP1 structures of TGF-β1-treated ARPE-19 cells are thickened compared with control cells. The strict apical localisation of TJP1 is only visible in control iPSC-RPE cells. Scale bar = 50 µm.

**Figure 7 - video 4 (Figure 7 – video 4.mp4): TGF-β1-treated ARPE-19 cells show stronger C3 signals, while signals are located apically in control iPSC-RPE cells.**

The video shows C3 staining by antibody 1 in an apical to basal direction in control and TGF-β1-treated ARPE-19 and iPSC-RPE cells. TGF-β1-treated ARPE-19 cells display slightly stronger C3 accumulations near the nucleus. Control iPSC-RPE cells exhibit a particular apical C3 location above the nuclear layer. Scale bar = 50 µm.

## Figure supplements

**Figure 1 - supplement 1 (Figure 1 – supplement 1. jpg): Fluorescence intensity analysis of cell phenotype stainings demonstrate differences for F-actin and TJP1.**

**(A)** Fluorescence intensity of F-actin is reduced in mature ARPE-19 cells compared with immature cells, with no difference observed in iPSC-RPE cells (*n* = 3). **(B)** Fluorescence intensity of VIM shows no significant difference between conditions (*n* = 3). **(C)** Fluorescence intensity of TJP1 is decreased in mature iPSC-RPE cells compared with immature cells, while no difference is observed in ARPE-19 cells (*n* = 3). **(D)** Fluorescence intensity of BEST1 shows no significant difference between conditions (*n* = 3). Non-parametric data (underlined) were analysed using the Wilcoxon test (Mdn and IQR); parametric data with independent-means *t*-test (M and SD). Tests were two-tailed, and statistical significance was defined as *P* < .050.

**Figure 1 - supplement 2 (Figure 1 – supplement 2. jpg): BEST1 is expressed exclusively in mature iPSC-RPE cells, with representative total protein staining of cell lysates.**

**(A)** Western blot analysis of cell lysates with immature (Im) and mature (Ma) cell lysates showed a distinct 70 kDa band for BEST1, present exclusively in mature iPSC-RPE cells. **(B)** Representative total protein staining of RPE cell lysates, used to confirm protein loading for BEST1 detection. M = marker.

**Figure 1 - supplement 3 (Figure 1 – supplement 3. jpg): Representative images show basis for image analysis of ARPE-19 and iPSC-RPE cell properties.**

**(A)** The calculations for **Figure 1J** and **K** are based on the cell area determined by the immunofluorescent localisation of TJP1, representing data from *n* = 3 images. **(B)** The neighbour analysis for **Figure 1L** was based on the TJP1 staining, with data derived from *n* = 2 images. **(C)** The calculation of nuclear area for **Figure 1N** was based on Hoechst nuclei staining, representing data from *n* = 10 images. Scale bar = 50 µm.

**Figure 3 - supplement 1 (Figure 3 – supplement 1. jpg): Immature and mature cells show different apical and basal complement secretion.**

The apical and basal secretion levels of C1q, C2, C4, C4B, C5, C5a, C9, MBL, FD, and properdin were assessed (*n* = 5). Except for C4B, all components were secreted at notably low levels. MFI values below 100 were considered not biologically relevant. Components with undetectable secretion, indicated by missing *P* values, exhibited exclusively MFI values of 0. Data represent pooled results from two independent culture replicates. Non-parametric data (underlined) were analysed using the Wilcoxon test (Mdn and IQR); parametric data with independent-means *t*-test (M and SD). Tests were two-tailed, and statistical significance was defined as *P* < .050.

**Figure 3 - supplement 2 (Figure 3 – supplement 2. jpg): Specificity antibody control shows detection of several members of the FH family.**

A specificity control of the polyclonal FH family antibody was conducted with human serum, FH-depleted (dpl) serum and purified FH, FHL-1, factor H-related protein 1 (FHR-1), factor H-related protein 2 (FHR-2), factor H-related protein 3 (FHR-3), factor H-related protein 4 (FHR-4), and factor H-related protein 5 (FHR-5). The antibody recognised FH at around 160 kDa, FHL-1 at 50 kDa, FHR-1 at 40 kDa, FHR-2 at 30 kDa, FHR-3 at 40 kDa and FHR-5 at 70 kDa.

**Figure 3 - supplement 3 (Figure 3 – supplement 3. jpg): Specificity antibody control shows exclusive detection of FH.**

A specificity control of the monoclonal FH antibody that specifically detects FH domain 16 was conducted with human serum, FH-depleted (dpl) serum and purified FH, FHL-1, FHR-1, FHR-2, FHR-3, FHR-4 and FHR-5. This antibody shows a band for FH at around 160 kDa and no bands for the remaining purified proteins.

**Figure 3 - supplement 4 (Figure 3 – supplement 4. jpg): Fluorescence intensity analysis of FH family, FH, and FI stainings show decreased intensities in mature cells.**

**(A)** Fluorescence intensity of F-actin is reduced in mature iPSC-RPE cells compared with immature cells, with no difference observed in ARPE-19 cells (*n* = 3). **(B)** Fluorescence intensity of FH is reduced in mature ARPE-19 cells compared with immature cells, with no difference observed in iPSC-RPE cells (*n* = 3). **(C)** Fluorescence intensity of FI is reduced in mature ARPE-19 cells compared with immature cells, with no difference observed in iPSC-RPE cells (*n* = 3). Non-parametric data (underlined) were analysed using the Wilcoxon test (Mdn and IQR); parametric data with independent-means *t*-test (M and SD). Tests were two-tailed, and statistical significance was defined as *P* < .050.

**Figure 4 - supplement 1 (Figure 4 – supplement 1. jpg): Specificity control of kit beads confirms detection of iC3b and C3c.**

Specificity control confirms C3 beads of the used kit detect C3, while C3b/iC3b beads primarily bind iC3b and C3c, with minor C3b detection (*n* = 3). For accuracy, C3b/iC3b beads were renamed iC3b beads.

**Figure 5 - supplement 1 (Figure 5 – supplement 1. jpg): No relevant C3 bands are visible in cell negative control of Western blot with cell media.**

Negative controls for C3 antibody Western blots were performed using cell media. Purified C3 and C3a served as positive controls, while DMEM-Gln (immature ARPE-19 medium), MEM-Nic (mature ARPE-19 medium), and iPSC-RPE medium were used as samples. **(A)** Detection with C3 antibody 1 showed signals for C3 α chain (∼115 kDa) and the C3 α’ chain (∼101 kDa) only in the C3 positive control. **(B)** Detection with C3 antibody 2 showed strong signals for the C3 α chain (∼115 kDa) and the β chain (∼75 kDa). A faint signal at ∼50 kDa was observed in DMEM-Gln, that is not relevant for Western blot analysis of RPE cell lysates. **(C)** Total protein staining of cell media for the blots shown in **(A)** and **(B)** confirmed protein loading.

**Figure 5 - supplement 2 (Figure 5 – supplement 2. jpg): Full membranes of Western blot analysis of purified proteins as positive controls alongside RPE cell lysate samples.**

Full blots including purified proteins as positive controls from the second replicate of the Western blots shown in **Figure 5B** and **E**. **(A)** Western blot stained with C3 antibody 1 (**Figure 5B**). Positive controls display clear bands for the C3 α chain in purified C3 and for C3d in purified C3d, with corresponding bands also visible in RPE cell lysates. **(B)** Western blot stained with C3 antibody 2 (**Figure 5E**). Positive controls exhibit clear bands for the C3 α chain in purified C3 and the β chain in purified C3, C3b, and iC3b, with corresponding bands also observed in RPE cell lysates.

**Figure 5 - supplement 3 (Figure 5 – supplement 3. jpg): Fluorescence intensity analysis of C3 stainings shows no differences.**

**(A)** Fluorescence intensity of C3 antibody 1 staining shows no significant difference between conditions (*n* = 3). **(B)** Fluorescence intensity of C3 antibody 2 staining shows no significant difference between conditions (*n* = 3). Non-parametric data (underlined) were analysed using the Wilcoxon test (Mdn and IQR); parametric data with independent-means *t*-test (M and SD). Tests were two-tailed, and statistical significance was defined as *P* < .050.

**Figure 5 - supplement 4 (Figure 5 – supplement 4. jpg): The full C3 β chain is only visible in mature iPSC-RPE cells.**

Immunoprecipitation analysis of immature (Im) and mature (Ma) ARPE-19 and iPSC-RPE cell lysates was conducted using a C3 β chain-specific antibody. **(A)** Lysates were immunoprecipitated with a C3 β chain-specific antibody bound to magnetic beads, followed by SDS-PAGE and Western blot. Detection was performed with **(B)** C3 antibody 2 and **(C)** the C3 β chain-specific antibody. A band corresponding to the C3 β chain at ∼75 kDa was observed exclusively in mature iPSC-RPE cells with both antibodies. **(D)** Total protein staining of precipitated RPE cell lysates, used to confirm protein loading for C3 β chain detection. M = marker.

**Figure 6 - supplement 1 (Figure 6 – supplement 1. jpg): Fluorescence intensity analysis of stainings of TGF-β1 signalling shows differences.**

**(A)** Fluorescence intensity of TGF-β1 is reduced in mature ARPE-19 cells compared with immature cells, but increased in mature iPSC-RPE cells (*n* = 3). **(B)** Fluorescence intensity of the TβR1 staining shows no significant difference between conditions (*n* = 3). Non-parametric data (underlined) were analysed using the Wilcoxon test (Mdn and IQR); parametric data with independent-means *t*-test (M and SD). Tests were two-tailed, and statistical significance was defined as *P* < .050.

**Figure 7 - supplement 1 (Figure 7 – supplement 1. jpg): Stress induction of ARPE-19 and iPSC-RPE cells alters TGF-β1 signalling, RPE marker transcript expression and complement regulation.**

**(A)** TGF-β1 treatment increases *TGFB1* transcript expression in ARPE-19 (*n* = 4). **(B)** Basal TGF-β1 secretion increases following treatment in both cell types (*n* = 4). **(C)** TGF-β1-treated ARPE-19 cells show diffuse TGF-β1 signals, while control cells are strong and more focussed. Control iPSC-RPE exhibit strong and punctate TGF-β1 signals that are absent in treated cells (**Figure 7 - video 1**). **(D)** Apical and basal FB secretion remains unchanged in TGF-β1-treated ARPE-19. Basal FB secretion is increased in TGF-β1-treated iPSC-RPE, while apical FB secretion shows no significant difference *(n* = 4). **(E)** TGF-β1 treatment reduces *CFH* transcript expression in ARPE-19 (*n* = 4). **(F)** Both apical and basal FH/FHL-1 secretion is reduced in TGF-β1-treated ARPE-19, while only apical FH/FHL-1 secretion decreases in TGF-β1-treated iPSC-RPE *(n* = 4). **(G)** *CFI* transcript expression decreases in TGF-β1-treated ARPE-19 (*n* = 4). **(H)** Both apical and basal FI secretion is reduced in TGF-β1-treated ARPE-19, while only apical FI secretion decreases in TGF-β1-treated iPSC-RPE *(n* = 4). Data represent pooled results from two independent treatment replicates. Non-parametric data (underlined) were analysed using the Wilcoxon test (Mdn and IQR); parametric data with independent-means *t*-test (M and SD). Tests were two-tailed, and statistical significance was defined as *P* < .050. Scale bar = 50 µm.

**Figure 7 - supplement 2 (Figure 7 – supplement 2. jpg): Fluorescence intensity analysis of TGF-β1 treatment stainings shows significant increase in treated iPSC-RPE cells for C3 antibody 1.**

**(A)** Fluorescence intensity of the TGF*-*β1 staining shows no significant difference between conditions (*n* = 3). **(B)** Fluorescence intensity of the F-actin staining shows no significant difference between conditions (*n* = 3). **(C)** Fluorescence intensity of the TJP1 staining shows no significant difference between conditions (*n* = 3). **(D)** Fluorescence intensity of the C3 antibody 1 intensity is increased in TGF*-*β1-treated iPSC-RPE cells compared with control cells, while no difference is observed in ARPE-19 cells (*n* = 3). Non-parametric data (underlined) were analysed using the Wilcoxon test (Mdn and IQR); parametric data with independent-means *t*-test (M and SD). Tests were two-tailed, and statistical significance was defined as *P* < .050.

## Appendix

**Appendix 1 (Appendix 1. jpg): No signals are visible in negative controls of secondary antibodies.**

**(A-D)** Negative controls without usage of primary antibodies in immature and mature ARPE-19 and iPSC-RPE cells show no immunofluorescent signals. Scale bar = 50 µm.

**Appendix 2 (Appendix 2.jpg): Transcriptomic analysis was conducted of ARPE-19 and RPESC-RPE cells using NCBI GEO tool GEO2R.**

(A) Analysis of the Samuel et al. dataset (GEO accession number GSE88848) shows elevated transcript levels of *CFH*, *CFI*, and *C3* in ARPE-19 cells with an RPE-typical phenotype following four months of cultivation (Long cultivation) compared to cells with shorter cultivation periods (Short cultivation) (Samuel et al., 2017). **(B)** Analysis of the Boles et al. dataset (GEO accession number GSE128143) reveals increased *CFH* and *CFI* transcript levels and decreased *C3* transcript levels in RPESC-RPE cells following combined TNF-α/TGF-β1 treatment (Combined treatment) compared to untreated controls (Control) (Boles et al., 2020).

**Appendix 3 (Appendix 3.xlsx): Detailed information about single nucleotide polymorphisms, self-designed primers, primary and secondary antibodies, purified proteins and sera used in this study.**

Overview of the analysed single nucleotide polymorphisms (rs1061170, rs2230199, rs10033900) in ARPE-19 and iPSC-RPE cells. Self-designed primer sequences are provided. Primary and secondary antibody sources are listed, including the anti-FH.16 antibody generously provided by Sanquin Research, Amsterdam, Netherlands. Sources of purified proteins and sera are indicated, including recombinant FHL-1 protein generously supplied by Mihály Józsi (Kopp et al., 2012).

## References

Ahlmann-Eltze, C., & Indrajeet, P. (2021). {ggsignif}: R Package for Displaying Significance Brackets for {’ggplot2’}. PsyArxiv. 10.31234/osf.io/7awm6

Al-Ani, A., Sunba, S., Hafeez, B., Toms, D., & Ungrin, M. (2020). In Vitro Maturation of Retinal Pigment Epithelium Is Essential for Maintaining High Expression of Key Functional Genes. International Journal of Molecular Sciences, 21(17), 6066. 10.3390/ijms21176066

Anderson, D. H., Mullins, R. F., Hageman, G. S., & Johnson, L. V. (2002). A role for local inflammation in the formation of drusen in the aging eye. American Journal of Ophthalmology, 134(3), 411–431. 10.1016/S0002-9394(02)01624-0

Anderson, D. H., Radeke, M. J., Gallo, N. B., Chapin, E. A., Johnson, P. T., Curletti, C. R., Hancox, L. S., Hu, J., Ebright, J. N., Malek, G., Hauser, M. A., Bowes Rickman, C., Bok, D., Hageman, G. S., & Johnson, L. V. (2010). The pivotal role of the complement system in aging and age-related macular degeneration: Hypothesis re-visited. Progress in Retinal and Eye Research, 29(2), 95–112. 10.1016/j.preteyeres.2009.11.003

Arbore, G., Kemper, C., & Kolev, M. (2017). Intracellular complement − the complosome − in immune cell regulation. Molecular Immunology, 89, 2–9. 10.1016/j.molimm.2017.05.012

Balaratnasingam, C., Messinger, J. D., Sloan, K. R., Yannuzzi, L. A., Freund, K. B., & Curcio, C. A. (2017). Histologic and Optical Coherence Tomographic Correlates in Drusenoid Pigment Epithelium Detachment in Age-Related Macular Degeneration. Ophthalmology, 124(5), 644–656. 10.1016/j.ophtha.2016.12.034

Barrett, T., Wilhite, S. E., Ledoux, P., Evangelista, C., Kim, I. F., Tomashevsky, M., Marshall, K. A., Phillippy, K. H., Sherman, P. M., Holko, M., Yefanov, A., Lee, H., Zhang, N., Robertson, C. L., Serova, N., Davis, S., & Soboleva, A. (2012). NCBI GEO: archive for functional genomics data sets—update. Nucleic Acids Research, 41(D1), D991–D995. 10.1093/nar/gks1193

Bharti, K., Miller, S. S., & Arnheiter, H. (2011). The new paradigm: retinal pigment epithelium cells generated from embryonic or induced pluripotent stem cells. Pigment Cell & Melanoma Research, 24(1), 21–34. 10.1111/j.1755-148X.2010.00772.x

Boles, N. C., Fernandes, M., Swigut, T., Srinivasan, R., Schiff, L., Rada-Iglesias, A., Wang, Q., Saini, J. S., Kiehl, T., Stern, J. H., Wysocka, J., Blenkinsop, T. A., & Temple, S. (2020). Epigenomic and Transcriptomic Changes During Human RPE EMT in a Stem Cell Model of Epiretinal Membrane Pathogenesis and Prevention by Nicotinamide. Stem Cell Reports, 14(4), 631–647. 10.1016/j.stemcr.2020.03.009

Bonilha, V. L., Bell, B. A., Hu, J., Milliner, C., Pauer, G. J., Hagstrom, S. A., Radu, R. A., & Hollyfield, J. G. (2020). Geographic Atrophy: Confocal Scanning Laser Ophthalmoscopy, Histology, and Inflammation in the Region of Expanding Lesions. Investigative Opthalmology & Visual Science, 61(8), 15. 10.1167/iovs.61.8.15

Cao, D., Leong, B., Messinger, J. D., Kar, D., Ach, T., Yannuzzi, L. A., Bailey Freund, K., & Curcio, C. A. (2021). Hyperreflective foci, optical coherence tomography progression indicators in age-related macular degeneration, include transdifferentiated retinal pigment epithelium. Investigative Ophthalmology and Visual Science, 62(10). 10.1167/IOVS.62.10.34

Chinchilla, B., & Fernandez-Godino, R. (2021). AMD-like substrate causes epithelial mesenchymal transition in iPSC-derived retinal pigment epithelial cells wild type but not C3-knockout. International Journal of Molecular Sciences, 22(15). 10.3390/ijms22158183

Chtcheglova, L. A., Ohlmann, A., Boytsov, D., Hinterdorfer, P., Priglinger, S. G., & Priglinger, C. S. (2020). Nanoscopic approach to study the early stages of epithelial to mesenchymal transition (EMT) of human retinal pigment epithelial (RPE) cells in vitro. Life, 10(8), 1–19. 10.3390/life10080128

Crabb, J. W., Miyagi, M., Gu, X., Shadrach, K., West, K. A., Sakaguchi, H., Kamei, M., Hasan, A., Yan, L., Rayborn, M. E., Salomon, R. G., & Hollyfield, J. G. (2002). Drusen proteome analysis: An approach to the etiology of age-related macular degeneration. Proceedings of the National Academy of Sciences, 99(23), 14682–14687. 10.1073/pnas.222551899

Dascalu, A. M., Grigorescu, C. C., Serban, D., Tudor, C., Alexandrescu, C., Stana, D., Jurja, S., Costea, A. C., Alius, C., Tribus, L. C., Dumitrescu, D., Bratu, D., & Cristea, B. M. (2024). Complement Inhibitors for Geographic Atrophy in Age-Related Macular Degeneration—A Systematic Review. Journal of Personalized Medicine, 14(9), 990. 10.3390/jpm14090990

Despriet, D. D. G., van Duijn, C. M., Oostra, B. A., Uitterlinden, A. G., Hofman, A., Wright, A. F., ten Brink, J. B., Bakker, A., de Jong, P. T. V. M., Vingerling, J. R., Bergen, A. A. B., & Klaver, C. C. W. (2009). Complement Component C3 and Risk of Age-Related Macular Degeneration. Ophthalmology, 116(3), 474–480.e2. 10.1016/j.ophtha.2008.09.055

Ebrahimi, K. B., Fijalkowski, N., Cano, M., & Handa, J. T. (2013). Decreased membrane complement regulators in the retinal pigmented epithelium contributes to age-related macular degeneration. The Journal of Pathology, 229(5), 729–742. 10.1002/path.4128

Edwards, A. O., Ritter, R., Abel, K. J., Manning, A., Panhuysen, C., & Farrer, L. A. (2005). Complement factor H polymorphism and age-related macular degeneration. Science, 308(5720), 421–424. 10.1126/science.1110189

Elvington, M., Liszewski, M. K., Liszewski, A. R., Kulkarni, H. S., Hachem, R. R., Mohanakumar, T., Kim, A. H. J., & Atkinson, J. P. (2019). Development and optimization of an ELISA to quantitate C3(H2O) as a marker of human disease. Frontiers in Immunology, 10(APR), 1–12. 10.3389/fimmu.2019.00703

Enzbrenner, A., Zulliger, R., Biber, J., Pousa, A. M. Q., Schäfer, N., Stucki, C., Giroud, N., Berrera, M., Kortvely, E., Schmucki, R., Badi, L., Grosche, A., Pauly, D., & Enzmann, V. (2021). Sodium iodate-induced degeneration results in local complement changes and inflammatory processes in murine retina. International Journal of Molecular Sciences, 22(17), 1–16. 10.3390/ijms22179218

Fernandez-Godino, R. (2018). Alterations in extracellular matrix/bruch’s membrane can cause the activation of the alternative complement pathway via tick-over. Advances in Experimental Medicine and Biology, 1074, 29–35. 10.1007/978-3-319-75402-4_4

Ferris, F. L., Wilkinson, C. P., Bird, A., Chakravarthy, U., Chew, E., Csaky, K., & Sadda, S. R. (2013). Clinical classification of age-related macular degeneration. Ophthalmology, 120(4), 844–851. 10.1016/j.ophtha.2012.10.036

Fields, M. A., Bowrey, H. E., Gong, J., Moreira, E. F., Cai, H., & Del Priore, L. V. (2017). Extracellular matrix nitration alters growth factor release and activates bioactive complement in human retinal pigment epithelial cells. PLOS ONE, 12(5), e0177763. 10.1371/journal.pone.0177763

Flaxman, S. R., Bourne, R. R. A., Resnikoff, S., Ackland, P., Braithwaite, T., Cicinelli, M. V, Das, A., Jonas, J. B., Keeffe, J., Kempen, J. H., Leasher, J., Limburg, H., Naidoo, K., Pesudovs, K., Silvester, A., Stevens, G. A., Tahhan, N., Wong, T. Y., Taylor, H. R., … Zheng, Y. (2017). Global causes of blindness and distance vision impairment 1990–2020: a systematic review and meta-analysis. The Lancet Global Health, 5(12), e1221–e1234. 10.1016/S2214-109X(17)30393-5

Fox, J. (2022). polycor: Polychoric and Polyserial Correlations. https://cran.r-project.org/package=polycor

Fox, J., & Weisberg, S. (2019). An {R} Companion to Applied Regression. https://socialsciences.mcmaster.ca/jfox/Books/Companion/

Fritsche, L. G., Chen, W., Schu, M., Yaspan, B. L., Yu, Y., Thorleifsson, G., Zack, D. J., Arakawa, S., Cipriani, V., Ripke, S., Igo, R. P., Buitendijk, G. H. S., Sim, X., Weeks, D. E., Guymer, R. H., Merriam, J. E., Francis, P. J., Hannum, G., Agarwal, A., … Abecasis, G. R. (2013). Seven new loci associated with age-related macular degeneration. Nature Genetics, 45(4), 433–439. 10.1038/ng.2578.Seven

Fritsche, L. G., Igl, W., Bailey, J. N. C., Grassmann, F., Sengupta, S., Bragg-Gresham, J. L., Burdon, K. P., Hebbring, S. J., Wen, C., Gorski, M., Kim, I. K., Cho, D., Zack, D., Souied, E., Scholl, H. P. N., Bala, E., ELee, K., Hunter, D. J., Sardell, R. J., … Heid, I. M. (2016). A large genome-wide association study of age-related macular degeneration highlights contributions of rare and common variants. Nature Genetics, 48(2), 134–143. 10.1038/ng.3448

Gambril, J. A., Sloan, K. R., Swain, T. A., Huisingh, C., Zarubina, A. V., Messinger, J. D., Ach, T., & Curcio, C. A. (2019). Quantifying retinal pigment epithelium dysmorphia and loss of histologic auto fluorescence in age-related macular degeneration. Investigative Ophthalmology and Visual Science, 60(7), 2481–2493. 10.1167/iovs.19-26949

Ghosh, S., Shang, P., Terasaki, H., Stepicheva, N., Hose, S., Yazdankhah, M., Weiss, J., Sakamoto, T., Bhutto, I. A., Xia, S., Zigler, J. S., Kannan, R., Qian, J., Handa, J. T., & Sinha, D. (2018). A Role for βA3/A1-Crystallin in Type 2 EMT of RPE Cells Occurring in Dry Age-Related Macular Degeneration. Investigative Opthalmology & Visual Science, 59(4), AMD104. 10.1167/iovs.18-24132

Grosjean, P., & Ibanez, F. (2018). pastecs: Package for Analysis of Space-Time Ecological Series. https://cran.r-project.org/package=pastecs

Hageman, G. S., Anderson, D. H., Johnson, L. V., Hancox, L. S., Taiber, A. J., Hardisty, L. I., Hageman, J. L., Stockman, H. A., Borchardt, J. D., Gehrs, K. M., Smith, R. J. H., Silvestri, G., Russell, S. R., Klaver, C. C. W., Barbazetto, I., Chang, S., Yannuzzi, L. A., Barile, G. R., Merriam, J. C., … Allikmets, R. (2005). A common haplotype in the complement regulatory gene factor H (HF1/CFH) predisposes individuals to age-related macular degeneration. Proceedings of the National Academy of Sciences, 102(20), 7227–7232. 10.1073/pnas.0501536102

Hansen, J. N., Rassmann, S., Stüven, B., Jurisch-Yaksi, N., & Wachten, D. (2021). CiliaQ: a simple, open-source software for automated quantification of ciliary morphology and fluorescence in 2D, 3D, and 4D images. The European Physical Journal E, 44(2), 18. 10.1140/epje/s10189-021-00031-y

Hazim, R. A., Volland, S., Yen, A., Burgess, B. L., & Williams, D. S. (2019). Rapid differentiation of the human RPE cell line, ARPE-19, induced by nicotinamide. Experimental Eye Research, 179, 18–24. 10.1016/j.exer.2018.10.009

Heesterbeek, T. J., Lorés-Motta, L., Hoyng, C. B., Lechanteur, Y. T. E., & den Hollander, A. I. (2020). Risk factors for progression of age-related macular degeneration. Ophthalmic and Physiological Optics, 40(2), 140–170. 10.1111/opo.12675

Heier, J. S., Lad, E. M., Holz, F. G., Rosenfeld, P. J., Guymer, R. H., Boyer, D., Grossi, F., Baumal, C. R., Korobelnik, J.-F., Slakter, J. S., Waheed, N. K., Metlapally, R., Pearce, I., Steinle, N., Francone, A. A., Hu, A., Lally, D. R., Deschatelets, P., Francois, C., … OAKS and DERBY study investigators. (2023). Pegcetacoplan for the treatment of geographic atrophy secondary to age-related macular degeneration (OAKS and DERBY): two multicentre, randomised, double-masked, sham-controlled, phase 3 trials. Lancet (London, England), 402(10411), 1434–1448. 10.1016/S0140-6736(23)01520-9

Isaac, L., & Isenman, D. E. (1992). Structural requirements for thioester bond formation in human complement component C3. Reassessment of the role of thioester bond integrity on the conformation of C3. The Journal of Biological Chemistry, 267(14), 10062–10069.

Johnson, P. T., Betts, K. E., Radeke, M. J., Hageman, G. S., Anderson, D. H., & Johnson, L. V. (2006). Individuals homozygous for the age-related macular degeneration risk-conferring variant of complement factor H have elevated levels of CRP in the choroid. Proceedings of the National Academy of Sciences, 103(46), 17456–17461. 10.1073/pnas.0606234103

Johnson, L. V., Leitner, W. P., Staples, M. K., & Anderson, D. H. (2001). Complement activation and inflammatory processes in drusen formation and age related macular degeneration. Experimental Eye Research, 73(6), 887–896. 10.1006/exer.2001.1094

Kim, S.-J., Kim, Y.-S., Kim, J. H., Jang, H. Y., Ly, D. Da, Das, R., & Park, K.-S. (2020). Activation of ERK1/2-mTORC1-NOX4 mediates TGF-β1-induced epithelial-mesenchymal transition and fibrosis in retinal pigment epithelial cells. Biochemical and Biophysical Research Communications, 529(3), 747–752. 10.1016/j.bbrc.2020.06.034

Kim, S., & Dynlacht, B. D. (2013). Assembling a primary cilium. Current Opinion in Cell Biology, 25(4), 506–511. 10.1016/j.ceb.2013.04.011

Kim, Y. H., He, S., Kase, S., Kitamura, M., Ryan, S. J., & Hinton, D. R. (2009). Regulated secretion of complement factor H by RPE and its role in RPE migration. Graefe’s Archive for Clinical and Experimental Ophthalmology, 247(5), 651–659. 10.1007/s00417-009-1049-y

Klein, R. J., Zeiss, C., Chew, E. Y., Tsai, J., Sackler, R. S., Haynes, C., Henning, A. K., Sangiovanni, J. P., Mane, S. M., Susan, T., Bracken, M. B., Ferris, F. L., Ott, J., Barnstable, C., & Hoh, J. (2005). Complement Factor H Polymorphism in Age-Related Macular Degeneration. Science, 308(5720), 385–389. 10.1126/science.1109557.Complement

Kopp, A., Strobel, S., Tortajada, A., Rodríguez de Córdoba, S., Sánchez-Corral, P., Prohászka, Z., López-Trascasa, M., & Józsi, M. (2012). Atypical Hemolytic Uremic Syndrome-Associated Variants and Autoantibodies Impair Binding of Factor H and Factor H-Related Protein 1 to Pentraxin 3. The Journal of Immunology, 189(4), 1858–1867. 10.4049/jimmunol.1200357

Kretschmer, V., Schneider, S., Matthiessen, P. A., Reichert, D., Hotaling, N., Glasßer, G., Lieberwirth, I., Bharti, K., De Cegli, R., Conte, I., Nandrot, E. F., & May-Simera, H. L. (2023). Deletion of IFT20 exclusively in the RPE ablates primary cilia and leads to retinal degeneration. PLOS Biology, 21(12), e3002402. 10.1371/journal.pbio.3002402

Kunchithapautham, K., Atkinson, C., & Rohrer, B. (2014). Smoke Exposure Causes Endoplasmic Reticulum Stress and Lipid Accumulation in Retinal Pigment Epithelium through Oxidative Stress and Complement Activation. Journal of Biological Chemistry, 289(21), 14534–14546. 10.1074/jbc.M114.564674

Laine, M., Jarva, H., Seitsonen, S., Haapasalo, K., Lehtinen, M. J., Lindeman, N., Anderson, D. H., Johnson, P. T., Järvelä, I., Jokiranta, T. S., Hageman, G. S., Immonen, I., & Meri, S. (2007). Y402H Polymorphism of Complement Factor H Affects Binding Affinity to C-Reactive Protein. The Journal of Immunology, 178(6), 3831–3836. 10.4049/jimmunol.178.6.3831

Lakkaraju, A., Umapathy, A., Tan, L. X., Daniele, L., Philp, N. J., Boesze-Battaglia, K., & Williams, D. S. (2020). The cell biology of the retinal pigment epithelium. Progress in Retinal and Eye Research, 78(February), 100846. 10.1016/j.preteyeres.2020.100846

Lee, J., Choi, J.-H., & Joo, C.-K. (2013). TGF-β1 regulates cell fate during epithelial–mesenchymal transition by upregulating survivin. Cell Death & Disease, 4(7), e714–e714. 10.1038/cddis.2013.244

Li, M., Li, H., Yang, S., Liao, X., Zhao, C., & Wang, F. (2021). L-carnitine attenuates TGF-β1-induced EMT in retinal pigment epithelial cells via a PPARγ-dependent mechanism. International Journal of Molecular Medicine, 47(6), 110. 10.3892/ijmm.2021.4943

Li, Y., Song, D., Song, Y., Zhao, L., Wolkow, N., Tobias, J. W., Song, W., & Dunaief, J. L. (2015). Iron-induced Local Complement Component 3 (C3) Up-regulation via Non-canonical Transforming Growth Factor (TGF)-β Signaling in the Retinal Pigment Epithelium. Journal of Biological Chemistry, 290(19), 11918–11934. 10.1074/jbc.M115.645903

Liao, D. S., Grossi, F. V., El Mehdi, D., Gerber, M. R., Brown, D. M., Heier, J. S., Wykoff, C. C., Singerman, L. J., Abraham, P., Grassmann, F., Nuernberg, P., Weber, B. H. F., Deschatelets, P., Kim, R. Y., Chung, C. Y., Ribeiro, R. M., Hamdani, M., Rosenfeld, P. J., Boyer, D. S., … Francois, C. G. (2020). Complement C3 Inhibitor Pegcetacoplan for Geographic Atrophy Secondary to Age-Related Macular Degeneration. Ophthalmology, 127(2), 186–195. 10.1016/j.ophtha.2019.07.011

Lindell, M., Kar, D., Sedova, A., Kim, Y. J., Packer, O. S., Schmidt-Erfurth, U., Sloan, K. R., Marsh, M., Dacey, D. M., Curcio, C. A., & Pollreisz, A. (2023). Volumetric Reconstruction of a Human Retinal Pigment Epithelial Cell Reveals Specialized Membranes and Polarized Distribution of Organelles. Investigative Opthalmology & Visual Science, 64(15), 35. 10.1167/iovs.64.15.35

Little, K., Llorián-Salvador, M., Tang, M., Du, X., Marry, S., Chen, M., & Xu, H. (2020). Macrophage to myofibroblast transition contributes to subretinal fibrosis secondary to neovascular age-related macular degeneration. Journal of Neuroinflammation, 17(1), 355. 10.1186/s12974-020-02033-7

Little, K., Llorián-Salvador, M., Tang, M., Du, X., O’Shaughnessy, Ó., McIlwaine, G., Chen, M., & Xu, H. (2020). A Two-Stage Laser-Induced Mouse Model of Subretinal Fibrosis Secondary to Choroidal Neovascularization. Translational Vision Science & Technology, 9(4), 3. 10.1167/tvst.9.4.3

Luo, C., Chen, M., & Xu, H. (2011). Complement gene expression and regulation in mouse retina and retinal pigment epithelium/choroid. Molecular Vision, 17(June), 1588–1597.

Ma, X., Xie, Y., Gong, Y., Hu, C., Qiu, K., Yang, Y., Shen, H., Zhou, X., Long, C., & Lin, X. (2023). Silibinin Prevents TGFβ-Induced EMT of RPE in Proliferative Vitreoretinopathy by Inhibiting Stat3 and Smad3 Phosphorylation. Investigative Opthalmology & Visual Science, 64(13), 47. 10.1167/iovs.64.13.47

Maller, J. B., Fagerness, J. A., Reynolds, R. C., Neale, B. M., Daly, M. J., & Seddon, J. M. (2007). Variation in complement factor 3 is associated with risk of age-related macular degeneration. Nature Genetics, 39(10), 1200–1201. 10.1038/ng2131

Marmorstein, A. D., Marmorstein, L. Y., Rayborn, M., Wang, X., Hollyfield, J. G., & Petrukhin, K. (2000). Bestrophin, the product of the Best vitelliform macular dystrophy gene (VMD2), localizes to the basolateral plasma membrane of the retinal pigment epithelium. Proceedings of the National Academy of Sciences, 97(23), 12758–12763. 10.1073/pnas.220402097

Martin, M., Leffler, J., Smolag, K. I., Mytych, J., Björk, A., Chaves, L. D., Alexander, J. J., Quigg, R. J., & Blom, A. M. (2016). Factor H uptake regulates intracellular C3 activation during apoptosis and decreases the inflammatory potential of nucleosomes. Cell Death and Differentiation, 23(5), 903–911. 10.1038/cdd.2015.164

May-Simera, H. L., Wan, Q., Jha, B. S., Hartford, J., Khristov, V., Dejene, R., Chang, J., Patnaik, S., Lu, Q., Banerjee, P., Silver, J., Insinna-Kettenhofen, C., Patel, D., Lotfi, M., Malicdan, M., Hotaling, N., Maminishkis, A., Sridharan, R., Brooks, B., … Bharti, K. (2018). Primary Cilium-Mediated Retinal Pigment Epithelium Maturation Is Disrupted in Ciliopathy Patient Cells. Cell Reports, 22(1), 189–205. 10.1016/j.celrep.2017.12.038

Merle, N. S., Church, S. E., Fremeaux-Bacchi, V., & Roumenina, L. T. (2015). Complement system part I - molecular mechanisms of activation and regulation. Frontiers in Immunology, 6(JUN), 1–30. 10.3389/fimmu.2015.00262

Merle, N. S., Noe, R., Halbwachs-Mecarelli, L., Fremeaux-Bacchi, V., & Roumenina, L. T. (2015). Complement system part II: Role in immunity. Frontiers in Immunology, 6(MAY), 1–26. 10.3389/fimmu.2015.00257

Miyatani, T., Tanaka, H., Numa, K., Uehara, A., Otsuki, Y., Hamuro, J., Kinoshita, S., & Sotozono, C. (2024). Clustered ARPE-19 cells distinct in mitochondrial membrane potential may play a pivotal role in cell differentiation. Scientific Reports, 14(1), 22391. 10.1038/s41598-024-73145-w

Neuwirth, E. (2022). RColorBrewer: ColorBrewer Palettes. https://cran.r-project.org/package=RColorBrewer

Newman, A. M., Gallo, N. B., Hancox, L. S., Miller, N. J., Radeke, C. M., Maloney, M. A., Cooper, J. B., Hageman, G. S., Anderson, D. H., Johnson, L. V, & Radeke, M. J. (2012). Systems-level analysis of age-related macular degeneration reveals global biomarkers and phenotype-specific functional networks. Genome Medicine, 4(2), 16. 10.1186/gm315

Ning, K., Bhuckory, M. B., Lo, C.-H., Sendayen, B. E., Kowal, T. J., Chen, M., Bansal, R., Chang, K.-C., Vollrath, D., Berbari, N. F., Mahajan, V. B., Hu, Y., & Sun, Y. (2023). Cilia-associated wound repair mediated by IFT88 in retinal pigment epithelium. Scientific Reports, 13(1), 8205. 10.1038/s41598-023-35099-3

Park, K. H., Fridley, B. L., Ryu, E., Tosakulwong, N., & Edwards, A. O. (2009). Complement Component 3 (C3) Haplotypes and Risk of Advanced Age-Related Macular Degeneration. Investigative Opthalmology & Visual Science, 50(7), 3386. 10.1167/iovs.08-3231

Patnaik, S. R., Kretschmer, V., Brücker, L., Schneider, S., Volz, A.-K., Oancea-Castillo, L. del R., & May-Simera, H. L. (2019). Bardet–Biedl Syndrome proteins regulate cilia disassembly during tissue maturation. Cellular and Molecular Life Sciences, 76(4), 757–775. 10.1007/s00018-018-2966-x

Pauly, D., Agarwal, D., Dana, N., Schäfer, N., Biber, J., Wunderlich, K. A., Jabri, Y., Straub, T., Zhang, N. R., Gautam, A. K., Weber, B. H. F., Hauck, S. M., Kim, M., Curcio, C. A., Stambolian, D., Li, M., & Grosche, A. (2019). Cell-Type-Specific Complement Expression in the Healthy and Diseased Retina. Cell Reports, 29(9), 2835–2848.e4. 10.1016/j.celrep.2019.10.084

Pfau, M., Schmitz-Valckenberg, S., Ribeiro, R., Safaei, R., McKeown, A., Fleckenstein, M., & Holz, F. G. (2022). Association of complement C3 inhibitor pegcetacoplan with reduced photoreceptor degeneration beyond areas of geographic atrophy. Scientific Reports, 12(1), 17870. 10.1038/s41598-022-22404-9

R Core Team. (2021). R: A Language and Environment for Statistical Computing. https://www.r-project.org/

Radeke, M. J., Radeke, C. M., Shih, Y.-H., Hu, J., Bok, D., Johnson, L. V., & Coffey, P. J. (2015). Restoration of mesenchymal retinal pigmented epithelial cells by TGFβ pathway inhibitors: implications for age-related macular degeneration. Genome Medicine, 7(1), 58. 10.1186/s13073-015-0183-x

Ricklin, D., Reis, E. S., Mastellos, D. C., Gros, P., & Lambris, J. D. (2016). Complement component C3 – The “Swiss Army Knife” of innate immunity and host defense. Immunological Reviews, 274(1), 33–58. 10.1111/imr.12500

Rutar, M., Natoli, R., Albarracin, R., Valter, K., & Provis, J. (2012). 670-nm light treatment reduces complement propagation following retinal degeneration. Journal of Neuroinflammation, 9(1), 724. 10.1186/1742-2094-9-257

Samuel, W., Jaworski, C., Postnikova, O. A., Kutty, R. K., Duncan, T., Tan, L. X., Poliakov, E., Lakkaraju, A., & Redmond, T. M. (2017). Appropriately differentiated ARPE-19 cells regain phenotype and gene expression profiles similar to those of native RPE cells. Molecular Vision, 23(June 2016), 60–89.

Schäfer, N., Grosche, A., Schmitt, S. I., Braunger, B. M., & Pauly, D. (2017). Complement components showed a time-dependent local expression pattern in constant and acute white light-induced photoreceptor damage. Frontiers in Molecular Neuroscience, 10(June), 1–17. 10.3389/fnmol.2017.00197

Schäfer, N., Rasras, A., Ormenisan, D. M., Amslinger, S., Enzmann, V., Jägle, H., & Pauly, D. (2021). Complement Factor H-Related 3 Enhanced Inflammation and Complement Activation in Human RPE Cells. Frontiers in Immunology, 12(November), 1–16. 10.3389/fimmu.2021.769242

Schäfer, N., Wolf, H. N., Enzbrenner, A., Schikora, J., Reichenthaler, M., Enzmann, V., & Pauly, D. (2020). Properdin modulates complement component production in stressed human primary retinal pigment epithelium cells. Antioxidants, 9(9), 1–19. 10.3390/antiox9090793

Schindelin, J., Arganda-Carreras, I., Frise, E., Kaynig, V., Longair, M., Pietzsch, T., Preibisch, S., Rueden, C., Saalfeld, S., Schmid, B., Tinevez, J.-Y., White, D. J., Hartenstein, V., Eliceiri, K., Tomancak, P., & Cardona, A. (2012). Fiji: an open-source platform for biological-image analysis. Nature Methods, 9(7), 676–682. 10.1038/nmeth.2019

Schneider, S., De Cegli, R., Nagarajan, J., Kretschmer, V., Matthiessen, P. A., Intartaglia, D., Hotaling, N., Ueffing, M., Boldt, K., Conte, I., & May-Simera, H. L. (2021). Loss of Ciliary Gene Bbs8 Results in Physiological Defects in the Retinal Pigment Epithelium. Frontiers in Cell and Developmental Biology, 9. 10.3389/fcell.2021.607121

Seddon, J. M. (2005). The US Twin Study of Age-Related Macular Degeneration. Archives of Ophthalmology, 123(3), 321. 10.1001/archopht.123.3.321

Sharma, R., George, A., Nimmagadda, M., Ortolan, D., Karla, B., Qureshy, Z., Bose, D., Dejene, R., Liang, G., Wan, Q., Chang, J., Jha, B. S., Memon, O., Miyagishima, K. J., Rising, A., Lal, M., Hanson, E., King, R., Campos, M. M., … Bharti, K. (2021). Epithelial phenotype restoring drugs suppress macular degeneration phenotypes in an iPSC model. Nature Communications, 12(1), 7293. 10.1038/s41467-021-27488-x

Sim, R. B., & Sim, E. (1983). Autolytic fragmentation of complement components C3 and C4 and its relationship to covalent binding activity. Annals of the New York Academy of Sciences, 421(1), 259–276. 10.1111/j.1749-6632.1983.tb18114.x

Soltis, R. D., Hasz, D., Morris, M. J., & Wilson, I. D. (1979). The effect of heat inactivation of serum on aggregation of immunoglobulins. Immunology, 36(1), 37–45.

Sorokin, S. (1962). Centrioles and the formation of rudimentary cilia by fibroblasts and smooth muscle cells. The Journal of Cell Biology, 15(2), 363–377. 10.1083/jcb.15.2.363

Sorokin, S. P. (1968). Reconstructions of centriole formation and ciliogenesis in mammalian lungs. Journal of Cell Science, 3(2), 207–230. 10.1242/jcs.3.2.207

Sripathi, S. R., Hu, M. W., Turaga, R. C., Mertz, J., Liu, M. M., Wan, J., Maruotti, J., Wahlin, K. J., Berlinicke, C. A., Qian, J., & Zack, D. J. (2021). Proteome Landscape of Epithelial-to-Mesenchymal Transition (EMT) of Retinal Pigment Epithelium Shares Commonalities with Malignancy-Associated EMT. Molecular and Cellular Proteomics, 20, 100131. 10.1016/J.MCPRO.2021.100131

Steinle, N. C., Pearce, I., Monés, J., Metlapally, R., Saroj, N., Hamdani, M., Ribeiro, R., Rosenfeld, P. J., & Lad, E. M. (2021). Impact of Baseline Characteristics on Geographic Atrophy Progression in the FILLY Trial Evaluating the Complement C3 Inhibitor Pegcetacoplan. American Journal of Ophthalmology, 227, 116–124. 10.1016/j.ajo.2021.02.031

Su, Y., Tang, Z., & Wang, F. (2021). Role of LINC01592 in TGF-β1-induced epithelial-mesenchymal transition of retinal pigment epithelial cells. Aging, 13(10), 14053–14064. 10.18632/aging.203023

Sugita, S., Makabe, K., Fujii, S., & Takahashi, M. (2018). Detection of complement activators in immune attack eyes after iPS-derived retinal pigment epithelial cell transplantation. Investigative Ophthalmology and Visual Science, 59(10), 4198–4209. 10.1167/iovs.18-24769

Tamiya, S., Liu, L., & Kaplan, H. J. (2010). Epithelial-Mesenchymal Transition and Proliferation of Retinal Pigment Epithelial Cells Initiated upon Loss of Cell-Cell Contact. Investigative Opthalmology & Visual Science, 51(5), 2755. 10.1167/iovs.09-4725

Tian, H., Xu, J., Tian, Y., Cao, Y., Lian, C., Ou, Q., & Wu, B. (2018). A cell culture condition that induces the mesenchymal-epithelial transition of dedifferentiated porcine retinal pigment epithelial cells. Experimental Eye Research. 10.1016/j.exer.2018.08.005

Trakkides, T. O., Schäfer, N., Reichenthaler, M., Kühn, K., Brandwijk, R. J. M. G. E., Toonen, E. J. M., Urban, F., Wegener, J., Enzmann, V., & Pauly, D. (2019). Oxidative stress increases endogenous complement-dependent inflammatory and angiogenic responses in retinal pigment epithelial cells independently of exogenous complement sources. Antioxidants, 8(11), 1–16. 10.3390/antiox8110548

Triglia, R. P., & Linscott, W. D. (1980). Titers of nine complement components, conglutinin and C3b-inactivator in adult and fetal bovine sera. Molecular Immunology, 17(6), 741–748. 10.1016/0161-5890(80)90144-3

Wang, L., Kondo, N., Cano, M., Ebrahimi, K., Yoshida, T., Barnett, B. P., Biswal, S., & Handa, J. T. (2014). Nrf2 Signaling Modulates Cigarette Smoke Induced Complement Activation in Retinal Pigmented Epithelial Cells. Free Radical Biology & Medicine, 70, 155–166. 10.1038/jid.2014.371

Wang, M., Wei, J., Li, H., & Wang, F. (2022). Changes in transepithelial electrical resistance and intracellular ion concentration in TGF-β-induced epithelial-mesenchymal transition of retinal pigment epithelial cells. American Journal of Translational Research, 14(4), 2728–2738.

Wang, Y., Shen, H., Pang, L., Qiu, B., Yuan, Y., Guan, X., & Xiang, X. (2023). Qihuang Granule protects the retinal pigment epithelium from oxidative stress via regulation of the alternative complement pathway. BMC Complementary Medicine and Therapies, 23(1), 55. 10.1186/s12906-023-03884-2

Warner, G. R., Bolker, B., Lumley, T., & Johnson, R. C. (2022). gmodels: Various R Programming Tools for Model Fitting. https://cran.r-project.org/package=gmodels

Wei, Q., Liu, Q., Ren, C., Liu, J., Cai, W., Zhu, M., Jin, H., He, M., & Yu, J. (2018). Effects of bradykinin on TGF-β1-induced epithelial-mesenchymal transition in ARPE-19 cells. Molecular Medicine Reports. 10.3892/mmr.2018.8556

Wickham, H., Averick, M., Bryan, J., Chang, W., D’Agostino McGowan, L., François, R., Grolemund, G., Hayes, A., Henry, L., Hester, J., Kuhn, M., Pedersen, T. L., Miller, E., Bache, S. M., Müller, K., Ooms, J., Robinson, D., Seidel, D. P., Spinu, V., … Yutani, H. (2019). Welcome to the {tidyverse}. Journal of Open Source Software, 4(43), 1686. 10.21105/joss.01686

Wong, W. L., Su, X., Li, X., Cheung, C. M. G., Klein, R., Cheng, C.-Y., & Wong, T. Y. (2014). Global prevalence of age-related macular degeneration and disease burden projection for 2020 and 2040: a systematic review and meta-analysis. The Lancet Global Health, 2(2), e106–e116. 10.1016/S2214-109X(13)70145-1

Yates, J. R. W., Sepp, T., Matharu, B. K., Khan, J. C., Thurlby, D. A., Shahid, H., Clayton, D. G., Hayward, C., Morgan, J., Wright, A. F., Armbrecht, A. M., Dhillon, B., Deary, I. J., Redmond, E., Bird, A. C., & Moore, A. T. (2007). Complement C3 variant and the risk of age-related macular degeneration. New England Journal of Medicine, 357(6), 553–561. 10.1056/NEJMoa072618

Zanzottera, E. C., Ach, T., Huisingh, C., Messinger, J. D., Spaide, R. F., & Curcio, C. A. (2016). Visualizing retinal pigment epithelium phenotypes in the transition to geographic atrophy in age-related macular degeneration. Retina, 36(Supplement 1), S12–S25. 10.1097/IAE.0000000000001276

Zanzottera, E. C., Messinger, J. D., Ach, T., Smith, R. T., & Curcio, C. A. (2015). Subducted and Melanotic Cells in Advanced Age-Related Macular Degeneration Are Derived From Retinal Pigment Epithelium. Investigative Opthalmology & Visual Science, 56(5), 3269. 10.1167/iovs.15-16432

Zanzottera, E. C., Messinger, J. D., Ach, T., Smith, R. T., Freund, K. B., & Curcio, C. A. (2015). The Project MACULA Retinal Pigment Epithelium Grading System for Histology and Optical Coherence Tomography in Age-Related Macular Degeneration. Investigative Opthalmology & Visual Science, 56(5), 3253. 10.1167/iovs.15-16431

Zauhar, R., Biber, J., Jabri, Y., Kim, M., Hu, J., Kaplan, L., Pfaller, A. M., Schäfer, N., Enzmann, V., Schlötzer-Schrehardt, U., Straub, T., Hauck, S. M., Gamlin, P. D., McFerrin, M. B., Messinger, J., Strang, C. E., Curcio, C. A., Dana, N., Pauly, D., … Stambolian, D. (2022). As in Real Estate, Location Matters: Cellular Expression of Complement Varies Between Macular and Peripheral Regions of the Retina and Supporting Tissues. Frontiers in Immunology, 13(June), 1–19. 10.3389/fimmu.2022.895519

