## Supplementary figures and images for "Maturation-dependent complement production and C3 processing in human retinal pigment epithelium cells"

### Appendix 1

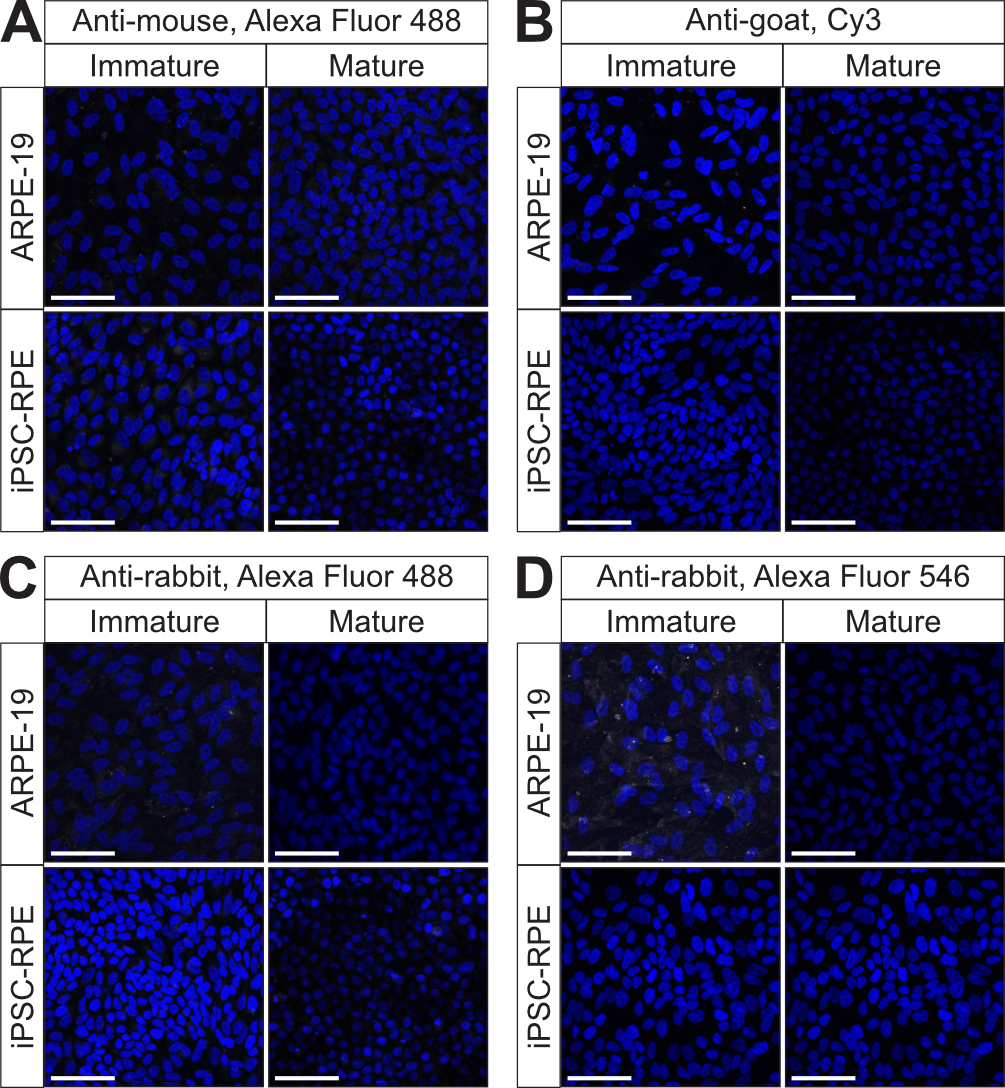

### Appendix 2

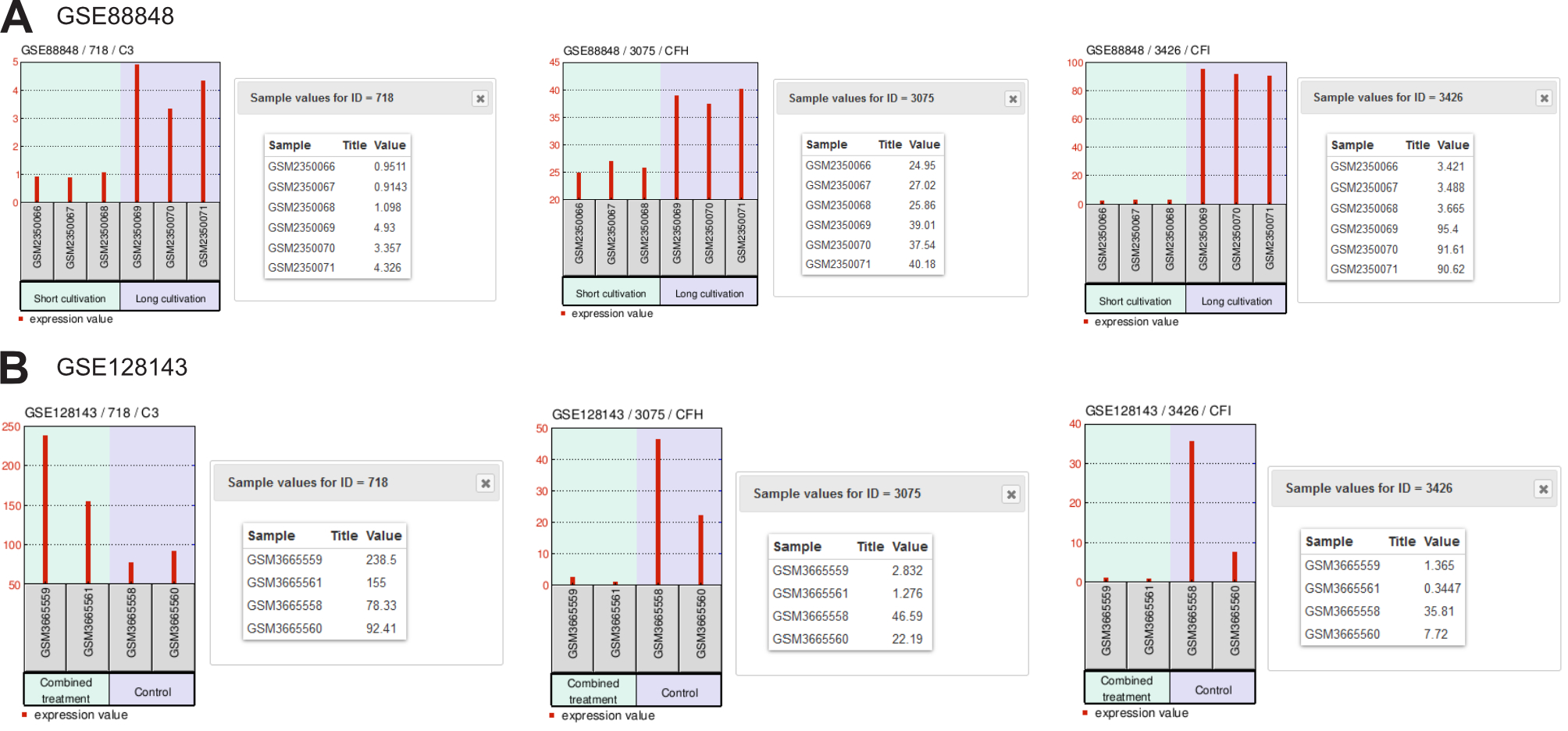

### Figure 1 - figure supplement 1

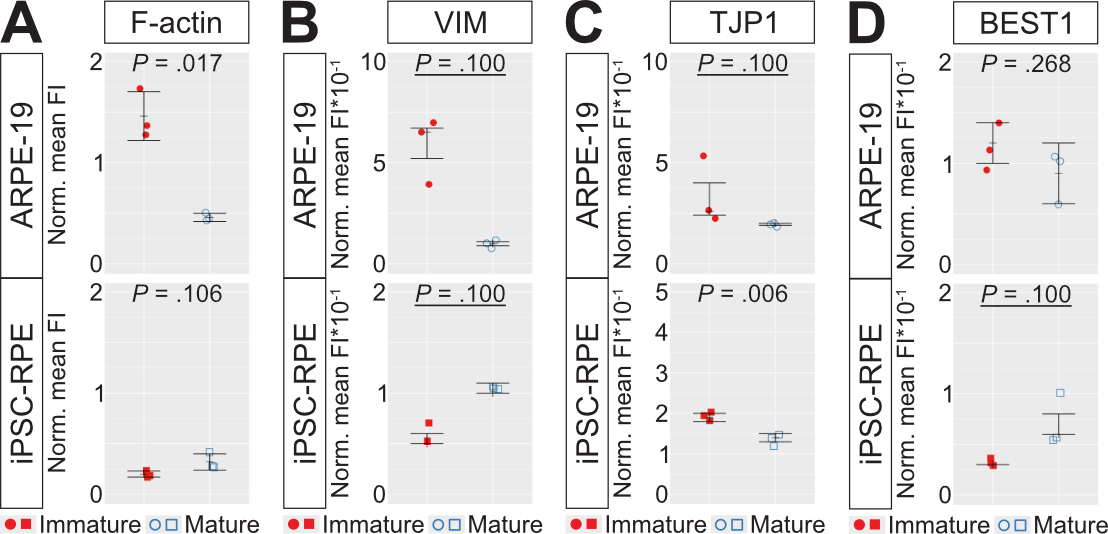

### Figure 1 - figure supplement 2

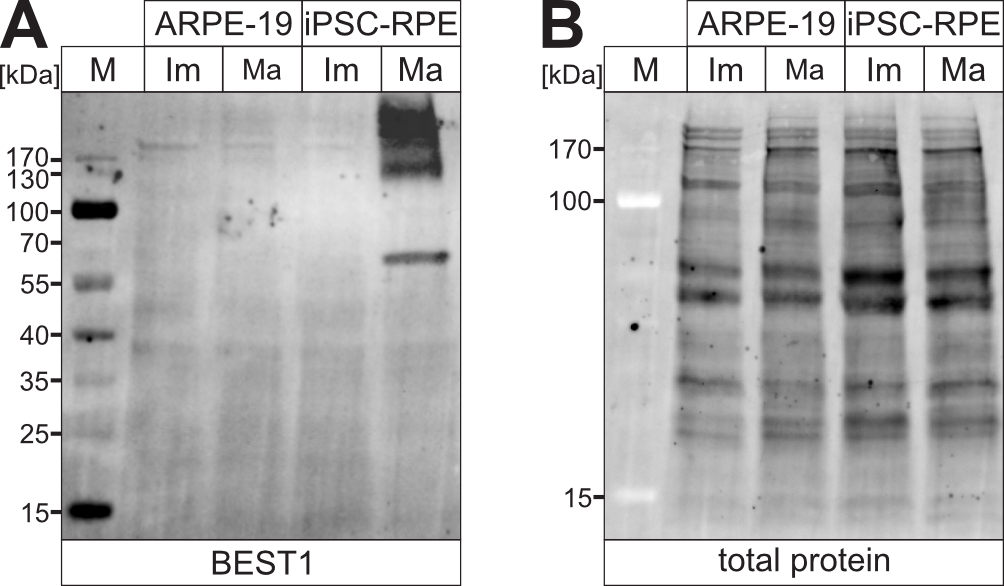

### Figure 1 - figure supplement 3

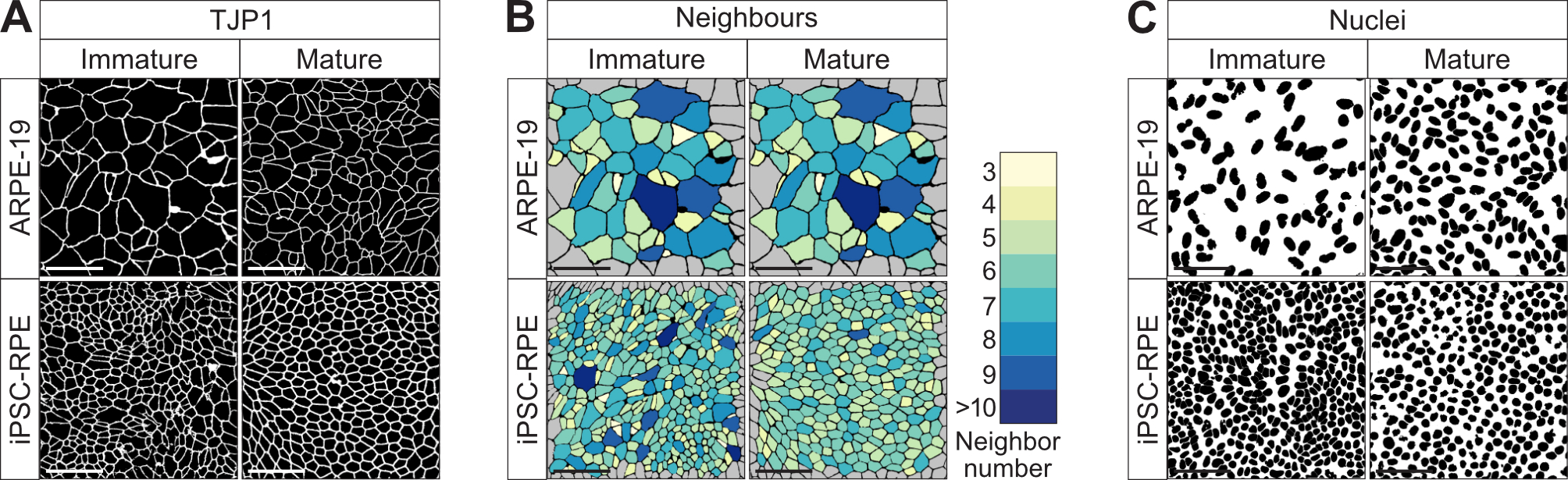

### Figure 3 - figure supplement 1

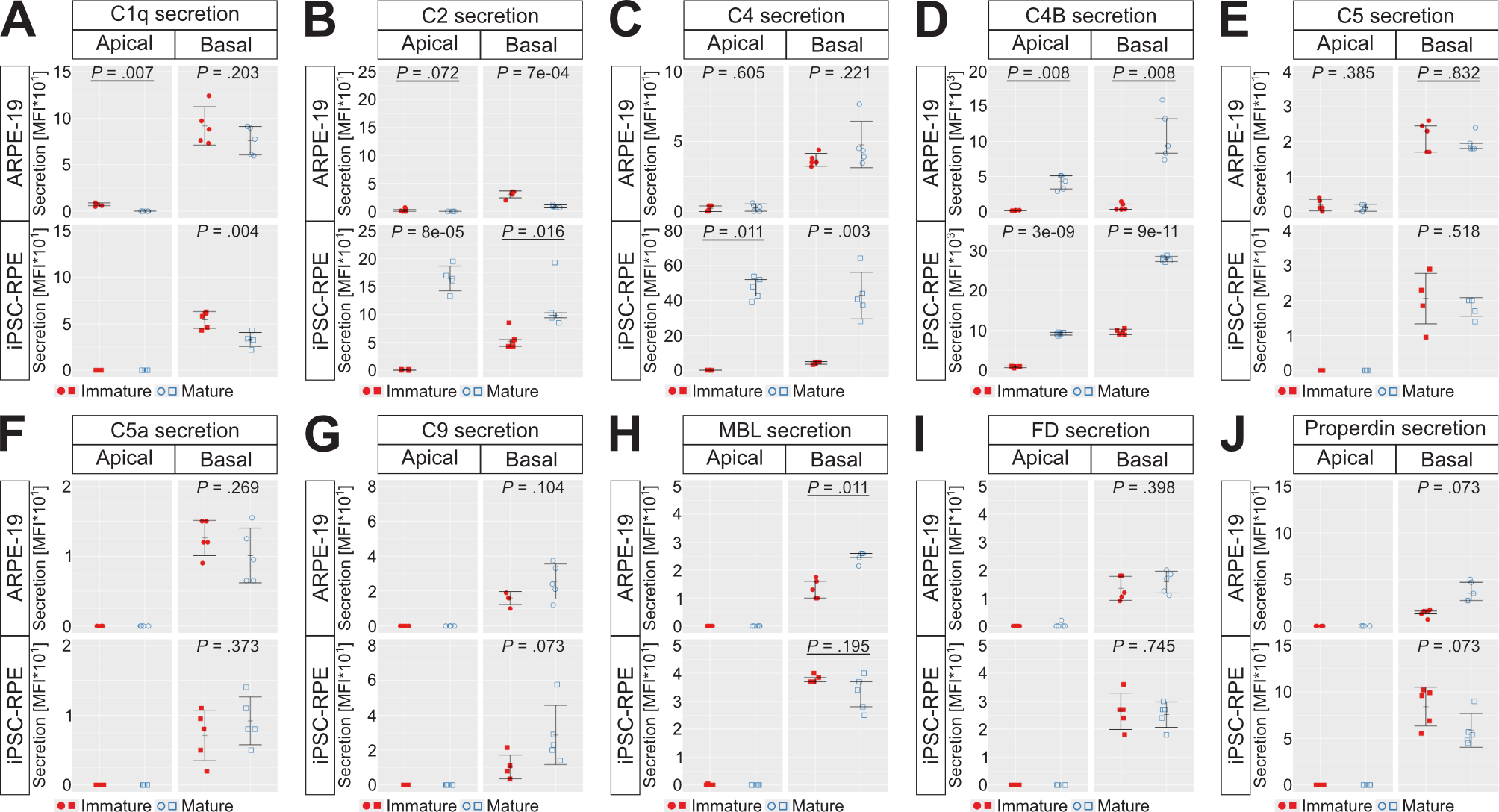

### Figure 3 - figure supplement 2

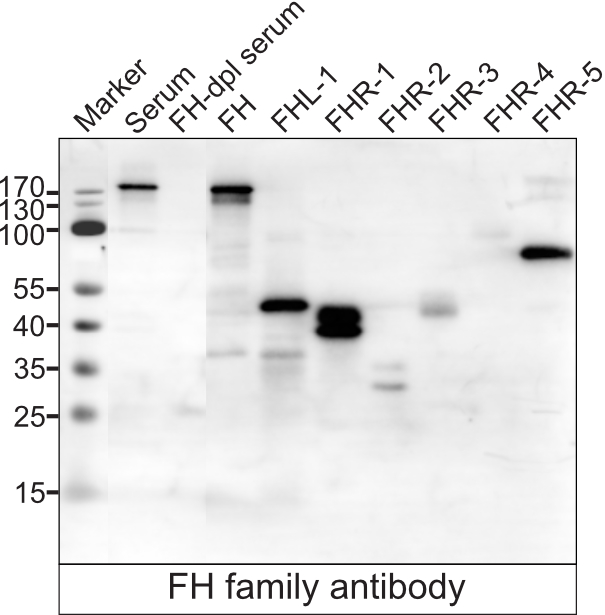

### Figure 3 - figure supplement 3

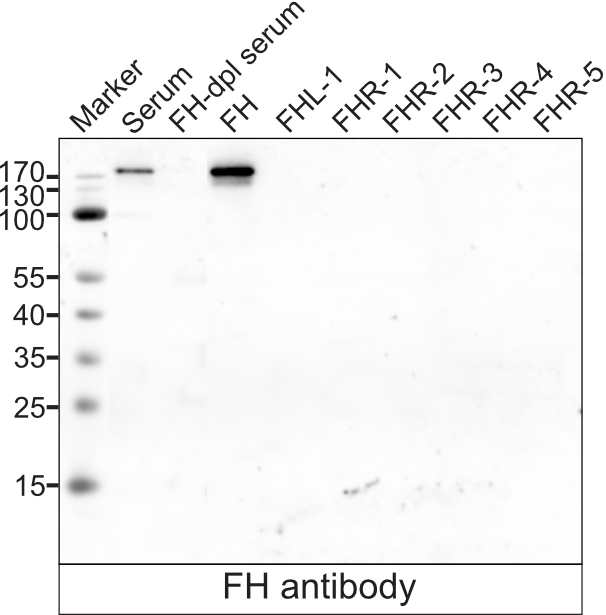

### Figure 3 - figure supplement 4

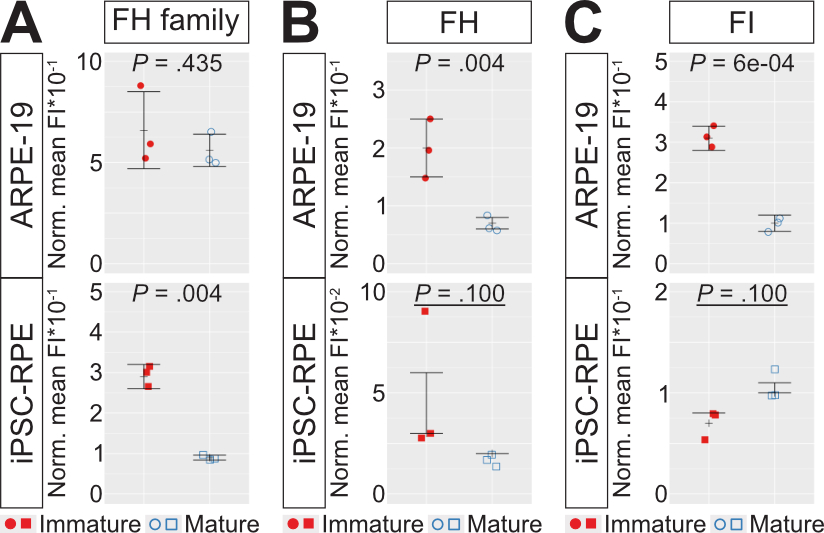

### Figure 4 - figure supplement 1

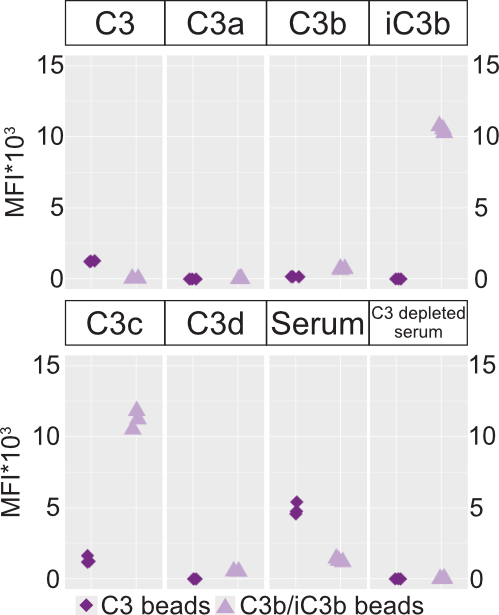

### Figure 5 - figure supplement 1

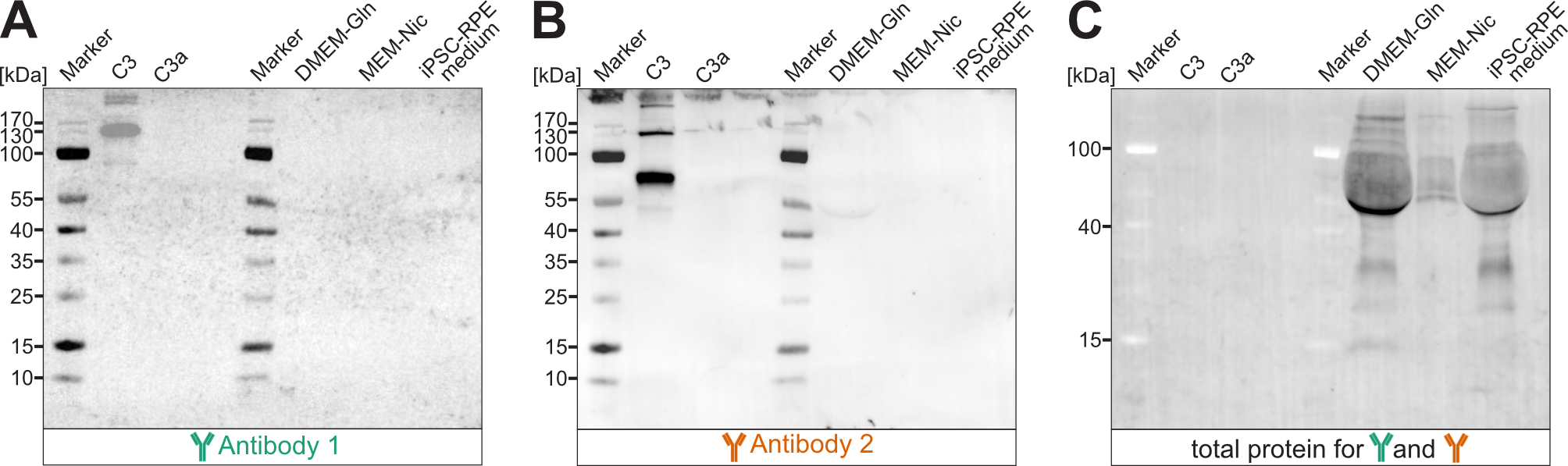

### Figure 5 - figure supplement 2

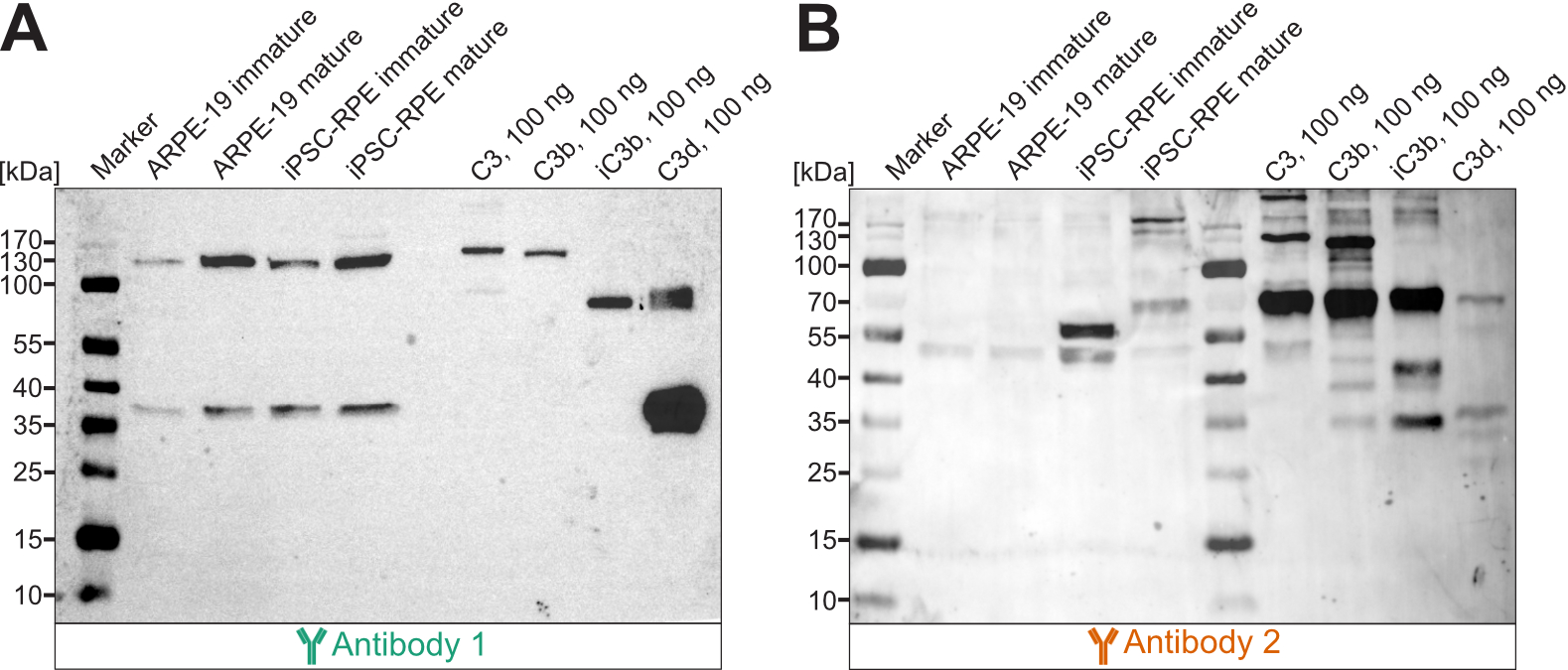

### Figure 5 - figure supplement 3

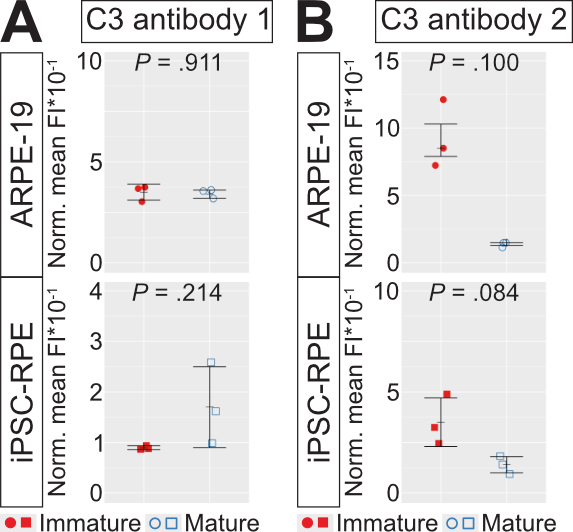

### Figure 5 - figure supplement 4

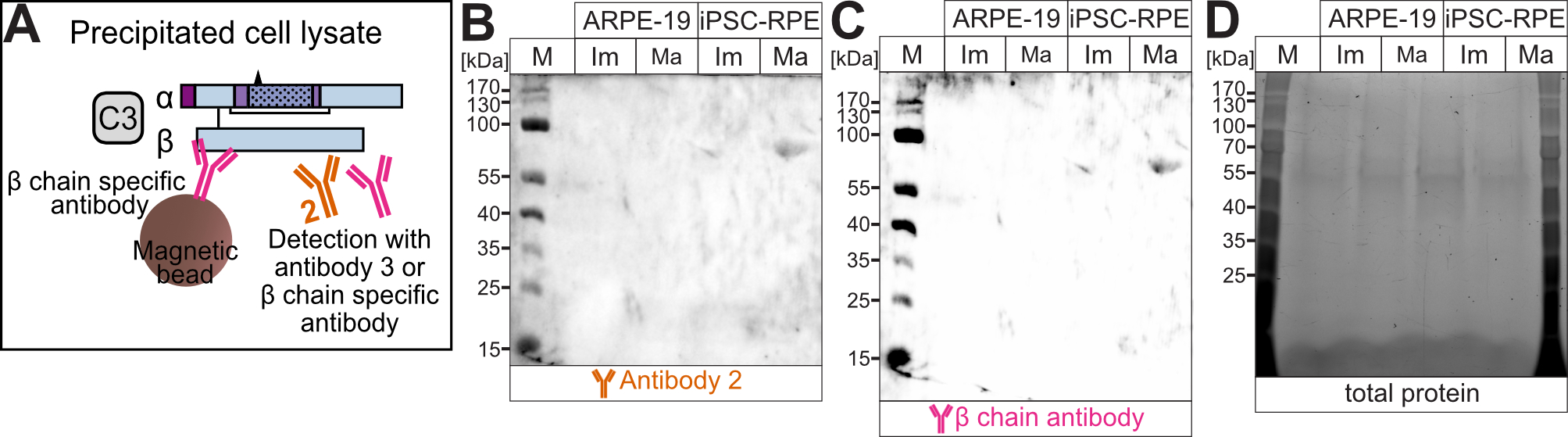

### Figure 6 - figure supplement 1

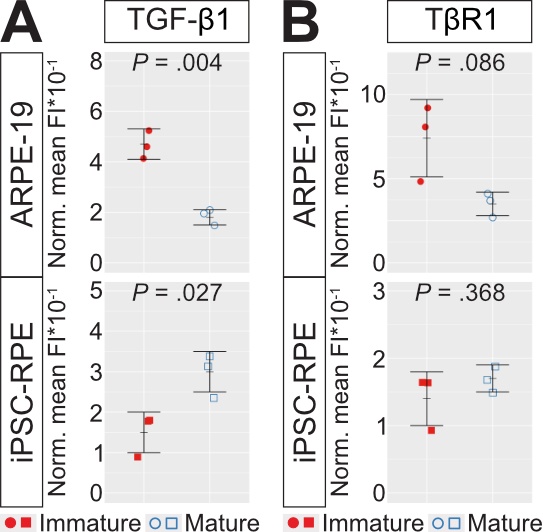

### Figure 7 - figure supplement 1

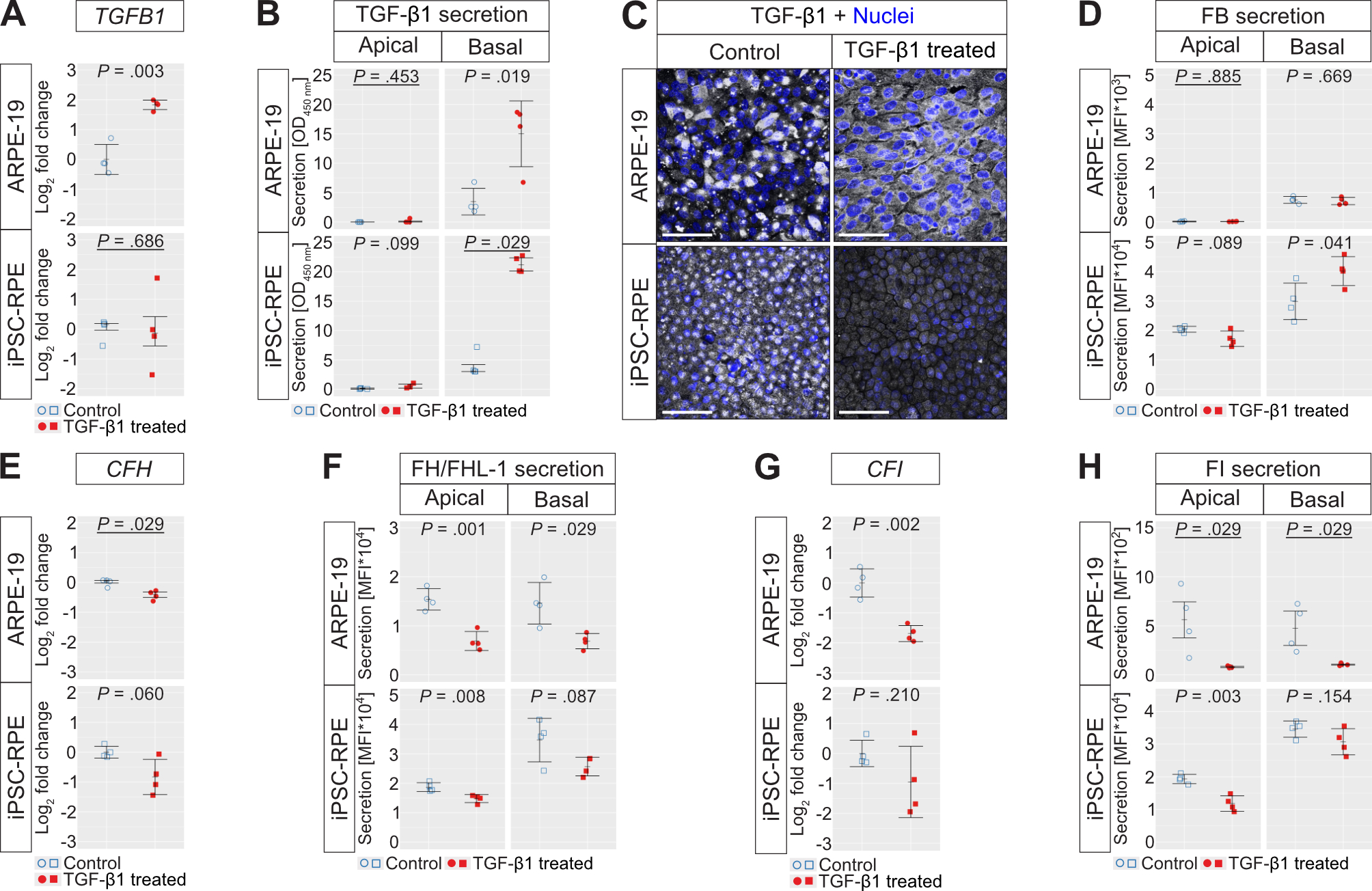

### Figure 7 - figure supplement 2

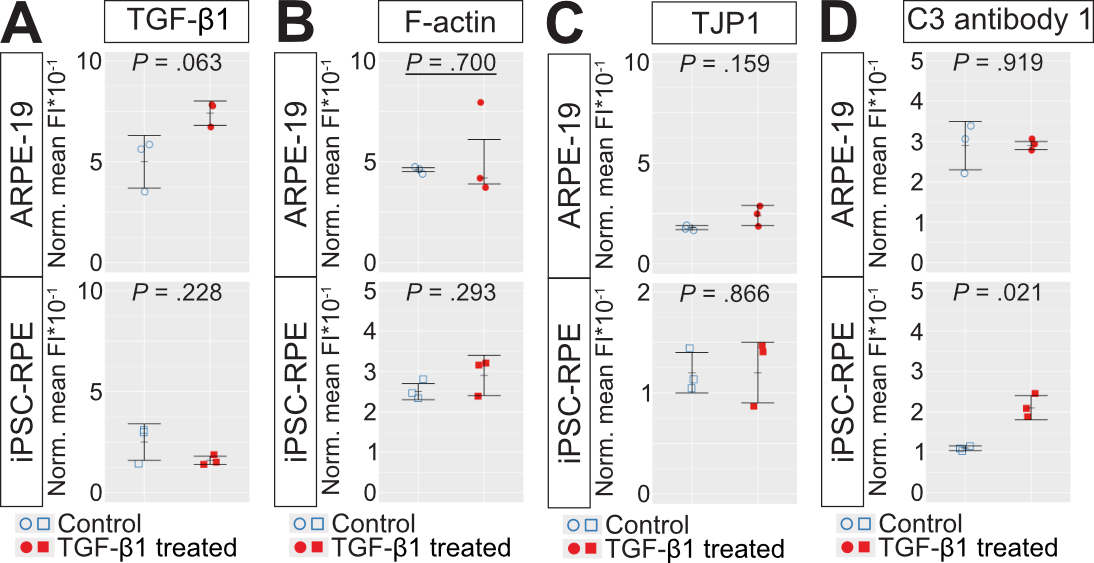
